# A cohesin traffic pattern genetically linked to gene regulation

**DOI:** 10.1101/2021.07.29.454218

**Authors:** Anne-Laure Valton, Sergey V. Venev, Barbara Mair, Eraj Khokhar, Amy H. Y. Tong, Matej Usaj, Katherine S. K. Chan, Athma A. Pai, Jason Moffat, Job Dekker

## Abstract

Cohesin-mediated loop extrusion folds interphase chromosomes at the ten to hundreds kilobases scale. This process produces structural features such as loops and topologically associating domains. We identify three types of *cis*-elements that define the chromatin folding landscape generated by loop extrusion. First, CTCF sites form boundaries by stalling extruding cohesin, as shown before. Second, transcription termination sites form boundaries by acting as cohesin unloading sites. RNA polymerase II contributes to boundary formation at transcription termination sites. Third, transcription start sites form boundaries that are mostly independent of cohesin, but are sites where cohesin can pause. Together with cohesin loading at enhancers, and possibly other *cis*-elements, these loci create a dynamic pattern of cohesin traffic along the genome that guides enhancer-promoter interactions. Disturbing this traffic pattern, by removing CTCF barriers, renders cells sensitive to knock-out of genes involved in transcription initiation, such as the SAGA and TFIID complexes, and RNA processing such DEAD-Box RNA helicases. In the absence of CTCF, several of these factors fail to be efficiently recruited to active promoters. We propose that the complex pattern of cohesin movement along chromatin contributes to appropriate promoter-enhancer interactions and localization of transcription and RNA processing factors to active genes.

**HIGHLIGHTS:**

- At least three types of chromatin boundaries regulate a cohesin traffic pattern.
- The cohesin traffic pattern guides enhancer-promoter interactions.
- Removing CTCF renders cells sensitive to deletion of RNA processing and gene regulation genes.
- Depleting CTCF affects localization of RNA processing and gene regulatory proteins.

## INTRODUCTION

Two main mechanisms govern folding of interphase chromosomes. First, at the megabase level, active and inactive chromatin domains spatially segregate to form A- and B-compartments respectively (Lieberman-Aiden et al., 2009; Simonis et al., 2006). This process is thought to be driven by the biophysical process of phase separation, in which chromatin domains of similar state selectively attract each other (Di Pierro et al., 2016; Erdel and Rippe, 2018; Falk et al., 2019; Hildebrand and Dekker, 2020; Jost et al., 2014; MacPherson et al., 2018; Nuebler et al., 2018). Second, at the scale of tens to hundreds of kilobases, the genome folds into Topologically Associating Domains (TADs) or loop domains, and locus-specific chromatin loops (Dixon et al., 2012; Nora et al., 2012; Rao et al., 2014; Sexton et al., 2012). These structures appear to be formed by a loop extrusion mechanism mediated by cohesin complexes (Alipour and Marko, 2012; Fudenberg et al., 2016; Sanborn et al., 2015). In this model, the cohesin complex dynamically extrudes loops and accumulates at sites where it gets blocked, most prominently at CTCF sites during interphase (Nora et al., 2017; Nuebler et al., 2018; Wutz et al., 2017). Cohesin also accumulates at other sites, e.g., active promoters and, under some conditions, near 3’ ends of genes and sites of convergent transcription (Busslinger et al., 2017). As a result, chromatin loops are often observed between promoters and distal CTCF sites (Kubo et al., 2021; Sanyal et al., 2012; Tolhuis et al., 2002).

Loop extrusion can explain a variety of features observed in Hi-C chromatin interaction maps (Fudenberg et al., 2017). First, focal enrichment of interactions (“dots” in Hi-C maps) between pairs of convergent CTCF sites are readily detected with Hi-C. Second, randomly positioned loops in between CTCF sites result in a general elevation of chromatin interactions in regions bounded by convergent CTCF sites resulting in the formation of TADs (observed as triangles at the diagonal in Hi-C maps). Third, when cohesin cannot extrude past boundary elements such as CTCF sites, chromatin interactions across boundary elements are relatively depleted (referred to as “insulation”, observed as depletion of Hi-C interaction between adjacent domains). All these features disappear when CTCF or cohesin is depleted (Nora et al., 2017; Rao et al., 2017; Wutz et al., 2017) or CTCF-sites are deleted (Hnisz et al., 2016; Narendra et al., 2015; Sanborn et al., 2015; Splinter et al., 2006; de Wit et al., 2015), while CTCF-CTCF loops are enhanced when the cohesin unloader WAPL is depleted (Haarhuis et al., 2017).

The molecular mechanism of loop extrusion is not known in detail, and many open questions remain. For instance, it is not known how and where cohesin is recruited to chromatin, how the complex actively extrudes loops, whether it occurs uni-directionally or bi-directionally, and what constitutes the exact subunit composition of extruding cohesin complexes on chromatin. In addition, it is not known whether other types of *cis*-elements besides CTCF-bound sites contribute to cohesin dynamics and stalling along chromosomes. The mechanism by which CTCF blocks cohesin in a CTCF-site orientation dependent manner is better understood. The N-terminus of CTCF directly binds to the SCC1/RAD21 and SA1/STAG subunits of cohesin (Li et al., 2020). In that configuration, cohesin is stalled and protected from WAPL-mediated unloading. Finally, 3’ ends of genes, and locations of convergent gene expression may be positions where cohesin is unloaded: when CTCF and WAPL are depleted, cohesin accumulates at these positions (Busslinger et al., 2017).

Roles for extrusion-mediated chromatin structural features in appropriate gene regulation are currently intensely debated. TADs are thought to regulate gene expression by allowing enhancer-promoter interactions within the domain, while disfavoring such interactions across their boundaries (Flavahan et al., 2016; Franke et al., 2016; Hnisz et al., 2016; Lupiáñez et al., 2015, 2016; Valton and Dekker, 2016). TAD boundaries tend to have multiple CTCF binding motifs and a recent study shows that CTCF sites at TAD boundaries are under evolutionary constraint highlighting their functional importance (Kentepozidou et al., 2020). Genetic re-arrangements that alter the order of *cis*-elements such as CTCF sites, and therefore TAD positions, can lead to altered gene expression in ways that are consistent with the model that TADs constrain promoter-enhancer interactions (Lupiáñez et al., 2015). In some cases, chromatin boundary and CTCF site deletion can expose a new enhancer to an oncogene, leading to overexpression of the gene and potentially contribute to tumorigenesis (Flavahan et al., 2016; Hnisz et al., 2016). However, acute global depletion of CTCF or the cohesin subunit RAD21 leads to changes in gene expression for only a small number of genes, despite genome-wide loss of TADs and CTCF-CTCF loops (CTCF depletion, (Nora et al., 2017)), or loss of all extrusion features including enhancer-promoter interactions (RAD21 depletion, (Rao et al., 2017)). Several models have been put forward to explain these apparently contradicting results, e.g., stating that looping interactions are not continuously needed to maintain promoter activity (Xiao et al., 2021a; Zuin et al., 2021), leading to loss of correlation between enhancer-promoter interactions and gene transcription within hours. Despite these theoretical advances, functions of loop extrusion, and cohesin blocking at specific sites, remain poorly understood.

Here, we analyzed Hi-C data obtained with cells in which CTCF, RAD21 or RNA polymerase II can be acutely depleted to describe the intricate local folding of chromosomes and the roles of different types of *cis*-elements in guiding cohesin-mediated loop extrusion. A complex picture of cohesin traffic along chromosomes emerges. To uncover functional roles for this intricate chromosome organization, we performed a genome-wide CRISPR screen in cells with altered cohesin traffic patterns following CTCF depletion. We identified genes involved in transcription initiation and RNA processing. A modified cohesin traffic pattern resulted in an incorrect localization of these transcription factors and slightly affected gene expression.

## RESULTS

### CTCF is necessary for establishing most of the TADs but not compartments

We reasoned that by acutely depleting the CTCF protein from cells, we would be able to study functional roles of chromatin loops formed between CTCF sites and potentially reveal other elements that could block the loop extrusion machinery. To that end, we engineered a HAP1-derived human cell line in which CTCF could efficiently be removed using an auxin-inducible degron system (Figure S1A). We found that adding a C-terminal auxin-inducible degron domain (AID) to CTCF led to inefficient CTCF depletion upon auxin treatment, as inferred from the persistence of looping interactions between CTCF sites (data now shown). One possibility is that HAP1 cells express CTCF isoforms that lack the full C-terminus and therefore the AID tag. To overcome this problem, we integrated a full-length CTCF cDNA tagged with AID at the N-terminus and with AID and eGFP at the C-terminus (Figure S1A, see Methods for details). This gene encodes the full length CTCF protein fused to two AIDs tags and one eGFP tag. We then knocked out the endogenous CTCF gene to obtain a cell line that we refer to as HAP1-CTCFdegron. Finally, we introduced the TIR1 gene to produce the HAP1-CTCFdegron-TIR1 cell line. The TIR1 protein binds to the auxin/degron complex and promotes its degradation by the proteasome. Thus, addition of auxin to HAP1-CTCFdegron-TIR1 cells resulted in efficient depletion of CTCF as assessed by western blotting (Figure S1B). FACS analysis of eGFP levels showed depletion of CTCF after 4 hours of auxin treatment (Figure S1C). The cell cycle profile was not altered in CTCF depleted cells (Figure S1D). We noted that even without addition of auxin the CTCF protein level was reduced compared to the level observed in HAP1-CTCFdegron cells that do not contain TIR1 (Figure S1B-C). We conclude that CTCF can be effectively degraded in HAP1-CTCFdegron-TIR1 cells in the presence of auxin, and that HAP1-CTCFdegron-TIR1 cells express reduced levels of CTCF even in the absence of auxin.

We performed Hi-C on HAP1-CTCFdegron and HAP1-CTCFdegron-TIR1 cells grown in the absence or presence of auxin for 48 hours (Figure S1H-K). In the absence of auxin, domain boundaries at CTCF sites and CTCF-CTCF loop formation are reduced in HAP1-CTCFdegron-TIR1 cells compared to HAP1-CTCFdegron cells (Figure S1K). This can be explained by reduced CTCF expression in HAP1-CTCFdegron-TIR1 cells. In the presence of auxin, depletion of CTCF resulted in near complete loss of looping interactions between CTCF sites, loss of enrichment of interactions within topologically associating domains and loss of insulation at domain boundaries. This is consistent with previous observations in CTCF degron cell lines (Nora et al., 2017; Wutz et al., 2017). Compartmentalization was only slightly affected. These changes were not observed in HAP1-CTCFdegron cells treated with auxin, indicating that these slight compartmentalization effects were significant (Figure S1I-J). Finally, we analyzed the relationship between Hi-C interaction frequency as a function of genomic distance between loci which allows us to infer the average loop size (Gassler et al., 2017). Interestingly, we found that the average size of loops, generated by extruding cohesins, increased progressively when CTCF levels were reduced (in HAP1-CTCFdegron-TIR1 cells in the absence of auxin) or entirely depleted (HAP1-CTCFdegron-TIR1 in the presence of auxin). This is expected when CTCF-mediated blocking of loop extrusion is reduced or abolished.

To study if CTCF was needed to establish TADs in the G1 phase of the cell cycle, we cultured HAP1-CTCFdegron-TIR1 cells with or without auxin for 12 hours to deplete CTCF. Under these conditions and for this time frame (12 hours), cells continued to go through the cell cycle. We then used FACS to isolate G1 cells that either lacked CTCF (GFP negative, in the presence of auxin) or contained CTCF (GFP positive, in the absence of auxin; Figure S1E-F). Given the ∼24 hour doubling time of HAP1-CTCFdegron-TIR1 cells, these cells must have entered G1 in the absence or presence of CTCF respectively. Hi-C analysis showed that establishment of compartmentalization was only slightly affected by the absence of CTCF. In contrast, TADs were mostly not established, which was observed in Hi-C interaction maps by increased interactions across TAD boundaries (Figure 1A-B). Looping interactions between CTCF sites, readily observed in Hi-C maps obtained with control cells, were greatly reduced in cells treated with auxin (Figure 1C). Finally, we observed a slight shift towards longer extruded loops in CTCF-depleted cells, consistent with reduced blocking of cohesin (Figure S1G). We conclude that CTCF is required for re-establishing TADs and CTCF-CTCF loops as cells exit mitosis and enter G1.

**Figure 1.**
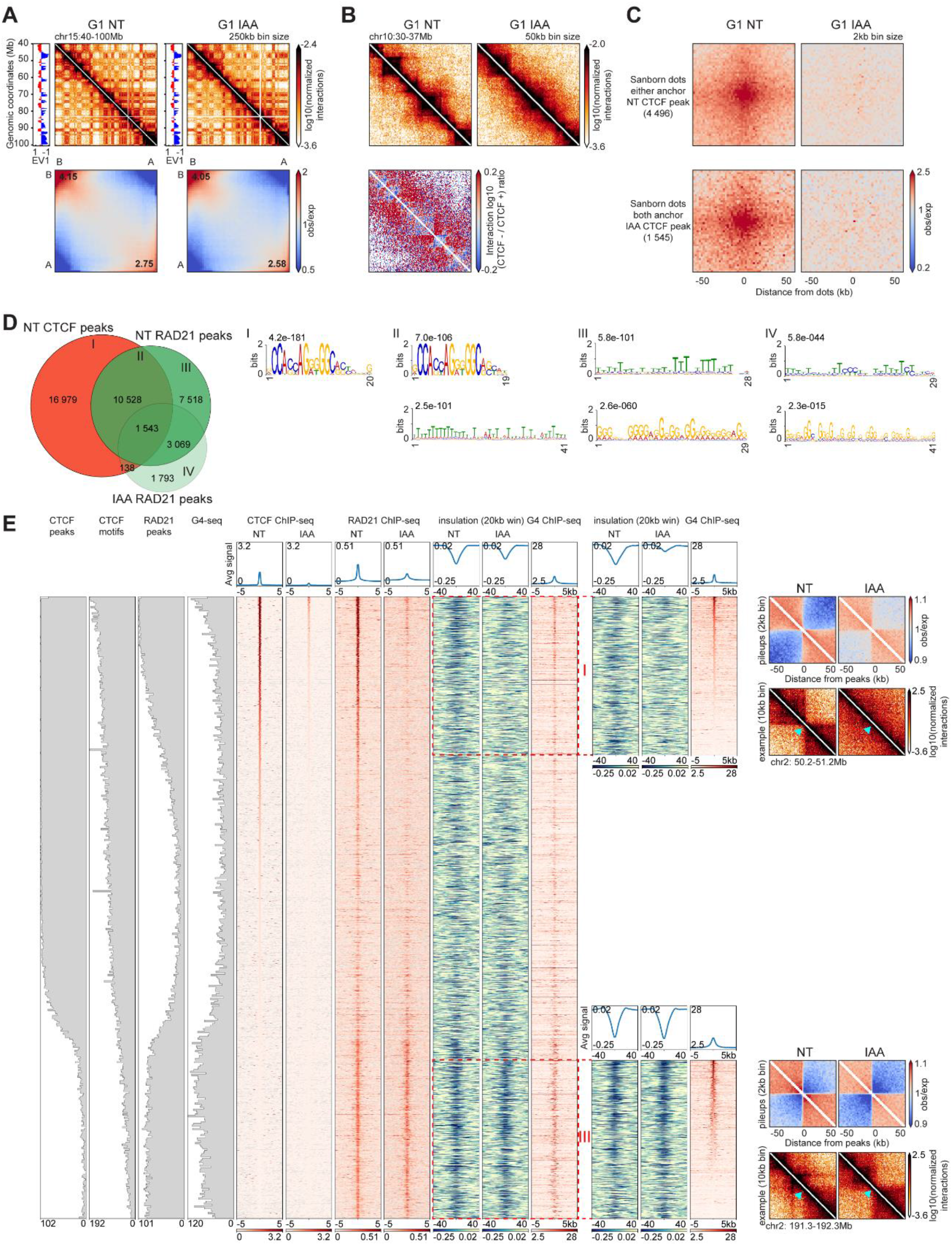
CTCF is necessary for establishing most of TADs and a subset of chromatin boundaries are CTCF independent and contain G-quadruplexes. (A) Hi-C contact heatmaps at 250kb resolution with the corresponding track of the first Eigenvector (EV1) for a 60Mb region on chromosome 15 for HAP1-CTCFdegron-TIR1 cells sorted in G1 phase in absence of auxin (G1 NT) or in presence of auxin (G1 IAA) (top). Genome-wide saddle plots of Hi-C data binned at 100kb resolution for NT and IAA treated cells. The numbers in the corners of the saddle plots indicate compartment strengths for A and B compartments (bottom). (B) Hi-C contact heatmaps at 50kb resolution for a 7Mb region on chromosome 10 for HAP1-CTCFdegron-TIR1 cells sorted in G1 phase in absence of auxin (G1 NT) or in presence of auxin (G1 IAA) with the resulting differential contact heatmap showing loss of insulation at boundaries (red) and loss of interactions within TADs (blue) after CTCF depletion. (C) Pileups of dots characterized in HAP1 cells (Sanborn et al., 2015) that have a CTCF peak in either anchor in the Non-Treated sample (NT) (4 496 dots) and those that have a CTCF peak in both anchors in the auxin sample (IAA) (1 545 dots) for HAP1-CTCFdegron-TIR1 cells sorted in G1. The dots were aggregated at the center of a 100kb window at 2kb resolution. (D) Venn diagram showing the overlap between CTCF peaks in the absence of auxin (NT), RAD21 peaks in the absence and presence of auxin (NT and IAA) (left). MEME motif search for NT CTCF-only peaks (I), overlap between NT CTCF peaks and NT RAD21 peaks (II), NT RAD21 peaks that do not overlap with CTCF (III), IAA RAD21-only peaks (IV) (right). (E) Stackups of the union list of all RAD21 and CTCF peaks, sorted on the Non-Treated (NT) CTCF ChIP-seq signal for the. CTCF and RAD21 ChIP-seq and calculated insulation in the absence and presence of auxin (NT and IAA) were plotted along with the published K562 G4 ChIP-seq. The distribution of CTCF peaks, CTCF motifs, RAD21 peaks and G4-seq signals were plotted along the pileups. Two groups of loci were highlighted with red dashed rectangles: the CTCF dependent insulation (I, top) and the CTCF independent insulation (III, bottom), sorted on G4 ChIP-seq signal. For each category, an interaction pileup, in absence and presence of auxin (NT and IAA), aggregated in a 100kb window at 2kb resolution was plotted along with a representative example of a boundary for each set. The light blue arrow shows the boundary location. See Figure S1

### A subset of domain boundaries is CTCF independent and contain G-quadruplexes

To assess how the genomic positioning of the cohesin loop extrusion complex is affected after CTCF depletion, we performed Chromatin ImmunoPrecipitation sequencing (ChIP-seq; two biological replicates) for CTCF and the cohesin subunit RAD21 in HAP1-CTCFdegron-TIR1 cells in the presence of CTCF (control cells) or absence of CTCF (auxin treated cells). We identified 29,188 CTCF peaks and 22,658 RAD21 peaks in the presence of CTCF (Figure 1D, see Methods for details). Only around 40% of the CTCF peaks overlapped with RAD21 peaks, showing that not all CTCF sites are associated with cohesin. Moreover, approximately 50% of the RAD21 peaks overlapped with CTCF peaks, suggesting that RAD21 can accumulate at locations devoid of CTCF. We performed MEME motif analysis on the set of loci that only contained CTCF peaks and found, as expected, the CTCF consensus motif (category I). When performing MEME analysis on the set of loci that contain both a CTCF peak and a RAD21 peak (category II) we again found the CTCF consensus motif as well as a weak T-rich motif. The RAD21-only peaks in cells with CTCF were enriched for two motifs and these differed from the CTCF motif: a weak T-rich motif and a G-rich motif (category III). Finally, we found that the set of RAD21 peaks only detected in cells in which CTCF is depleted was also enriched for G-rich and T-rich motifs (category IV) (Figure 1D). These results show that RAD21 can also accumulate at motifs different from the CTCF motif. The G-rich motif we identified shows strong resemblance to G-quadruplexes (G4s). G4s are G-rich sequences with four runs of guanines and at least three guanines per run (G≥3NXG≥3NXG≥3NXG≥3NX) and can form non-canonical DNA structures by forming stacks with Hoogsteen bonds between the opposite guanines. G4s are often found at active promoters (see below).

CTCF-bound sites are well known boundary elements and are able to block cohesin in an orientation dependent manner. Sites that block cohesin-mediated extrusion often form domain boundaries in Hi-C interaction maps. We were interested in determining whether the additional sites we identified that accumulate cohesin (RAD21) but do not bind CTCF were also able to form chromatin domain boundaries. Domain boundary formation can be quantified by the insulation metric, which measures the extent to which long-range chromatin interactions across a boundary are reduced compared to a global average (Crane et al., 2015). We determined insulation at the different types of elements that accumulate RAD21 described above. We created a union list of all RAD21 peaks detected in either the presence or absence of CTCF and all the CTCF peaks detected in CTCF-expressing cells. The final union list contained 39,233 peaks in total (see Methods). We then analyzed CTCF binding, RAD21 binding and insulation for these sites in CTCF expressing and CTCF-depleted HAP1-CTCFdegron-Tir1 cells and ranked these sites by the level of CTCF binding in CTCF expressing cells. We also assessed G-quadruplex locations by plotting published data obtained with G4 ChIP-seq in K562 cells and G4-seq for all loci included in the union list (Chambers et al., 2015; Mao et al., 2018). We identified three major groups of elements that differ in CTCF and RAD21 binding. The first group of sites bind CTCF at high levels with most sites being included in the set of significantly enriched CTCF peaks. These sites also contained high levels of RAD21, and the majority of these were included in the set of significant RAD21 peaks. These sites displayed strong insulation, indicating they form domain boundaries. A small fraction of these sites contained G4 sequences. Sites in this group lose RAD21 binding and insulation in cells depleted for CTCF. The second group of sites also bound CTCF at high levels and were often included in the set of significant CTCF peaks. However, RAD21 was not found enriched at these sites, and these sites did not display insulation, indicating they were not at chromatin domain boundaries. Few of the sites in this group contained G4 sequences. The third group of sites did not show enriched CTCF binding but displayed relatively high levels of RAD21 binding and insulation in control cells. Many of these sites contained G4 sequences and displayed relatively strong G4 ChIP-seq signals. In cells that were depleted for CTCF, these sites continued to accumulate RAD21 and to display insulation. Importantly, the large majority of these CTCF-independent and G4-containing boundaries did not overlap with compartment boundaries (Figure 1E, Figure S1L), suggesting they were *bona fide* cohesin-bound chromatin domain boundaries. We concluded that RAD21 can accumulate at at least two different types of locations and form domain boundaries, likely through blocked loop extrusion: 1) at CTCF sites, where RAD21 accumulation is dependent on CTCF; and 2) at a set of sites that are devoid of CTCF and that contain G4 sequences. At these sites, RAD21 binding and insulation are CTCF independent.

### Cohesin accumulates at G4-containing active promoters

G4s are often found at telomeres and promoters and have been identified as transcription regulators (Blackburn, 1991; Henderson et al., 1987; Huppert and Balasubramanian, 2007; Maizels and Gray, 2013). To assess the genomic and chromatin context of G4-containing regions that could contribute to the observed RAD21 accumulation and chromatin insulation, we took the lists of CTCF ChIP-seq peaks and G4 ChIP-seq and created three subsets of sites: CTCF-only, CTCF + G4, and G4-only peaks. We then analyzed CTCF binding, RAD21 binding and insulation at these peaks in cells with CTCF or in cells depleted for CTCF. We also analyzed G-quadruplex locations, RNA polymerase II (RNA polII) binding, and coding gene locations from published datasets (Chambers et al., 2015; Fong et al., 2015; Mao et al., 2018). First, CTCF-only sites accumulated RAD21 near their center, and this accumulation disappeared when CTCF was depleted. Insulation at these sites was also CTCF dependent. Second, the strongest binding of CTCF and RAD21 was observed at sites that overlapped both CTCF peaks and G4 peaks. RAD21 binding and insulation was only slightly reduced at these sites when CTCF was depleted. Third, the set of loci that only contained G4 peaks also accumulated RAD21, consistent with the results shown in Figure 1E. RAD21 enrichment and insulation at G4-only sites was CTCF-independent. We noticed a slight increase of RAD21 accumulation at G4-only peaks in the cells depleted for CTCF. Possibly, in the absence of CTCF, RAD21 is no longer blocked at CTCF sites and now more frequently accumulates at sites containing G4 sequences. This agrees with previous observations in mouse that cohesin accumulates more at promoters (i.e., G4 containing sequences) in the absence of CTCF (Busslinger et al., 2017). Importantly, we find that most G4-only sites, and a subset of the CTCF+G4 sites, were located at annotated promoters/Transcription Start Site (TSS) of genes that showed RNA polII binding in HAP1 cells (Figure 2A-B). The large majority of the insulating G4-containing sites did not overlap with compartment boundaries (Figure S2D), ruling out the possibility that insulation at these sites is the result of compartmentalization.

**Figure 2.**
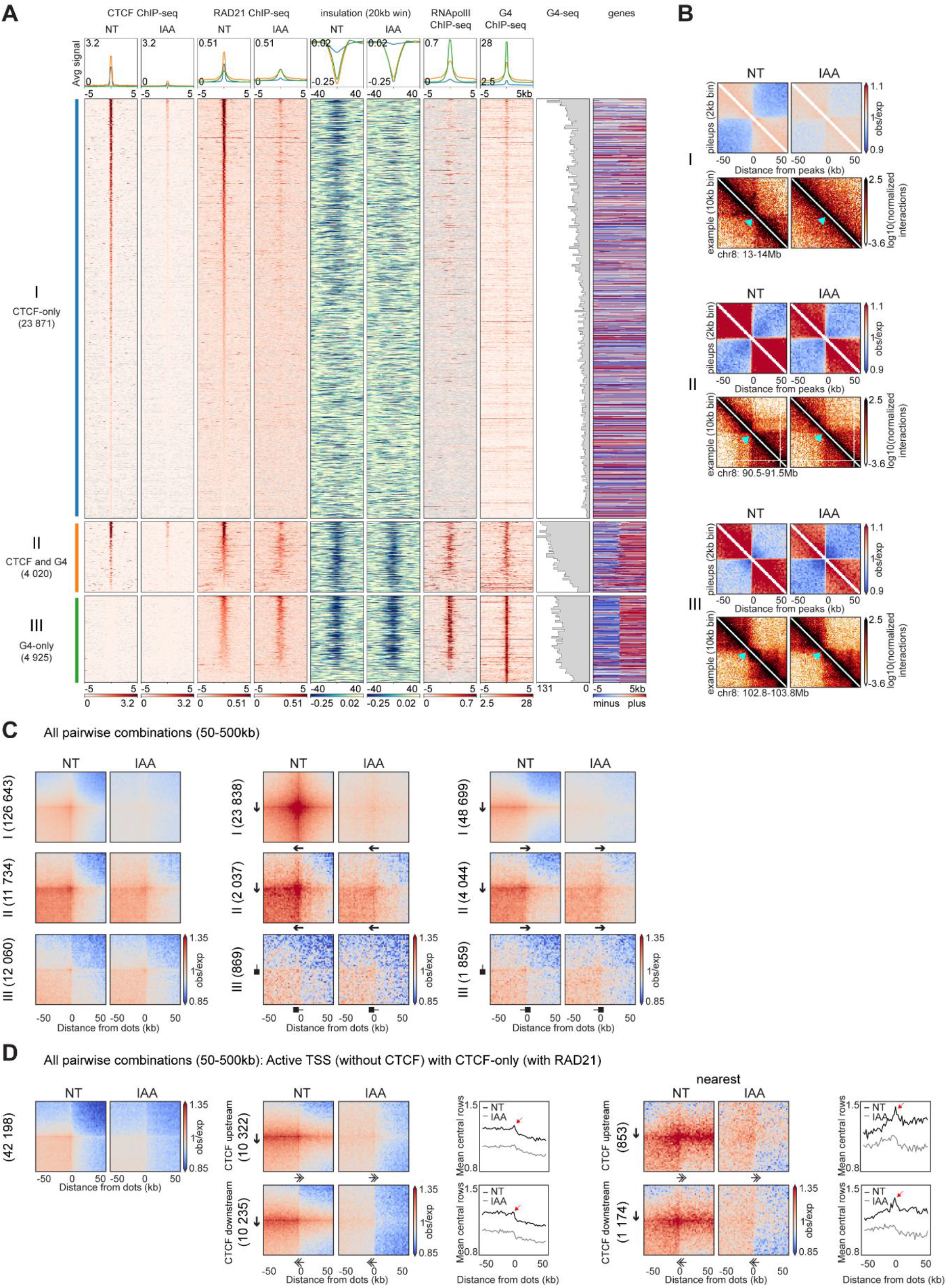
Cohesin accumulates at G4-containing active promoters/TSSs that are chromatin boundaries. (A) Stackups for three categories: the CTCF-only peaks (I, blue), the CTCF and G4 peaks (II, orange) and the G4-only peaks (III, green), all sorted on Non-Treated (NT) RAD21 ChIP-seq signal. The CTCF and RAD21 ChIP-seq signals were plotted, with the insulation in the HAP1-CTCFdegron-TIR1 in the absence or presence of auxin (NT and IAA). The published HAP1 RNA polII ChIP-seq and the K562 G4 ChIP-seq signals were plotted. The distribution of G4-seq signals was plotted along the stackup and depiction of annotated genes was added on the last column. Genes on the forward strand are represented in red (plus) and genes on the reverse strand are represented in blue (minus), grey color corresponds to loci without annotated transcripts. (B) Interaction pileups for each category represented in (A) (I, CTCF-only, II, CTCF and G4, III, G4-only) were plotted along with a representative example of a boundary for each set, both in absence and presence of auxin (NT and IAA), aggregated in a 100kb window at 2kb resolution. The light blue arrow shows the boundary location. (C) Dot pileup aggregation plots, for all the pairwise combinations of each category represented in (A) (I, CTCF-only, II, CTCF and G4, III, G4-only) separated by 50-500kb, in absence and presence of auxin (NT and IAA), for a 100kb window at 2kb resolution. The black arrows represent the CTCF motif and the direction of the arrow, the motif orientation. The black squares represent the G4 and the location of the black square, the G4 orientation (right, plus strand). (D) Dot pileup aggregation plots, for all the pairwise combinations of active TSS and CTCF-only peaks (with RAD21) separated by 50-500kb, in absence and presence of auxin (NT and IAA), for a 100kb window at 2kb resolution. All the pairwise interactions are plotted (left). CTCF (upstream)-TSS pairwise interactions and CTCF (downstream)-TSS pairwise interactions are plotted. A quantification of the aggregation pileups is plotted (mean of the 5 central bins at the CTCF site). The red arrow represents the peak of interactions between CTCF site and TSS (middle). CTCF (upstream)-TSS pairwise interactions and CTCF (downstream)-TSS pairwise interactions are plotted without any CTCF peaks or TSS in between them. A quantification of the aggregation pileups is plotted (mean of the 5 central bins at the CTCF site). The red arrow represents the peak of interactions between CTCF site and TSS (nearest, right). The black arrows represent the CTCF motif and the direction of the arrow, the motif orientation. The double arrows represent the TSS and the direction of the arrow, the TSS orientation. See Figure S2

We found that formation of G4 sequences (as determined by G4 ChIP-seq) was highly correlated with the activity of a TSS, as determined by RNA polII ChIP-Seq, H3K4me3 ChIP-seq, and RNA-seq (Figure S2A-B). Therefore, we cannot at this time separate, roles of these features in RAD21 binding and insulation. We conclude that cohesin accumulates in a CTCF independent manner at active promoters containing G4 sequences.

Sites that block cohesin-mediated loop extrusion, such as CTCF-bound sites, can form chromatin domain boundaries as well as loops with other such sites (Guo et al., 2015; Rao et al., 2014; Vietri Rudan et al., 2015; de Wit et al., 2015). Thus, we asked whether G4-containing sites can form domain boundaries and/or engage in long-range looping interactions. We aggregated all pairwise combinations between different sets of sites separated by 50-500kb. We confirmed that CTCF-only peaks were able to form loops with nearby CTCF-only peaks in a CTCF-dependent manner. Such looping interactions occur most frequently when CTCF sites are in a convergent orientation, as expected (Figure 2C, Figure S2C, set I)). Looping interactions between sites containing both G4 and CTCF peaks (set II) were less dependent on CTCF, but also were most frequent when CTCF sites are in a convergent orientation. Finally, sites in set III that contained G4 peaks and active promoters form CTCF-independent domain boundaries but no obvious focal enrichment (i.e. “dots”) of TSS-TSS interactions were observed (Figure 2C, Figure S2C).

We then quantified interactions between different types of cohesin-bound sites. We found that active TSSs (that lack CTCF binding) frequently interacted with nearby (50-500kb up- or downstream) CTCF-only sites (that contain RAD21) (Figure 2D, Figure S2C). As expected, these interactions were CTCF dependent and were most frequent when the CTCF motif pointed toward the TSS. The orientation of the TSS itself appeared less consequential, although some quantitative differences could be observed. In Hi-C interaction maps, lines of enriched interactions were visible from the distal CTCF sites towards the active TSS. No such lines were detected anchored on TSSs. When we quantified the strength of this enrichment of CTCF-anchored interactions, we observed a peak in interactions centered on the TSS (indicated by arrows in Figure 2D). All these features disappeared when CTCF was depleted. We interpreted these results as follows: cohesin actively extrudes chromatin until it is blocked on one side by CTCF while continuing to extrude on the other side towards a TSS. When it reaches the TSS, extrusion pauses and results in a local enrichment of CTCF-TSS interactions. Cohesin can subsequently occasionally extrude beyond the TSS. In the absence of CTCF, cohesin is still able to extrude chromatin loops and when it reaches a TSS it will pause, producing domain boundaries at TSSs as detected by Hi-C.

### 3’ ends of active genes form cohesin-dependent domain boundaries

Previous studies have shown that in the absence of the cohesin unloading factor WAPL and CTCF, cohesin accumulates at 3’ ends of active genes. No such accumulation is seen in WAPL expressing cells (Busslinger et al., 2017). This indicates that in normal cells, cohesin is efficiently unloaded once it reaches the 3’ end of genes. Unloading of cohesin at 3’ ends of active genes is expected to result in the formation of chromatin domain boundaries because cohesin cannot extrude past the end of the gene. To test this, we calculated local insulation near the transcription termination site (TTS) of active genes that do not contain CTCF-bound sites. We detected local minima in the insulation scores, indicating the formation of domain boundaries. RAD21 ChIP-seq confirmed that cohesin was not accumulating at these sites, presumably because it was unloaded efficiently. We again noted that these domain boundaries at TTSs mostly did not overlap with compartment boundaries (Figure S3A). We noticed that the local insulation minima were less precisely positioned as compared to those located at TSSs and CTCF-bound sites, and their detection required calculating insulation scores using a larger genomic window (100kb instead of 20kb). Insulation at TTSs was unaffected after depletion of CTCF. Strongly insulating TTSs correlated with R-loop peaks and suggested a link between R-loops and insulation at TTS (Figure 3A).

**Figure 3.**
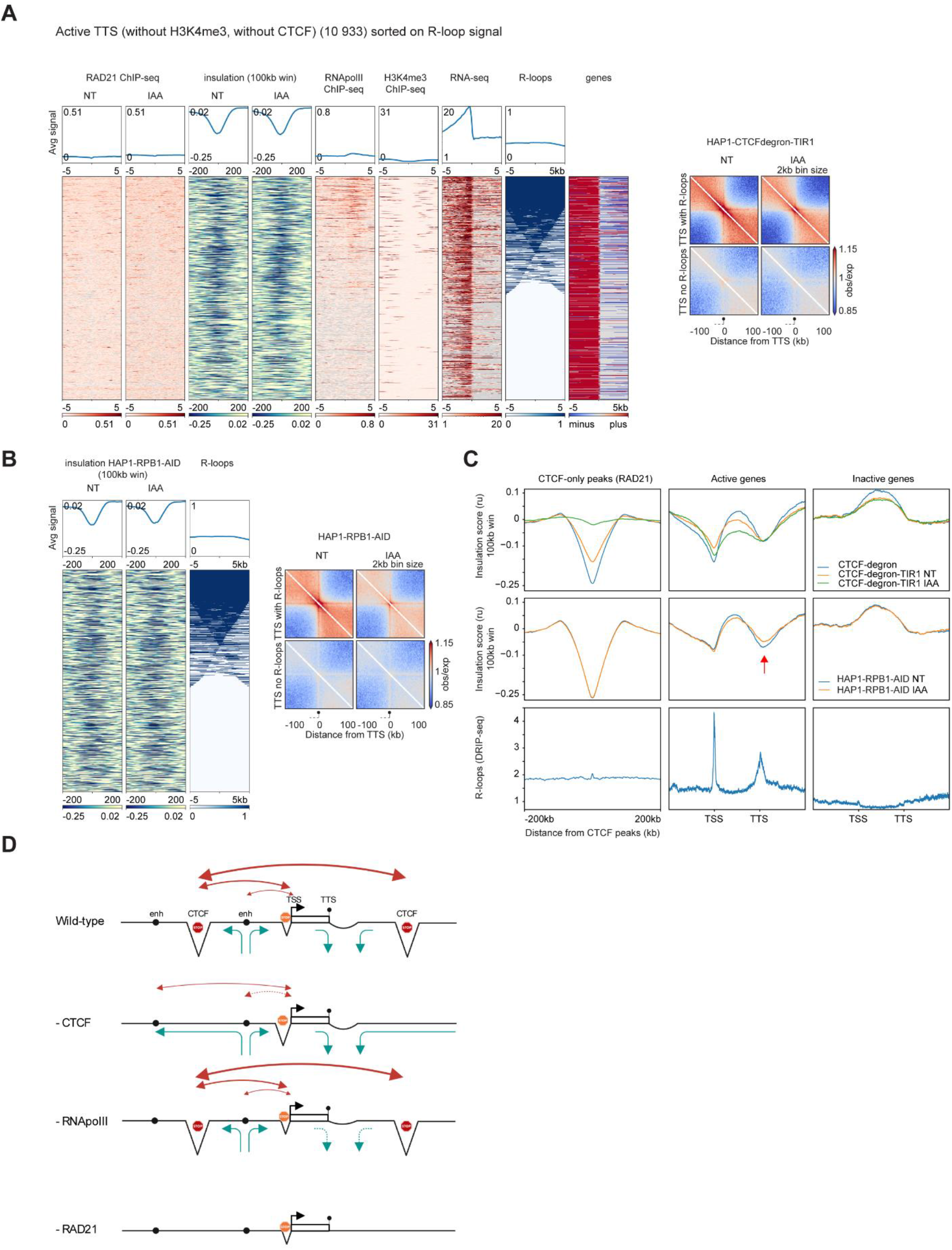
3’ ends of active genes form cohesin-dependent chromatin boundaries that are weakly affected by RNA polII depletion. (A) Stackups for active TTSs (without H3K4me3 active TSS histone mark and without CTCF peaks) sorted on a consensus list of R-loops. RAD21 ChIP-seq and calculated insulation in the absence or presence of auxin (NT and IAA) were plotted along with the published HAP1 RNA polII ChIP-seq, the HAP1 H3K4me3 ChIP-seq, the RNA-seq in HAP1-CTCFdegron-TIR1 in absence of auxin (NT), the consensus list of R-loop signals and the orientated genes. Genes were flipped according to their orientations to have the body of the gene on the left of the TTS. Orientated interaction pileups for HAP1-CTCFdegron-TIR1, in absence and presence of auxin (NT and IAA), aggregated in a 200kb window at 2kb resolution were plotted for the TTS with R-loops and the TTS without R-loops. The black circle on a stick represents the TTS and the body of the gene is represented by a dash line on the left of the TTS. (B) Stackups for active TTSs (without H3K4me3 active TSS histone mark and without CTCF peaks) for the calculated insulation in the HAP1-RPB1-AID in absence and presence of auxin (NT and IAA) and for the consensus list of R-loop signals, sorted on a consensus list of R-loops. Orientated interaction pileups for HAP1-RPB1-AID, in absence and presence of auxin (NT and IAA), aggregated in a 200kb window at 2kb resolution were plotted for the TTS with R-loops and the TTS without R-loops. The black circle on a stick represents the TTS and the body of the gene is represented by a dash line on the left of the TTS. (C) Average insulation profiles across CTCF-only peaks (with RAD21) (left plots), across scaled active genes without CTCF at TSS and TTS (middle plots) and across scaled inactive genes (right plots) for the HAP1-CTCFdegron, HAP1-CTCFdegron-TIR1 in absence and presence of auxin (NT and IAA) (first row), for the HAP1-RPB1-AID in absence and presence of auxin (NT and IAA) (second row) and for the published R-loop DRIP-seq in K562 (third row). (D) Model for a cohesin traffic pattern. Cohesin traffic pattern (green arrows) in four cell lines (Wild-type, depleted for CTCF (-CTCF), depleted for RNA polII (-RNA polII) and depleted for RAD21 (-RAD21)) for a defined chromosomal locus. Triangles represent the three types of chromatin boundaries we identified across the locus: at CTCF sites, at TSS and at TTS. In this model, cohesin is loaded at enhancers (black circle), is blocked at CTCF sites, is blocked/paused at TSS (black arrow) and is unloaded at TTS (black circle on a stick). This pattern of cohesin dynamics results in promoter-enhancer interactions, CTCF sites-promoter interactions and CTCF-CTCF sites interactions (red arrows). CTCF depletion redefines the cohesin traffic resulting in re-wired promoter-enhancer interactions. RNA polII depletion mostly affects the cohesin unloading at TTS resulting in weaker insulation at TTS. RAD21 depletion abolishes the cohesin trafficking and only keeps the insulation at TSS which seems to be mostly cohesin independent. See Figure S3

We next wanted to determine whether domain boundary formation at TTSs depends on cohesin. We made use of publicly available Hi-C data obtained with HCT116 cells depleted for the cohesin subunit RAD21 (Natsume et al., 2016; Rao et al., 2017). We again observed local insulation at TTSs of active genes in HCT116 cells (Figure S3F-G). After depletion of RAD21, this insulation was lost. Combined, these results suggest that domain boundary formation at TTSs is the result of rapid unloading of cohesin at these sites, preventing loop extrusion beyond the TTS. We conclude that domain boundary formation at active TTSs and at CTCF-bound sites both depend on cohesin (Figure S3F-G). Insulation at TSSs was still observed even after depleting RAD21. One possible explanation is that even in auxin-treated HCT116-RAD21-AID cells, some RAD21 can be detected at TSSs by ChIP-seq. This suggests that RAD21 at TSS is less efficiently depleted (Figure S3F). Similar locus-specific differences in protein depletion have been observed for CTCF degron cell lines (Luan et al., 2021). We also note that in HCT116 cells, the level of RAD21 at active TSSs was lower than the level observed in HAP1-CTCFdegron-Tir1 cells. This could be the result of the lower levels of CTCF expressed in the latter (Busslinger et al., 2017).

Combined, our analyses identify insulating domain boundary formation at at least three different types of *cis*-elements: distal CTCF-only sites with RAD21, active TSSs and active TTSs. In Figure 3C, we illustrate this general pattern by plotting the average insulation profiles across distal CTCF sites (left plots), across scaled active genes (middle plots) and across scaled inactive genes (right plots) (see definition of active and inactive genes in Methods). For this analysis, we only plotted data for active genes that did not have CTCF binding at their TSS and TTS. We noticed the existence of gene domains between TSS and TTS with higher local interactions as previously described in *Drosophila* (Rowley et al., 2019). Interestingly, this analysis revealed that depletion of CTCF not only led to loss of insulation at CTCF sites but also led to reduced interactions within active genes, as reflected in a decrease in the insulation score throughout the genes.

### Effect of RNA polymerase II depletion on boundaries at starts and ends of active genes

The formation of insulation boundaries at TSSs and TTSs was only observed at actively expressed genes (Figure 3C). We noticed that R-loops are detected at TSSs and TTSs of many active genes (Kuznetsov et al., 2018). Previous studies have linked R-loop formation to cohesin binding (Sanz et al., 2016) and cohesin stalling (Laffleur et al., 2021). Together, these observations led us to explore whether active transcription was involved in cohesin-mediated boundary formation at TSSs and TTSs. We generated a HAP1 cell line that expresses the endogenous RNA polII subunit RPB1 fused to an N-terminal and C-terminal auxin-inducible degron domain. In the presence of auxin, RPB1 was efficiently depleted within 4 hours and the cell cycle profile was not altered (Figure S3B). We performed Hi-C with HAP1-RPB1-AID cells after 4 hours of auxin treatment, and control cells treated with ethanol. Hi-C interaction frequency as a function of genomic distance between loci, compartments, TAD boundaries and CTCF-CTCF looping interactions only slightly changed after removal of RNA polII for 4 hours (Figure S3C-F). The small subset of TAD boundaries that disappeared after RNA polII depletion was likely due to high levels of RPB1 accumulation at a few sites that may result in loop extrusion blocking (one example, Figure S3E).

HAP1-RPB1-AID cells in absence of auxin already expressed a lower level of RPB1 compared to the parental HAP1 line (Figure S3B). Relative to the HAP1-WT cells, we noticed that the HAP1-RPB1-AID cells had a decrease in long range interactions that correlated with weaker A and B compartments (Figure S3C-D). We next calculated insulation genome-wide. Insulation profiles across CTCF sites were indistinguishable in RPB1 depleted cells and control cells. Insulation profiles along genes did not change much in cells depleted for RPB1, except for a notable increase in interactions across the TTS (Figure 3C, indicated by a red arrow). One possible explanation is that in control cells, transcribing RNA polII complexes push cohesin complexes towards the TTS, where they are unloaded leading to boundary formation. Upon depletion of RPB1, less cohesin is pushed towards the TTS. Alternatively, in the absence of RPB1, the local chromatin structure around TTS changes, e.g., loss of R-loops, and this prevents cohesin unloading and the associated boundary formation. We conclude that RNA polII is not required for maintenance of domain boundaries at active TSSs, but that it quantitatively contributes to boundary formation at TTSs (Figure 3B-C, Figure S3G).

### Genetic dependency between CTCF and RNA processing proteins identified by a loss of function CRISPR screen after CTCF depletion

Our results indicate that cohesin extrusion and movement along the genome is constrained by blocking sites (CTCF, TTSs), unloading sites (TTSs) and RNA polII. Combined, these features lead to the formation of chromatin domain boundaries at these sites, looping interactions between these elements, and generally elevated interactions between loci within domains. The functions of this complex cohesin “traffic” pattern are not well characterized (Figure 3D). We hypothesized that cells in which this cohesin traffic pattern is altered, e.g., through depletion of CTCF, would be particularly sensitive to genetic perturbations of functions that depend on these phenomena. To test this, we performed genome-wide CRISPR screens in HAP1-CTCFdegron-Tir1 cells (expressing reduced levels of CTCF) and compared the results to similar screens performed in HAP1-CTCF-degron cells (expressing higher levels of CTCF). We also performed screens in the presence of different concentrations of auxin to further reduce the CTCF protein level (Figure 4A, Figure S4A). Under these conditions, cell proliferation was only slightly reduced for cells in which CTCF was depleted (Figure S4B). Possibly, the remaining levels of CTCF were sufficient for growth, and/or auxin resistance emerged. We sequenced the pool of guide RNAs (sgRNAs) in HAP1-CTCFdegron-Tir1 and HAP1-CTCFdegron cell populations grown in the presence or absence of auxin and identified sgRNAs that became depleted or enriched in HAP1-CTCFdegron-Tir1 cells (with or without auxin) relative to in HAP1-CTCFdegron cells. Details of the screen are described in the Methods section. As expected, we found that sgRNAs targeting essential genes (Hart and Moffat, 2016; Hart et al., 2014, 2015) disappeared progressively over time, while most non-essential genes did not change (Figure S4C, left panels). Our screens recovered gold standard essential gene sets (Hart et al., 2015, 2017) with high precision-recall, indicating high technical quality (Figure S4C, right panels).

**Figure 4.**
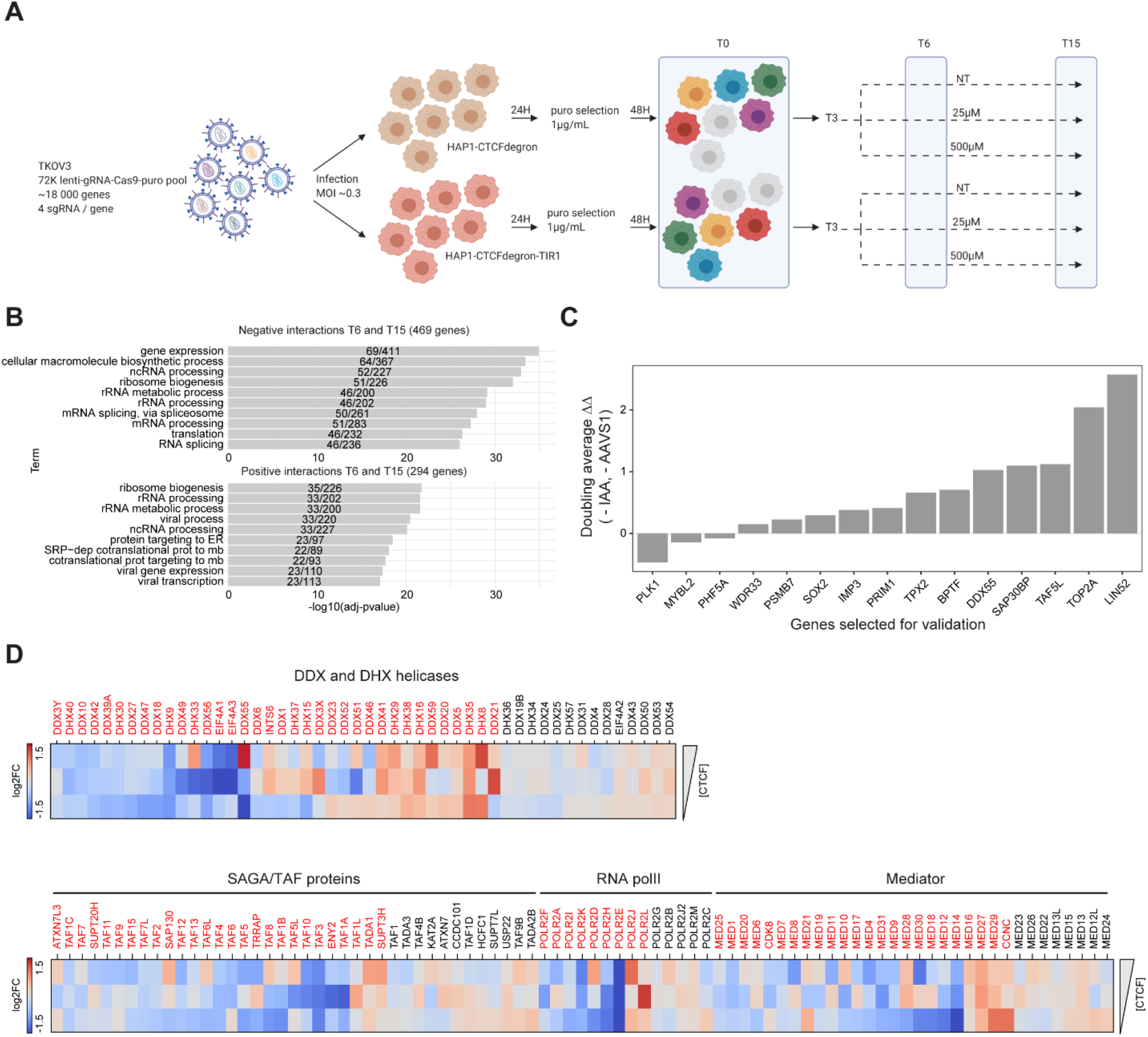
A genetic dependency between CTCF and transcription regulation factors. (A) Schematic of the CRISPR screen workflow for the two cell lines HAP1-CTCFdegron (brown cells) and HAP1-CTCFdegron-TIR1 (red cells) with three auxin conditions (NT, 25µM and 500µM). Colored cells represent lentivirus infections and knock-outs with different sgRNAs. On the represented timepoints (T0, T6 and T15), cells were harvested for genomic DNA extraction, PCR amplification, library preparation and next-generation sequencing. (B) Functional enrichment analysis for negative and positive interactions identified in the screen. (C) Validation for a selection of genes from the gene hits. Cells with proliferation defects when CTCF and the gene hit are depleted have positive values (ΔΔ). Cells with better proliferation when CTCF and the gene hit are depleted have negative values (ΔΔ). (D) Heatmaps representing the log2 fold changes normalized by the HAP1-CTCFdegron (NT, 25µM or 500µM IAA) for DDX- and DHX-box family helicases, SAGA/TAF proteins, RNA polII subunits and mediator subunits with decreasing amount of CTCF (HAP1-CTCFdegron-TIR1 in the absence of auxin (NT) and in the presence of auxin (25µM and 500µM IAA)). Genes indicated in red have a log2 fold change of > |0.4|. See Figure S4

We used two metrics to identify genes that, upon deletion, affected proliferation more strongly in cells depleted for CTCF than in cells expressing higher levels of CTCF. First, we selected genes that, when knocked out, resulted in a >= 2-fold change in sgRNA abundance in the HAP1-CTCFdegron-Tir1 cells as compared to the HAP1-CTCFdegron cells (for any of the auxin concentrations: 0µM, 25µM and 500µM auxin; see Methods). We included the no-auxin sample comparison in the analysis since the HAP1-CTCFdegron-TIR1 cell line has lower CTCF levels compared to HAP1-CTCFdegron cells, even without auxin treatment as described above (Figure S4A, D). Second, we identified genes that, when knocked out, resulted in a >= 2-fold change in sgRNA abundance in the HAP1-CTCFdegron-Tir1 cells treated with 25µM and 500µM auxin compared to HAP1-CTCFdegron-Tir1 cells grown in the absence of auxin. Finally, we removed ‘auxin-specific hits’ (see methods) (Figure S4E). Using these cut-offs, we identified a set of 469 genes whose loss reduced proliferation of HAP1-CTCFdegron-Tir1 cells compared to HAP1-CTCFdegron cells (negative genetic interactions, see Methods). We also identified 294 genes whose loss increased proliferation of HAP1-CTCFdegron-Tir1 (positive interactions, see Methods). The screen identified several genes that are known to be involved in CTCF-related processes, including SMC1A, Topoisomerase II, BPTF and LIN52. SMC1A is a cohesin subunit needed for loop extrusion and is blocked at CTCF sites, Topoisomerase II interacts with CTCF and localizes at CTCF sites (Uusküla-Reimand et al., 2016). BPTF is a subunit of the SNF2L-containing chromatin remodeler complex that has been shown to be involved in positioning nucleosomes around CTCF sites and keeping CTCF sites accessible (Barisic et al., 2019; Wiechens et al., 2016). BPTF has also been found to co-occupy CTCF sites (Valletta et al., 2020). LIN52 is part of the DREAM complex and is involved in the insulator function of CTCF in *Drosophila* and at the chicken β-globin locus (Bohla et al., 2014; Korenjak et al., 2014; Li et al., 2011).

Gene Ontology (GO) analysis revealed that the set of 469 negative interaction hits and the set of 294 positive interaction hits identified in our screens were enriched for genes involved in gene expression and RNA processing (Figure 4B). These results indicated that depletion of CTCF rendered cells vulnerable to transcription and RNA processing defects. We selected 14 hits with a broad spectrum of functions (including TPX2, BPTF, DDX55, SAP30BP, TAF5L and LIN52) for validation and included one hit from the set of positive interactions (PLK1). We knocked these genes out using two sgRNAs used in the screens and validated the negative impact on proliferation in CTCF-depleted cells for 10 of them. For PLK1, we indeed measured faster proliferation. Four hits did not validate in this assay (Figure 4C).

We noticed an enrichment of the DEAD-box helicase genes among our hits. Previous studies had identified DDX5 as an interaction partner of CTCF that plays a role at insulators in *Drosophila* and vertebrates (Lei and Corces, 2006; Yao et al., 2010). To assess proliferation defects upon DEAD-box helicase loss in the context of varying levels of CTCF depletion, we inspected the normalized log2 fold changes of fifty DEAD-box helicase genes in cells expressing high CTCF levels (untreated HAP1-CTCFdegron-TIR1 cells) to decreasing amounts of CTCF (auxin-treated HAP1-CTCFdegron-TIR1 cells). Profiles with similar patterns of changes under CTCF depletion were then manually grouped together. Genes with a differential log2 fold change of at least 0.4 were considered to produce strong changes when CTCF was depleted (see Methods). Depletion of more than two thirds of the studied DEAD-box helicases displayed proliferation defects in cells expressing lower levels of CTCF cells (36/50; both positive and negative interactions). This result indicated that DEAD-box helicases, and possibly RNA processing, might be of particular importance when CTCF is depleted (Figure 4D).

The screens also identified a number of subunits from the SAGA complex and TBP-associated factors (TAFs), PolII and Mediator complexes (Figure 4D). A previous study had shown that TAF3, which is part of the core promoter recognition complex TFIID, is recruited by CTCF to promoters and mediates looping interaction between promoter and TSS (Liu et al., 2011). The SAGA complex has histone acetylation activities and is involved in transcription activation (Baker and Grant, 2007). Mediator complexes are transcription regulators and have been shown to be involved in cohesin mediated interactions (Kagey et al., 2010; Phillips-Cremins et al., 2013). All these genes are linked to transcription regulation. This genetic dependency between CTCF and transcription regulation factors strongly point towards links between chromosome folding and regulation of gene expression.

### DDX55 and TAF5L physically interact with CTCF and cohesin

The results of our genome-wide screen suggest that cells expressing low levels of CTCF are more vulnerable to defects in machineries associated with RNA processing and transcription initiation. We selected two hits for further analysis: DDX55, a representative DEAD-box protein that showed a strong growth defect when deleted in CTCF-depleted cells and TAF5L, a subunit of the SAGA complex.

To determine whether these proteins were physically associated with CTCF and/or cohesin, we performed co-immunoprecipitation assays (co-IP) using antibodies against DDX55 and TAF5L. We found that DDX55 and TAF5L both interacted with CTCF and cohesin (RAD21 and SMC1A). This interaction was not DNA or RNA dependent since it was not altered after DNAse (turbonuclease) or RNAseA treatment (Figure 5A, Figure S5A, S5D-E). Given that CTCF and cohesin interacted with each other, we next wanted to determine whether the interactions of DDX55 and TAF5L with CTCF or with cohesin were indirect. We conducted the same co-IP after CTCF depletion with auxin treatment and found that DDX55 and TAF5L still interacted with the cohesin complex, suggesting that the interaction between DDX55, TAF5L and cohesin could occur without CTCF (Figure 5A, Figure S5A, SD-E). Additionally, we performed co-IP against TAF6L, another SAGA subunit. We also show that TAF6L interacted with CTCF and cohesin and that this interaction was DNA and RNA independent. Similar to TAF5L, the interaction between TAF6L and cohesin could occur without CTCF (Figure S5C). This result suggested that the whole SAGA complex can interact with CTCF and/or cohesin. We also took advantage of the previously generated HCT116 cell line carrying a RAD21-AID degron to examine whether the interaction between the SAGA complex and CTCF was dependent on cohesin. Using these cells, we found that the interaction between DDX55, TAF5L and CTCF was not affected by degradation of RAD21. We conclude that DDX55 and TAF5L interact with cohesin and with CTCF independently (Figure 5B, Figure S5B, SD-E).

**Figure 5.**
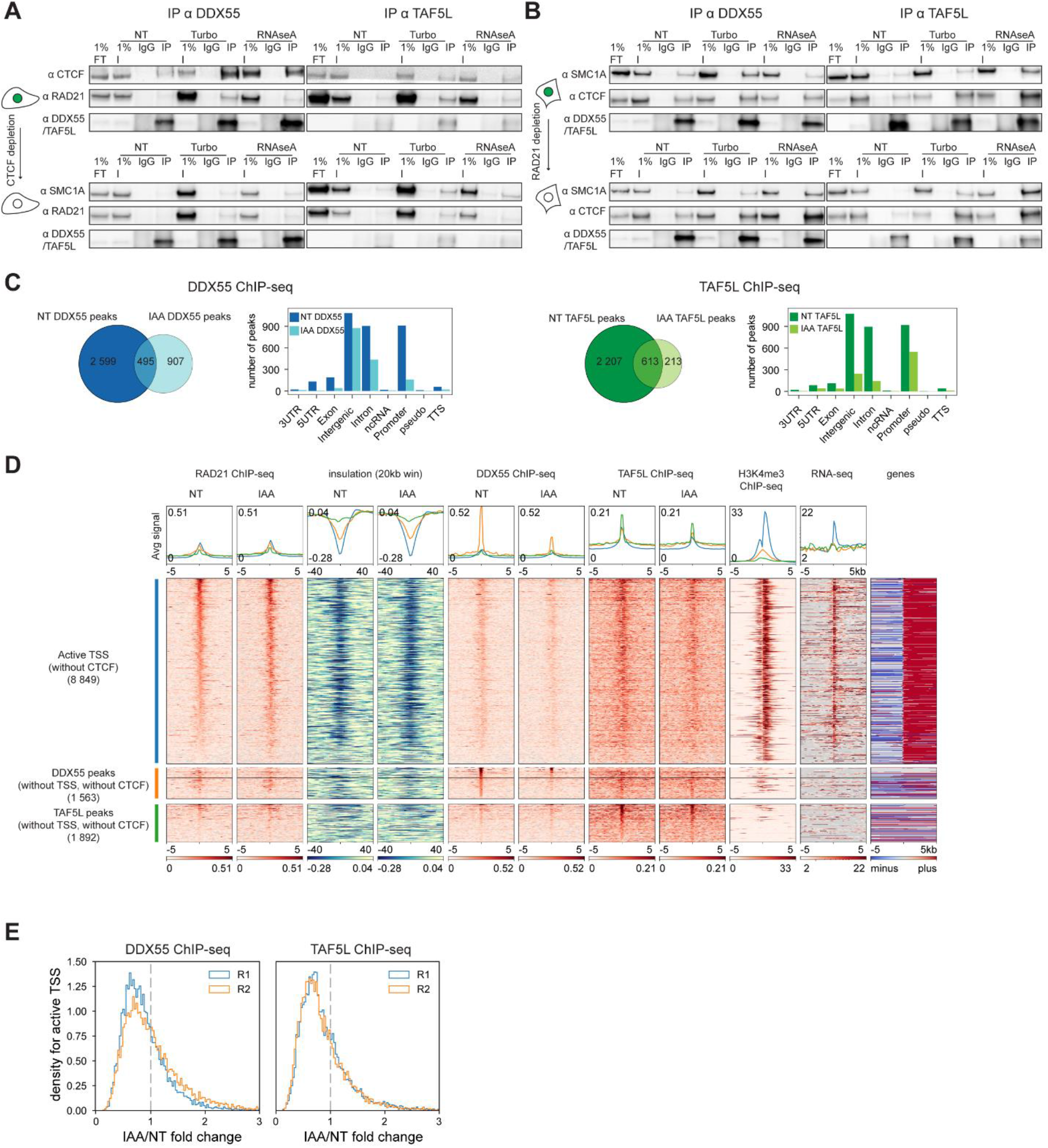
DDX55 and TAF5L physically interact with CTCF and cohesin. Chromatin binding of DDX55 and TAF5L is reduced after CTCF depletion. (A) Western blot co-IPs against DDX55 and TAF5L in HAP1-CTCFdegron-TIR1 in absence and presence of auxin (NT and IAA), treated with either turbonuclease (DNA - and RNA-) or RNAseA (RNA-). (B) Western blot co-IPs against DDX55 and TAF5L in RAD21-AID degron in absence and presence of auxin (NT and IAA), treated with either turbonuclease (DNA - and RNA -) or RNAseA (RNA -). (C) Venn diagram for DDX55 (left) and TAF5L (right) ChIP-seq peaks called in HAP1-CTCFdegron-TIR1 cells in absence and presence of auxin (NT and IAA). Gene annotation bar plots of the ChIP-seq peaks called in HAP1-CTCFdegron-TIR1 in absence and presence of auxin (NT and IAA). (D) Stackups for active TSS without CTCF (blue) sorted on Non-Treated (NT) RAD21 ChIP-seq signal, for DDX55 peaks (without TSS, without CTCF) (orange) sorted on Non-Treated (NT) DDX55 ChIP-seq, and for TAF5L peaks (without TSS, without CTCF) (green) sorted on Non-Treated (NT) TAF5L ChIP-seq. RAD21 ChIP-seq, calculated insulation, DDX55 ChIP-seq, TAF5L ChIP-seq and RNA seq signals in HAP1-CTCFdegron-TIR1 cells in absence and presence of auxin (NT and IAA) were plotted along with published HAP1 H3K4me3 ChIP-seq and orientated genes. For TSS, genes were flipped according to their orientations to have the body of the gene on the right of the TSS. Genes were not flipped for DDX55 and TAF5L peaks: genes on the forward strand are represented in red (plus) and genes on the reverse strand are represented in blue (minus), grey color corresponds to loci without annotated transcripts. (E) Stackup quantification for the active TSS (without CTCF) for DDX55 and TAF5L ChIP-seq for both replicates: a distribution of the ratios (fold change) of a given signal between auxin-treated and non-treated conditions. A fold change < 1 represents less binding of DDX55 or TAF5L at active TSS after CTCF depletion. See Figure S5

### Chromatin binding of DDX55 and TAF5L is reduced after CTCF depletion

We next wanted to assess whether chromatin binding and localization of DDX55 and TAF5L were affected by CTCF depletion. We performed DDX55 and TAF5L ChIP-seq in HAP1-CTCFdegron-TIR1 cells in the presence of CTCF (control cells) or absence of CTCF (auxin treated cells; two biological replicates). We identified 3,094 DDX55 peaks and 2,820 TAF5L peaks in the presence of CTCF. The low number of peaks for DDX55 and TAF5L ChIP-seq indicated that these factors either acted at only a limited set of loci, or generally did not bind in a highly localized focal pattern that could be detected by computational methods that call peaks (see below). Peaks were localized mostly at TSSs, introns and intergenic regions. After CTCF depletion with auxin treatment, the number of called peaks was greatly reduced (DDX55 : 1,402; TAF5L: 826) suggesting that CTCF controls DDX55 and TAF5L localization (Figure 5C, see methods for details).

Next, we determined DDX55 and TAF5L levels at the set of active TSSs that did not bind CTCF (defined in Figure 2D) and at CTCF-only sites. Very little DDX55 and TAF5L was observed at CTCF-only sites or TTSs (Figure S5G). We found that DDX55 and TAF5L were both enriched at active TSSs (Figure 5D), even though many of these loci were not identified as significant peaks above. Active TSSs that did not bind CTCF also showed strong binding of RAD21 and displayed insulation that were both independent of CTCF (Figure 5D). Interestingly, visual inspection of the ChIP-seq data suggested that after CTCF depletion with auxin, the levels of DDX55 and TAF5L binding to TSSs were reduced. We quantified this by calculating the ratio of the DDX55 or TAF5L ChIP-seq levels at each of the TSSs in control cells and CTCF depleted cells (Figure 5E). We observed that this ratio was mostly below 1, for two independent ChIP-seq replicates, confirming that binding of these factors at TSSs was reduced upon CTCF depletion. We noticed that the DDX55 and TAF5L accumulations at sites that displayed DDX55 or TAF5L peaks (above) but that did not overlap with CTCF peaks were also reduced after CTCF depletion (bottom categories, Figure 5D). We concluded that DDX55 and TAF5L positioning at active gene promoters depends on distal CTCF.

### DDX55 or TAF5L depletion has only minor effects on chromosome folding

We next asked whether DDX55 and TAF5L play roles in chromosome folding. We depleted the DDX55 and TAF5L proteins in HAP1-CTCFdegron-TIR1 cells in two ways. First, we used a pool of siRNAs to deplete DDX55 or TAF5L with an efficiency close to 100% (Figure S6A, S6E). Second, we generated DDX55 and TAF5L knock-out clones using sgRNAs also used in the genome wide CRISPR screen. We could not generate homozygous knock-outs for DDX55 because DDX55 is essential ((Bartha et al., 2018), core essential genes https://github.com/hart-lab/bagel) but succeeded in generating homozygous TAF5L knock-outs (Figure S6A, see Methods). DDX55 and TAF5L depletions with siRNA or in knock-out clones did not affect the cell cycle (Figure S6B). We noticed that depleting DDX55 and TAF5L did not affect gene expression for most of the components of the loop extrusion machinery. Only CTCF was misregulated in the DDX55 and TAF5L knock-out clones (Figure S6E). We then performed Hi-C on the DDX55 and TAF5L depleted cell lines (siRNA and knock-out clones) in the absence or presence of auxin to co-deplete CTCF. Depletion of DDX55 or TAF5L had only minor global effects on Hi-C interaction maps (Figure 6A). Compartmentalization, average loop sizes and overall distance dependence of interactions were unaffected (Figure S6C-D). To examine effects on local chromatin conformation, we again calculated average insulation profiles across distal CTCF-only RAD21 containing sites, across active genes lacking CTCF bound sites at TSS and TTS, and across inactive genes (as in Figure 3C). We found that depletion of DDX55 or TAF5L led to minor changes in the local minima in insulation scores at CTCF sites, TSSs or TTSs indicating that formation of domain boundaries at these sites did not require these factors. Interestingly, depletion of DDX55 and TAF5L changed the conformation of active genes somewhat: intragenic interactions generally increased as reflected by increased insulation scores throughout the gene domain. When CTCF was co-depleted with DDX55 or TAF5L, we found that insulation minima at CTCF sites were lost, as expected, while insulation at TSSs and TTSs was largely unaffected. Intragenic interactions were decreased compared to cells in which CTCF was not depleted. These results were comparable to the effect of CTCF depletion in cells expressing normal levels of DDX55 and TAF5L (as in Figure 3C). Therefore, the effects of CTCF depletion on intragenic interaction frequencies were independent of DDX55 and TAF5L levels. Similarly, DDX55 or TAF5L depletions in the absence of CTCF resulted in somewhat increased intragenic interactions, similar to what is observed in the presence of CTCF, suggesting that the effect of DDX55 or TAF5L depletion on intragenic interactions were independent of CTCF levels. We conclude that DDX55 and TAF5L are not required for chromatin domain boundary formation but that CTCF, DDX55 and TAF5L independently influence the conformation of active genes: CTCF depletion results in fewer intragenic interactions, whereas depletion of DDX55 or TAF5L results in more frequent intragenic interactions.

**Figure 6.**
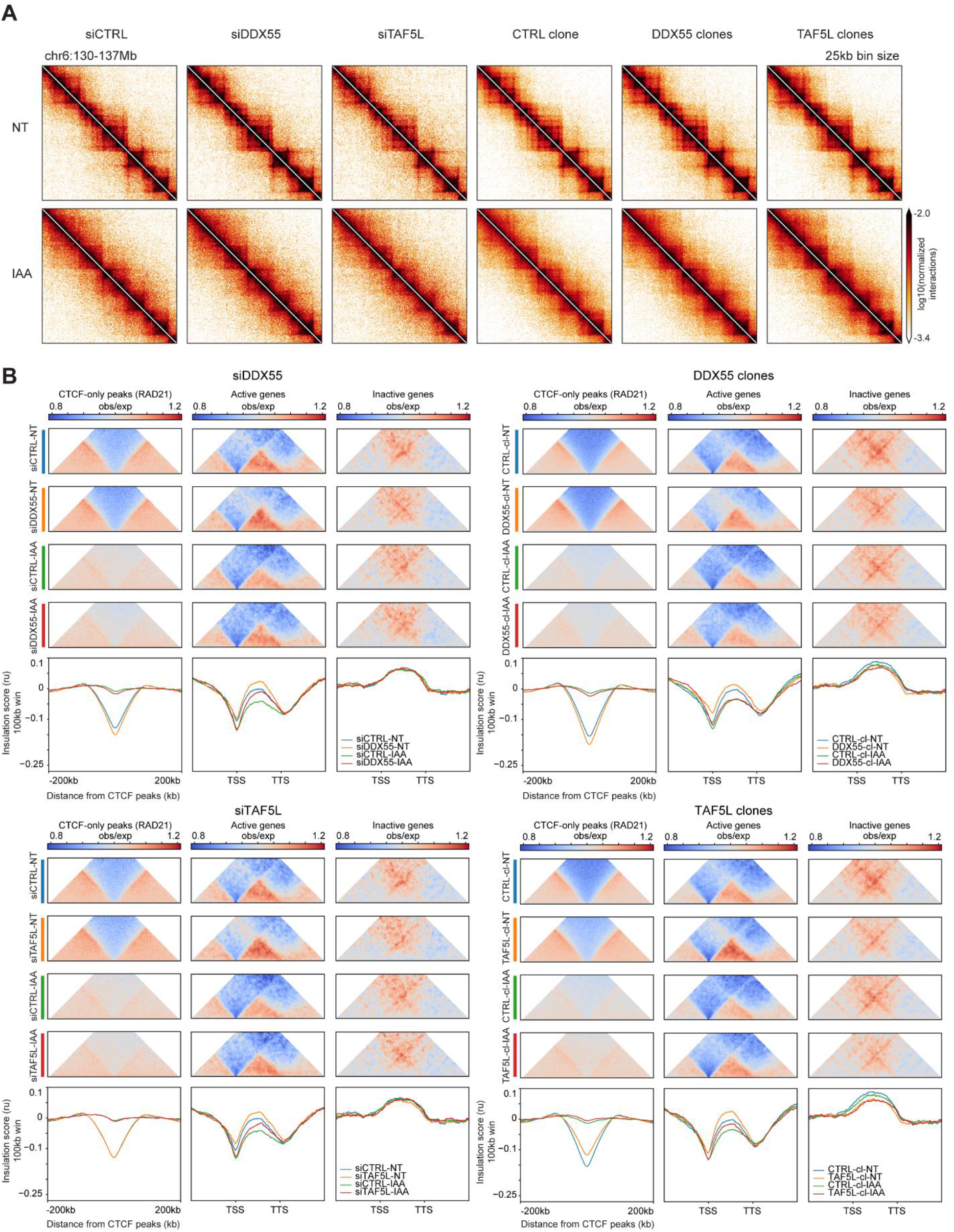
DDX55 and TAF5L depletions alter conformation of active genes independently of CTCF. (A) Hi-C contact heatmaps at 25kb resolution for a 7Mb region on chromosome 6 for HAP1-CTCFdegron-TIR1 cells depleted for DDX55 or TAF5L (siRNA and knock-out clones) in presence and absence of CTCF (NT and IAA). (B) Average insulation profiles across distal CTCF-only peaks (with RAD21) (left plots), across scaled active genes without CTCF at TSS and TTS (middle plots) and across scaled inactive genes (right plots) for the HAP1-CTCFdegron-TIR1 in absence and presence of auxin (NT and IAA) and with DDX55 or TAF5L depleted (siRNA and knock-out clones) at 5kb resolution. The representative scaled interaction pileups are plotted on top of the average insulation profiles. See Figure S6

### CTCF, DDX55 and TAF5L depletion effects on enhancer-promoter interactions and gene expression

Finally, we investigated the effect of CTCF, DDX55 and TAF5L depletion on active promoter-enhancer interactions and gene expression. For this analysis, we defined enhancers as sites that are DNAseI hypersensitive, are enriched in H3K27Ac, but are not TSSs or CTCF-bound sites. We then aggregated Hi-C data for all pairwise combinations between active TSSs and enhancers separated by 50-500kb. We split the set of enhancer-promoter pairs in two groups: those that are separated by a CTCF-bound site and those without an intervening CTCF-bound site. We also analyzed enhancers located up- and downstream of the TSS separately.

In cells expressing CTCF, we detected enriched interactions between promoters and enhancers only for those pairs that had no CTCF located in between them. After CTCF depletion, enhancer-promoter interactions got rewired: interactions of promoters with distal enhancers that were located on the other side of CTCF sites increased, whereas interactions between promoters and enhancers not separated by CTCF sites decreased. This rewiring is expected when CTCF acts as an insulator, possibly through blocking of cohesin-mediated loop extrusion. Interestingly, we noted that interactions with downstream enhancers were not as prominent as interactions with enhancers located upstream of the TSS, as had been observed in analyses of targeted gene sets (Sanyal et al., 2012).

After DDX55 and TAF5L depletions, we observed somewhat increased enhancer-promoter interactions. This increase was more readily detected in cells where CTCF was also depleted. To analyze promoter-enhancer interactions in another way, we calculated aggregated and scaled Hi-C maps for enhancer-promoter pairs with or without intervening CTCF sites (Figure S7A). This analysis showed that while insulation at TSS was readily detected, enhancers display very limited insulation, and only over short genomic distances. We again observed that depletion of CTCF resulted in rewiring enhancer-promoter interactions where interactions with enhancers across CTCF sites were increased and interactions with enhancers not separated by CTCF sites were decreased. This analysis also revealed more clearly the preference for TSSs to interact with upstream enhancers as compared to downstream enhancers. In cells where DDX55 or TAF5L were knocked out, we again observed somewhat increased promoter-enhancer interactions, both in the presence and absence of CTCF. In cells where DDX55 and TAF5L were depleted by siRNA, the effects on enhancer-promoter interactions were smaller and more variable.

Finally, we examined the effects of the orientation of the CTCF sites located in between promoters and enhancers (Figure 7B) on promoter-enhancer interactions. Limiting our analysis to the relatively small set of loci where there was only a single intervening CTCF site, or multiple that were all in the same orientation, we found that promoter-enhancer interactions were only blocked when the CTCF motif orientation was pointing towards the enhancer. For this subset of pairs, interactions increased when CTCF was depleted. The CTCF-orientation dependence strongly suggested that these interactions are 1) mediated through cohesin-dependent loop extrusion, and that cohesin is extruding from the enhancers towards the TSS. Possibly, enhancers load cohesin. In this analysis we again find that downstream enhancers interact more rarely with the TSS (Sanyal et al., 2012).

**Figure 7.**
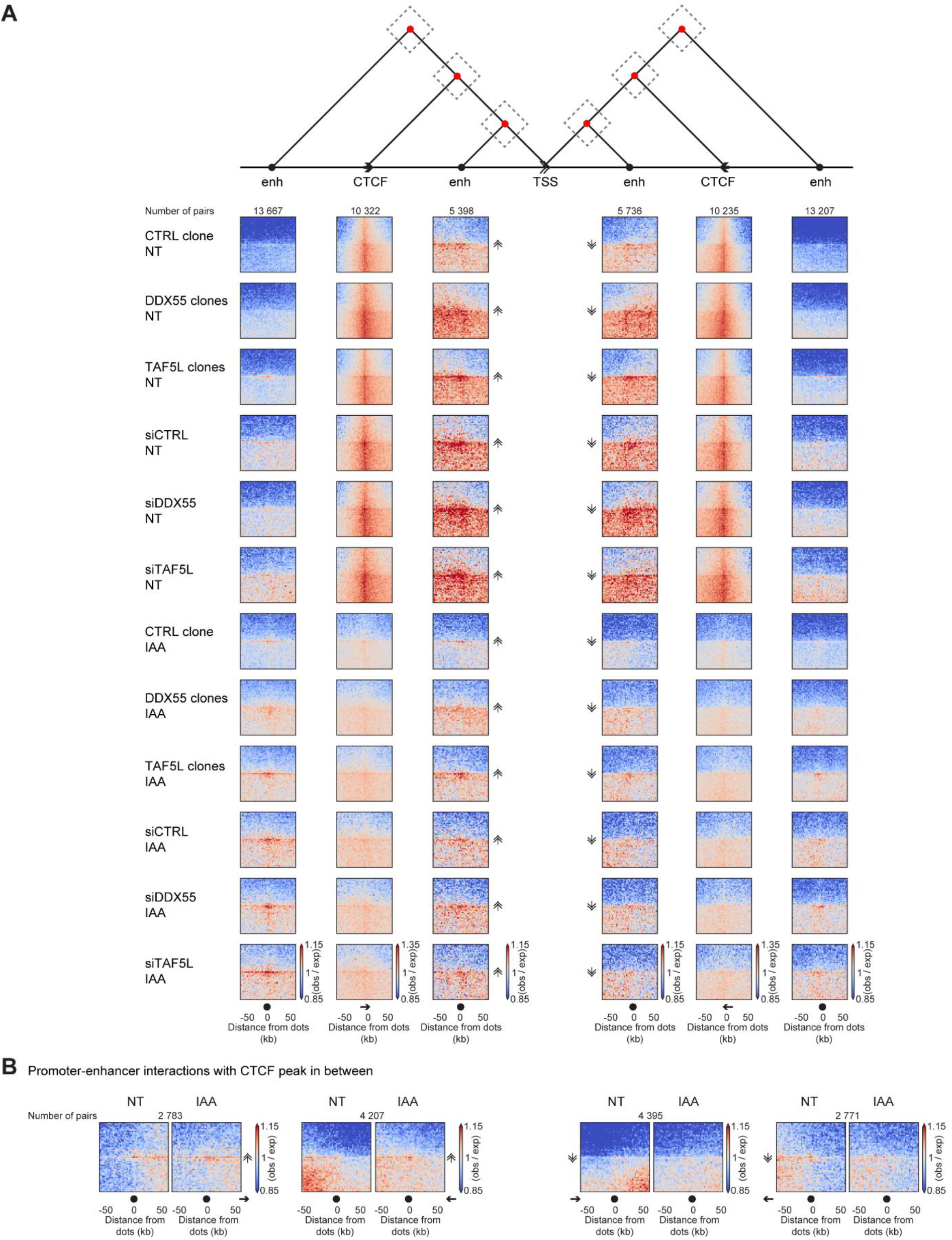
Effect of CTCF, DDX55 and TAF5L depletions on enhancer-promoter interactions and gene expression. (A) Dot pileups for all pairwise combinations between active TSSs (without CTCF) and enhancers (without CTCF or TSS) separated by 50-500kb or combinations of active TSSs (without CTCF) and CTCF-only peaks (with RAD21). Dot pileups were separated into: upstream or downstream of the TSS and having or not having a CTCF peak in between the TSS and the enhancer. The dots were aggregated at the center of a 100kb window at 2kb resolution. The schematic on top of the pileups visualizes the different studied interactions. The double arrow represents the TSS and its orientation, the black arrow represents the CTCF peak and the black circle represents the enhancer. The red dots represent interactions between enhancer and TSS or CTCF and TSS. (B) Dot pileups for all pairwise combinations between enhancer (without CTCF or TSS) and active TSSs (without CTCF) separated by 50-500kb with at least one CTCF peak in between them. Pileups were grouped by enhancer upstream or downstream of TSS, TSS orientation and CTCF motif orientation (four different combinations). See Figure S7

To see if these changes in enhancer-promoter interactions resulted in altered gene expression, we assessed the levels of RNAs after CTCF, DDX55 and TAF5L depletions by RNA-seq. Confirming previous results, CTCF depletion did not result in massive changes in gene expression (∼1,300 differentially expressed genes) (Figure S7B) (Luan et al., 2021; Nora et al., 2017; Rao et al., 2017; Schwarzer et al., 2017). However, the number of differentially expressed genes increased with the efficiency of CTCF depletion. DDX55 and TAF5L depletions weakly affected the number of differentially expressed genes. However, the double depletions of CTCF and DDX55 or CTCF and TAF5L resulted in synergistic effects with more changes in gene expression (∼600 genes). Depleting CTCF, DDX55 or TAF5L also resulted in differential splicing of a set of genes that was different from the set of genes that was differentially expressed. The number of differentially spliced genes slightly increased with the double depletions (CTCF and DDX55 or CTCF and TAF5L) (Figure S7B). Together, depleting CTCF, DDX55 and TAF5L resulted in some differentially expressed and spliced genes.

## DISCUSSION

Through analysis of Hi-C data obtained with cells in which CTCF, RAD21 or RNA polII could be rapidly depleted, we describe a complex pattern of cohesin traffic defined by different types of *cis*-elements where cohesin is loaded, paused, blocked or unloaded. Cohesin may be loaded at or near enhancers, weakly paused or blocked at TSSs, efficiently blocked and stalled at CTCF sites and unloaded at TTSs (Figure 3D). Combined, these elements determine the pattern of cohesin-mediated extrusion across the genome. Our genome-wide genetic interaction screen suggests that this elaborate pattern of cohesin-mediated chromatin extrusion, and resulting long-range interactions between promoters, enhancers and CTCF sites, is involved in transcription initiation and RNA processing.

### A cohesin traffic pattern defined by three types of chromatin boundaries

We explored chromatin folding at the scale of tens of kilobases genome-wide and found that at least three distinct types of *cis*-elements induce chromatin insulation, i.e. represent locations across which chromatin interactions are depleted resulting in domain boundary formation. For instance, CTCF-bound elements are known to block cohesin-mediated loop extrusion in a binding site orientation dependent manner (Guo et al., 2015; Rao et al., 2014; de Wit et al., 2015). Blocking of extrusion will prevent long-range interactions across CTCF-bound sites. Consistently, we detected strong insulation at the subset of CTCF-bound sites where we also observed RAD21 accumulation, and this insulation was both CTCF and RAD21 dependent.

We describe two additional types of *cis*-elements that display chromatin insulation: TSSs and TTSs of active genes. Insulation at TSSs and TTSs is quantitatively distinct from insulation at CTCF sites, and appears to be caused by different mechanisms. To rule out confounding effects of CTCF binding to a subset of TSSs and TTSs, we focused our analyses on those that are devoid of CTCF binding sites. TSSs display relatively strong, but highly localized insulation that is quantitatively comparable to the observed insulation at CTCF-bound sites. Cohesin is detected at TSSs, but compared to the level of cohesin accumulation at CTCF sites, RAD21 enrichment at TSSs is relatively low, even in CTCF depleted cells where cohesin is known to be redirected towards TSSs (Busslinger et al., 2017). Therefore, it appears that the strong insulation observed at TSSs does not require high steady state levels of cohesin accumulation. Further, while rapid depletion of RAD21 leads to near complete loss of insulation at CTCF-bound sites, insulation at TSSs is hardly affected. Interestingly, the level of RAD21 binding to TSSs in RAD21 depleted cells is only somewhat reduced, in contrast to RAD21 levels at CTCF sites that are strongly reduced. Locus-specific differences in efficiency of factor depletion was recently also described for CTCF (Luan et al., 2021). Furthermore, we noted that the relative level of RAD21 accumulation at active TSSs in HCT116 cells is even lower, but that insulation at TSSs in those cells is still readily detected (Figure S3F). Clearly, strong insulation at TSSs can be present even with very small amounts of RAD21 binding. Alternatively, insulation at TSSs is not dependent on cohesin at all, but instead is the result of other loop extrusion factors or is driven by different mechanisms. We note that active TSSs in *Saccharomyces cerevisiae* also display strong insulation (Hsieh et al., 2015). This is interesting because in that organism there is no cohesin bound to chromatin in G1. Possibly, active TSSs have specific local chromatin features that can induce chromatin domain boundary formation via alternative mechanisms. They typically contain large open chromatin regions and many TSSs have sequences that can form G-quadruplex (G4) structures. A previous study identified G4s as being enriched at TAD boundaries. This study showed that genome-wide G4s have higher levels of CTCF and cohesin binding and display increased insulation (Hou et al., 2019). Recent studies highlight that the Myc-associated zinc finger protein (MAZ) can act as an insulating cofactor with CTCF. It can also function as a boundary independently of CTCF and interacts with cohesin (Ortabozkoyun-Kara et al., 2020; Xiao et al., 2021b). Interestingly, MAZ can bind and unfold G-quadruplex structures (Cogoi et al., 2010, 2014). Yin Yang 1 (YY1), which is known to contribute to enhancer-promoter interactions, can directly bind to G4s (Weintraub et al., 2017). Stabilizing G4 structures with drugs like PDS or TMPyP4 affects YY1 binding and genome organization (Li et al., 2021). The binding of MAZ and YY1 to G4, specifically at TSSs, associated or not with CTCF, could either make strong barriers for loop extrusion or favor a local chromatin environment that facilitates domain boundary formation through some other mechanism.

The third element that forms domain boundaries as reflected in insulation is found at the TTS of active genes. We again focused on the majority of TTSs that do not contain CTCF binding sites. At these sites, insulation is not affected by CTCF depletion. Insulation at TTSs is quantitatively distinct from that observed at CTCF sites and TSSs: it is weaker and forms a broad zone of insulation. Intriguingly, we did not detect RAD21 at active TTSs by ChIP-seq, but insulation at TTSs is lost in cells where RAD21 is depleted. Previous studies had shown that in cells depleted for CTCF and the cohesin unloading factor WAPL, cohesin accumulates at 3’ ends of active genes and especially at sites of convergent transcription (Busslinger et al., 2017). Combined, these observations indicate that active TTSs are sites where, in normal cells, cohesin is unloaded in a WAPL-dependent manner. The lack of RAD21 detection by ChIP-seq can be explained when unloading is fast and efficient. We propose efficient unloading of cohesin at TTSs produces insulation because cohesin cannot extrude past the unloading site. Insulation at TTSs is partly dependent on RNA polII: depleting RNA polII results in weaker insulation at TTSs.

Blocking transcription elongation using DRB in CTCF WAPL double knock-out cells results in less accumulation of cohesin at TTSs (Busslinger et al., 2017). One hypothesis is that the weakening of R-loops by RNA polII removal results in less unloading of cohesin at TTSs and thus weaker insulation. An alternative hypothesis could be that cohesin is pushed through the gene by RNA polII towards the TTS where it is unloaded. Previous studies have shown that condensin and RNA polII can interplay (Brandão et al., 2019) and it has been shown in yeast that cohesin could be pushed by the transcription machinery (Lengronne et al., 2004). Either model suggests interplay and interference between extruding cohesin and RNA polymerase. This interference may also contribute to the lower frequency of interaction of TSSs with downstream enhancers.

We noticed that TTSs with the strongest insulation also appear to display relatively high levels of R-loops. Possibly R-loops favor cohesin unloading at TTSs. Pan and co-workers linked R-loops with loop extrusion factors. They found that the cohesin subunits STAG1 and STAG2 bind R-loops *in vitro*, and that STAG1 and STAG2 ChIP-seq binding sites overlap with R-loops (Pan et al., 2020).

Taken together, each of the three elements that display insulation and boundary formation activity differ in the mechanisms by which they do so: insulation at CTCF sites and TTSs both depend on cohesin but for different reasons. Cohesin is stalled and stabilized at CTCF sites, and cohesin is efficiently unloaded at TTSs. Insulation of TSSs appears largely independent of the level of RAD21 accumulation, and other processes may contribute. Finally, enhancers display only very weak insulation (Figure S7A). Enhancer-promoter interactions are directed by CTCF site orientation in a way that suggests that cohesin could be loaded at enhancers and extrude towards the promoter. This complex and dynamic cohesin traffic pattern may be important for appropriate gene regulation, e.g., through recruiting and then delivering transcription related complexes to target genes. A similar model for cohesin dynamics has been proposed by Liu and co-workers based on analysis of cells where either WAPL or RAD21 are depleted (Liu et al., 2021). Both WAPL and RAD21 depletions lead to loss of cohesin dynamics and, in the case of WAPL depletion, accumulation of cohesin at CTCF sites and loss of cohesin from dynamic sites.

The vertebrate V(D)J recombination system has proven to be an excellent model to study the interplay between loop extrusion, CTCF sites and R-loops (for a review on V(D)J recombination system and loop extrusion: (Peters, 2021)). R-loops have been shown to form at the IgH switch regions and to be crucial for efficient class switching (Daniels and Lieber, 1995; Reaban and Griffin, 1990; Yu et al., 2003). A DEAD-box helicase, DDX1 is required for class switch recombination (CSR). DDX1 binds to RNA G4s that result in R-loop formation at the IgH switch regions (Ribeiro de Almeida et al., 2018). A very recent study links the loop extrusion factor cohesin and R-loops (Laffleur et al., 2021). These authors find a correlation between R-loop accumulation after depleting DIS3, a component of the RNA exosome, and cohesin displacement at the CSR locus. These studies bring together CTCF, G4s, R-loops and cohesin for CSR at the IgH locus. Our study suggests that these factors contribute to chromatin folding and domain boundary formation genome-wide.

### Function of the cohesin traffic pattern

Depletion of cohesin or CTCF changes chromosome folding genome-wide but leads to altered gene expression for only a relatively small set of genes within the first hours. Therefore, analysis of (immediate) effects on transcription has not been very informative for studying roles of cohesin and CTCF on gene regulation. We therefore used a different approach, genetic interaction analysis, to explore possible functions for the cohesin traffic pattern. We identified factors that, upon deletion, changed the growth rate of cells only when CTCF levels were low and cohesin positioning along chromosomes was altered. We identified several classes of genes involved in RNA processing, R-loop formation and transcription initiation. These synthetic interactions with CTCF depletion suggest that CTCF plays roles in various aspects of gene expression. First, many DEAD-box containing RNA helicases were identified. This is a large family of proteins involved in a variety of processes including R-loop formation/unwinding, splicing and RNA processing in general (García-Muse and Aguilera, 2019). In previous studies, several DEAD-box helicase proteins have been implicated in CTCF functions. DDX5 and its associated RNA activator RSA have been identified as CTCF- and cohesin-interacting factors that are required for the insulator function of CTCF (Yao et al., 2010), possibly by reducing cohesin localization at CTCF sites. In *Drosophila*, Rm62 has been implicated in the activity of the insulator binding factor CP190 (Lei and Corces, 2006). Second, we identified a set of proteins implicated in transcription initiation, such as TAFs, that are part of the SAGA, TFIID, and RNA polII complexes. These complexes are important for both basal transcription and enhancer-driven activated gene expression.

We studied the roles of DDX55 and TAF5L in more detail. Our results indicate that DDX55 and TAF5L do not appear to play a major role in chromatin folding: their depletion mostly did not affect compartmentalization or domain boundary formation at CTCF, TSS and TTS sites, or formation of loops between these elements. DDX55 or TAF5L depletion did alter intra-genic interaction frequencies somewhat and independently of CTCF.

Interestingly, we found that CTCF depletion leads to reduced accumulation of DDX55 and TAF5L at CTCF sites and at active TSSs including those that do not contain CTCF-binding sites. This observation points to an indirect role for CTCF in recruiting and positioning these factors and possibly other transcription related complexes to distal active genes most likely through cohesin mediated mechanisms. DDX55 and TAF5L could get recruited to distal CTCF sites and then be transported to TSSs through cohesin action. Consistent with this model, we found that DDX55 and TAF5L physically interact with both CTCF and cohesin.

Depletion of DDX55 or TAF5L, in the presence or absence of CTCF, did not result in major and global changes in gene expression and splicing as measured by RNA-seq. This was not unexpected given that previous studies had also found that depletion of CTCF or cohesin only affected expression of a small number of genes (Luan et al., 2021; Nora et al., 2017; Rao et al., 2017; Schwarzer et al., 2017). Another study showed that removing all the major CTCF binding sites in the mouse Sox9-Kcnj2 TAD boundary results in insulation loss at this boundary without major changes in gene expression (Despang et al., 2019). It has also been shown that CTCF can regulate splicing and alternative polyadenylation of mRNA (Alharbi et al., 2021; Nanavaty et al., 2020; Ruiz-Velasco et al., 2017; Shukla et al., 2011). However, the number of genes with splicing defects in these studies is similar to what we found in our study. Similarly, in yeast some components of the SAGA complex can be homozygous knocked out without causing major expression phenotypes (Gaillard et al., 2009). Explanations for the lack of major transcriptional changes include redundancy with other related complexes (e.g., other histone acetyl transferases), or the timescale of events (here hours). Recent analyses suggest that transcriptional state of a TSS can be relatively long-lived so that acute depletion of factors that mediate enhancer-driven activation do not have a noticeable effect of transcription until many hours, or even cell cycles later (Xiao et al., 2021a; Zuin et al., 2021).

In summary, our work delineates roles of CTCF, cohesin and RNA polII in defining a local chromatin folding landscape with different types of domain boundaries at key *cis*-elements. Defects in setting up this landscape correctly, e.g. when CTCF is depleted, make cells sensitive to loss of factors involved in RNA processing and transcription initiation. We propose that the complex pattern of cohesin movement along chromatin, and the roles of CTCF and RNA polII in defining this pattern, contributes to appropriate localization of transcription and RNA processing factors to active genes. How these phenomena control gene expression remains an open question.

## AUTHOR CONTRIBUTIONS

A.-L.V. and J.D. conceived and designed the study. A.-L.V. engineered cell lines, performed Hi-C, ChIP-seq, RNA-seq. A.-L.V. and S.V.V. analyzed Hi-C, ChIP-seq, RNA-seq and other relevant datasets. A.-L.V., B.M., A.H.Y.T., J.M. designed the strategy for the CRISPR screens. generated the lentiviruses for the CRISPR screens. A.-L.V. and B.M. performed the CRISPR screens. A.-L.V., B.M., M.U. analyzed the CRISPR screen data. E.K. and A.A.P. analyzed splicing in the RNA-seq data. A.-L.V. and J.D. wrote the manuscript with input from all authors.

## METHODS

### RESOURCE AVAILABILITY

#### Lead contact

Further information and requests for resources and reagents should be directed to and will be fulfilled by the lead contact, Job Dekker.

#### Materials availability

All unique reagents generated in this study are available from the lead contact upon request.

#### Data and code availability

- The data datasets generated in this publication have been deposited in NCBI’s Gene Expression Omnibus (Edgar et al., 2002).

- All original code has been deposited in GitHub.

- Any additional information required to reanalyze the data reported in this paper is available from the lead contact upon request.

### EXPERIMENTAL MODEL AND SUBJECT DETAILS

#### Cell culture and cell lines

Human HAP1 cell line was purchased from Horizon Discovery. The wild-type and mutated HAP1 cell lines (HAP1-CTCFdegron, HAP1-CTCFdegron-TIR1, HAP1-RPB1-AID, DDX55 and TAF5L knock-out clones) were cultured at 37°C with 5% CO2 in IMDM GlutaMAX™ Supplement (Gibo, 31980097) with 10% FBS (Gibco, 16000069), 1% Penicillin-Streptomycin (Gibco, 15140122).

HCT116-RAD21-AID cells were a gift from Masato Kanemaki (Natsume et al., 2016). They were cultured at 37°C with 5% CO2 in McCoy’s 5A medium GlutaMAX™ Supplement (Gibco, 36600021) with 10% FBS (Gibco, 16000069), 1% Penicillin-Streptomycin (Gibco, 15140122).

Cell lines were routinely tested for mycoplasma infection and tested negative (MycoAlertTM Mycoplasma Detection Kit, Lonza).

#### Antibiotic selection treatment

Blasticidin S HCl (10mg/mL) was ordered from ThermoFisher (A1113903). Selection was done with 10µg/mL blasticidin.

Puromycin Dihydrochloride (10mg/mL) was ordered from ThermoFisher (A1113803). Selection was done with 1.5µg/mL puromycin.

Hygromycin B Gold (100 mg/mL) was ordered from Invivogen (ant-hg-1). Selection was done with 450µg/mL hygromycin.

#### Auxin (IAA) treatment

Auxin (IAA, 3-Indoleacetic acid) was purchased from Millipore Sigma (45533-250MG) and dissolved in ethanol. Auxin was directly added to the cell culture plates at the indicated concentrations (25µM for partial CTCF depletion or 500µM for total CTCF, RPB1 and RAD21 depletions) and times (HAP1-CTCFdegron and HAP1-CTCFdegron-TIR1 : 48H for the asynchronous cells, 12H for the G1 sorted cells, HAP1-RPB1-AID : 4H, HCT116-RAD21-AID : 2H).

#### siRNA transfections

Pool of siRNA were ordered from Dharmacon (siGENOME Non-Targeting siRNA Pool #2, SMARTpool: siGENOME DDX55 siRNAl and siGENOME TAF5L siRNA). siRNAs were resuspended in sterile ultra-pure water. Transfections were done with lipofectamine (Lipofectamine™ RNAiMAX Transfection Reagent, Thermofisher Scientific, 13778075) and Opti-MEM (Thermofisher Scientific, 31985062) following the manufacturer’s recommendations. Final concentration of siRNA used was 40nM and incubated for 72 hours. If auxin was needed, media was removed after 24 hours and replaced by auxin containing media for the remaining 48 hours.

### METHOD DETAILS

#### Plasmid construction

Each plasmid was analyzed by Sanger sequencing to confirm successful cloning.

##### guide RNA cloning (sgRNA)

sgRNA were cloned in pSpCas9(BB)-2A-Puro (PX459) V2.0 (Feng Zhang laboratory, Addgene 62988). Briefly, the pX459 plasmid was digested with BbsI, the sgRNA primers were phosphorylated, annealed and ligated into the BbsI linearized backbone following the Feng Zhang laboratory protocol (Ran et al., 2013).

##### Endogenous CTCF knock-out targeting constructs

To knock-out the endogenous CTCF, sgRNAs targeting the promoter and the 3’ UTR of the endogenous CTCF gene were cloned (∼79kb deletion).

##### CTCF cDNA construct: HA-AID-CTCFcDNA-AID-eGFP-blasticidin

The HA-AID-CTCFcDNA-AID-eGFP-blasticidin vector was assembled by Gibson Assembly (NEBuilder HiFi DNA Assembly Master Mix, NEB, E2621L) in the pENTR221 kanamycin vector using the following templates: the CAGGS promoter (which contains the cytomegalovirus (CMV) early enhancer element, the promoter region, the first exon, and the first intron of chicken β-ACTIN gene, and the splice acceptor of the rabbit β-GLOBIN gene) was amplified from pEN396-pCAGGS-Tir1-V5-2A-PuroR (gift of Elphege Nora, Benoit Bruneau, addgene 92142), the minimal functional AID tag (aa 71-114) was amplified with forward primer containing HA tag from pEN244-CTCF-AID[71-114]-eGFP-FRT-Blast-FRT (gift of Elphege Nora, Benoit Bruneau, addgene 92140), the CTCF cDNA was amplified from a pCMV6-Entry vector containing CTCF cDNA (Origene, RC202416), the AID-eGFP-2A-bls was amplified from pEN244-CTCF-AID[71-114]-eGFP-FRT-Blast-FRT (gift of Elphege Nora, Benoit Bruneau, addgene 92140), the polyA signal was amplified from pEN396-pCAGGS-Tir1-V5-2A-PuroR (gift of Elphege Nora, Benoit Bruneau, addgene 92142). Amplifications were performed with the Q5 High-Fidelity DNA Polymerase (NEB, M0491L).

##### TIR1-hygro construct

The TIR1-hygro vector was assembled by Gibson Assembly (NEBuilder HiFi DNA Assembly Master Mix, NEB, E2621L) replacing puromycin gene by hygromycin gene in the pEN396-pCAGGS-Tir1-V5-2A-PuroR (gift of Elphege Nora, Benoit Bruneau, addgene 92142).

##### Endogenous RPB1 targeting constructs

###### C-terminal

To target the C-terminal part of RPB1, sgRNA targeting the last exon of RPB1 gene was cloned.

###### N-terminal

To target the N-terminal part of RPB1, sgRNA targeting the first exon, around the start codon of RPB1 gene was cloned.

##### AID C-terminal RPB1-AID-eGFP-blasticidin construct

The AID C-terminal RPB1-AID-eGFP-blasticidin vector was assembled by Gibson Assembly (NEBuilder HiFi DNA Assembly Master Mix, NEB, E2621L) in the pENTR221 kanamycin vector using the following templates: the 5’ homology arm (1 680bp) and 3’ homology arm (1 558bp) were amplified from HAP1 genomic DNA, the minimal functional AID tag (aa 71-114)-eGFP was amplified from pEN244-CTCF-AID[71-114]-eGFP-FRT-Blast-FRT (gift of Elphege Nora, Benoit Bruneau, addgene 92140), the T2A was amplified from pEN396-pCAGGS-Tir1-V5-2A-PuroR (gift of Elphege Nora, Benoit Bruneau, addgene 92142), the blasticidin resistance gene was amplified from PSF-CMV-BLAST (Sigma-Aldrich, OGS588-5UG). Amplifications were performed with the Q5 High-Fidelity DNA Polymerase (NEB, M0491L).

##### AID N-terminal AID-RPB1 construct

The AID N-terminal AID-RPB1 vector was assembled by Gibson Assembly (NEBuilder HiFi DNA Assembly Master Mix, NEB, E2621L) in the pENTR221 kanamycin vector using the following templates: the 5’ homology arm (1 079bp) and 3’ homology arm (1 077bp) were amplified from HAP1 genomic DNA and the minimal functional AID tag (aa 71-114)-eGFP was amplified from pEN244-CTCF-AID[71-114]-eGFP-FRT-Blast-FRT (gift of Elphege Nora, Benoit Bruneau, addgene 92140).

##### AAVS1 (control locus), DDX55 and TAF5L knock-out constructs

To create deletions in the AAVS1 locus, primers were designed in the AAVS1 locus. To create DDX55 knock-out, the sgRNAs used in the genome wide CRISPR screen and targeting the second exon of DDX55 gene were cloned. To create TAF5L knock-out, the sgRNAs used in the genome wide CRISPR screen and targeting the third exon of TAF5L gene were cloned.

#### Genome modifications

Plasmids used for transfections were purified using ZymoPURE II Plasmid Midiprep Kit (Zymo Research, D4201). Plasmids were linearized using PvuI-HF (NEB, R3150L). Linearized plasmids were further purified with phenol chloroform extraction and ethanol precipitation. HAP1 cells were transfected using turbofectin (Origene, TF81001) following the manufacturer’s recommendations.

The differences between the different construct transfections are described below:

##### HAP1-CTCFdegron-TIR1

1.5µg of linearized HA-AID-CTCFcDNA-AID-eGFP-blasticidin vector was transfected. 24 hours after the transfection, blasticidin (10µg/mL) containing media was added and resistant cells were selected for 48 hours. A second transfection was then performed using 2µg of four sgRNA-CRISPR-vectors (4*0.5µg) on the pool of blasticidin resistant cells to knock-out the endogenous CTCF. After 24 hours, puromycin (1.5µg/mL) containing media was added and resistant cells were selected for 48 hours. Serial dilutions were then done on 96-well plates without antibiotic selection to generate single cell clones. To test for integration of HA-AID-CTCFcDNA-AID-eGFP-blasticidin and effective CTCF knock-out, cells from individual clones were trypsinized, half was left in the 96-well plate and the other half was used for genomic DNA extraction. Clones that harbored the endogenous CTCF knock-out and the integration of the HA-AID-CTCFcDNA-AID-eGFP-blasticidin construct were sequenced. Clone (referred to as HAP1-CTCFdegron in our study) with the correct sequence was used for TIR1 integration. 2µg of linearized TIR1-hygro vector were then transfected into the HAP1-CTCFdegron clone. After 24 hours, hygromycin (450µg/mL) containing media was added and resistant cells were selected for 48 hours. Serial dilutions were then done on 96-well plates without antibiotic selection to generate single cell clones. Clones were then PCR tested and sequenced for correct TIR1 integration on single clones. The clone used in this study is referred to as HAP1-CTCFdegron-TIR1.

##### HAP1-RPB1-AID

1.5µg of linearized RPB1-AID-eGFP-blasticidin vector and 1.5µg of C-terminal RPB1 sgRNA were transfected into HAP1 cells. 24 hours after the transfection, puromycin (1.5µg/mL) containing media was added and resistant cells were selected for 48 hours. Puromycin media was then washed, and cells were grown for 48 hours without antibiotics. Blasticidin resistant cells were selected by adding blasticidin (10µg/mL) containing media for 7 days. The pool of blasticidin resistant cells was then transfected with 2µg of linearized TIR1-hygro vector. After 24 hours, serial dilution of cells to select single cell clones were performed in hygromycin (450µg/mL) containing media. Clones were then PCR tested and sequenced for correct AID-eGFP and TIR1 integrations on single clones.

##### AAVS1, DDX55 and TAF5L knock-outs

2µg of sgRNA targeting the AAVS1, DDX55 and TAF5L loci were transfected into HAP1 cells. 24 hours after the transfection, puromycin (1µg/mL) containing media was added and resistant cells were selected for 48 hours. Serial dilutions of cells in media without selection were then done to select single cell clones. Clones were PCR tested and sequenced for indels on both alleles. AAVS1 clone (control) harbors a 23bp deletion on both alleles. DDX55 clone 1 harbors one allele with a 3bp deletion, deleting deux amino acids (I and P) and replacing it by another one (T). The second allele has a 4bp deletion creating a frameshift and premature stop codon in exon 3. DDX55 clone 2 harbors one allele with a 6bp deletion, deleting three amino acids (PLF) and replacing it by another one (L). The second allele has a 12bp deletion deleting 4 amino acids (ATIP). The amount of mutated DDX55 protein is reduced in both clones. TAF5L clone 1 is homozygous with a 7bp deletion in the third exon of the TAF5L gene, creating a premature stop codon. TAF5L clone 2 is homozygous with a 13bp deletion creating a premature stop codon. These TAF5L knock-out clones do not express the TAF5L protein.

#### Genomic DNA extraction for PCR to test clones

Cells were spun, resuspended in 30µL of SB buffer (10mM Tris pH 8.0, 25mM NaCl, 1mM EDTA, 200µg/ml Proteinase K), incubated 1 hour at 65°C and 10min at 95°C, spun and 1µL of the supernatant was used for PCR.

### CRISPR screen validation

Validation was performed on 16 genes, with 2 different sgRNA targeting the gene of interest on HAP1-CTCFdegron-TIR1 cells.

2µg of targeting sgRNA plasmids were transfected (separately for the sgRNA targeting the same gene) using turbofectin (Origene, TF81001) following the manufacturer’s recommendations. After 24 hours, puromycin (1.5µg/mL) containing media was added to select cells that integrated the plasmids. After 48 hours, cells were counted and timempoint considered as T0. Passaging was then performed following the scheme used in the genome-wide CRISPR screen. Three days later (T3), cells were counted, and re-seeded into three conditions (NT, 25µM and 500µM auxin) in duplicates in 24-well plates. Cells were counted and re-seeded for the three conditions every three days until reaching T15. Cumulative growth curves were plotted with the number of counted cells. We calculated the doubling average ΔΔ, by first calculating the cumulative doubling averages per gene (two sgRNA per gene) for each time point. Then, we subtracted the cumulative doubling of the auxin treated from the non-treated (NT - IAA) per gene for each time point. Subsequently, we subtracted the control value (AAVS1) per gene for each time point. Finally, we calculated the mean of all time points for each experiment replicate. A positive ΔΔ value indicates a growth defect when the gene is knocked out and CTCF is depleted. A negative ΔΔ value indicates a better proliferation when the gene is knocked out and CTCF is depleted. To confirm that indels occurred, cells were harvested at T15 and genomic DNA extraction was performed. PCR was then done on the extracted gDNA from cells that went through the transfections (mutated amplicon) and for cells that were not transfected (Wild-type amplicon). PCR products were purified using GFX PCR DNA and Gel Band Purification Kit (Cytiva, 28903470) and sent for Sanger sequencing. Synthego (https://www.synthego.com/products/bioinformatics/crispr-analysis, (Hsiau et al., 2019) was then used to assess the percentage of the different modified alleles in the targeted genes using the wild-type amplicons as controls.

#### Flow cytometry

Cells were dissociated with accutase (ThermoFisher Scientific, A11105-01), resuspended in PBS, spun, and resuspended in 250µL of cold PBS. To assess the cell cycle profile (DNA content), 750µL of 100% ethanol was slowly added to fix cells in 75% ethanol. Cells were stored in -20°C for at least 24 hours. Fixed cells were spun, re-suspended in 1X PBS with propidium iodide (PI) (final concentration 50µg/mL) and RNAseA (0.5mg/mL) and incubated for 30 min at room temperature protected from light. To assess GFP content, cells were washed once with PBS and fixed with 4% PFA for 10 min. Cells were spun and cell pellets were resuspended in 1mL of PBS. Cells were sorted on a FACSCALIBUR or LSRII or MACSQUANT. Analysis was performed using the Flowjo software.

#### FACS sorting for Hi-C

Cells were prepared for FACS sorting following (Bonev et al., 2017).

Briefly, cells were detached using StemPro Accutase Cell Dissociation Reagent (Thermo Fisher Scientific, A1110501), spun and resuspended in 1% formaldehyde HBSS solution. Cells were fixed for 10 min at room temperature. To quench the formaldehyde and terminate the cross-linking reaction 125mM glycine was added and incubated for 5 min at room temperature. Cells were then incubated on ice for 15 min. Formaldehyde solution was then removed and cells were washed once with cold dPBS. Cells were resuspended in 0.1% saponin in PBS to have 1M cells/mL and 1µL/mL of fxCycle far red dye together with 10µL/mL RNAse at 10mg/mL were then added. Cells were incubated 30 min at room temperature protected from light. Cells were then spun and resuspended in 0.5% BSA in PBS to have a final concentration of cells of 10M cells/mL. Cells were sorted on a BD FACSAria Cell Sorter. For the HAP1-CTCFdegron-TIR1 in absence of auxin, G1 cells that were GFP positive were sorted (CTCF +). For the HAP1-CTCFdegron-TIR1 in presence of auxin, G1 cells that were GFP negative were sorted (CTCF -). About 0.5-2M cells were sorted. If Hi-C was not performed immediately after cell sorting, pellets were flash frozen in liquid nitrogen and kept at -80°C until starting the cell lysis.

#### Western blots

Cells were dissociated with accutase (ThermoFisher Scientific, A11105-01), resuspended in PBS, spun, washed with PBS, spun again and kept at -20°C. At least 1M cells were resuspended in 100µL of RIPA buffer (ThermoFisher Scientific, 89900) for 30 min on ice to lyse the cells. Lysates were spun for 30 min at 4°C and the supernatants containing the soluble proteins were harvested. Protein concentration was calculated using a Pierce™ BCA Protein Assay Kit (ThermoFisher Scientific, 23227). 20µg of protein was loaded per lane. Samples were mixed with Pierce™ Lane Marker Reducing Sample Buffer (ThermoFisher Scientific, 39000) and run on a NuPAGE™ 3-8% Tris-Acetate Protein Gel with NuPAGE™ Tris-Acetate SDS Running Buffer (ThermoFisher Scientific, LA0041) in a XCell SureLock™ Mini-Cell (ThermoFisher Scientific, EI0001). Transfer onto a Nitrocellulose Membrane, 0.2 µm (BioRad, 1620112) was performed using the XCell SureLock™ Mini-Cell (ThermoFisher Scientific, EI0001) in Pierce™ 10X Western Blot Transfer Buffer, Methanol-free (ThermoFisher Scientific, 35045) for 2 hours at 30V. Membranes were blocked for 2 hours at room temperature with 5% milk in TBST prior to antibody incubation overnight at 4°C. Antibodies were added in 5% milk with TBST. Membranes were washed 6 times 10 min in TBST at room temperature, incubated with HRP secondary antibodies (Cell signaling, 7074) 1:1000 in 5% milk with TBST for 2 hours at room temperature, washed 6 times 10 min with TBST at room temperature, revealed with SuperSignal™ West Dura Extended Duration Substrate (ThermoFisher Scientific, 34076) and analyzed on Biorad ChemiDoc system.

#### Co-Immunoprecipitation (co-IP)

Co-IP protocol was adapted from (Hansen et al., 2019). Cells were grown on 15cm plates, washed with dPBS and harvested with accutase. For each co-IP about 30M cells were used. Each pellet was resuspended in 1mL of low salt lysis buffer (5mM PIPES pH 8.0, 85mM KCl, 0.5% NP-40 and 1X HALT protease inhibitor) and incubated on ice for 10 min. Nuclei were pelleted for 10 min, 4 000 rpm at 4°C and resuspended in 1mL of low salt lysis buffer. Each set of co-IP had 3 samples (non-treated, turbonuclease and RNAseA). The turbonuclease samples were treated with 1 200 units of turbonuclease and the RNAseA samples were treated with 0.1mg/mL of RNAseA. Samples were incubated for 4 hours at 4°C on a rotator. After the incubation a 50µL sample was taken from each tube to check the efficiency of the DNA and RNA degradation. The NaCl concentration of the rest of the samples was adjusted to 200mM and samples were incubated on ice for 30 min. Samples were centrifuged for 10 min, maximum speed at 4°C to extract the protein. Proteins were quantified with BCA and 1mg of protein was used for the co-IP. 1mg of the lysate was precleared for 4 hours at 4°C with 80µL of protein G dynabeads magnetic beads (10004D) washed once in coIP buffer (0.2M NaCl, 25mM HEPES, 1mM MgCl2, 0.2mM EDTA, 0.5% NP-40 and 1X HALT protease inhibitor). After pre-clearing, 1% of input was kept to check CTCF and RAD21 depletion and also to load on the Western blot gels. The 1mL lysate was divided into two tubes of 500µL and incubated overnight either with 5µL of rabbit IgG (Normal Rabbit IgG, #2729, 1mg/mL) or 5µL DDX55 (Bethy,l 1mg/mL, A303-027A) or 15µL TAF5L (Proteintech, 19274-1-AP, 0.333mg/mL) or 5µL TAF6L (ABclonal, A14369, 3.38mg/mL). The next day, 40µL of protein G dynabeads magnetic beads washed once in coIP buffer (10004D) were added to each tube and incubated 2 hours at 4°C. Then, the beads were washed 5 times, 5 min with 500µL of coIP buffer using a magnetic rack at room temperature. Flow Through (FT) and last wash were kept for the Western blot gels. To elute the proteins, the beads were resuspended in 20µL of 2X SDS buffer, heated for 5 min at 100°C and the supernatants were taken after placing the tubes on the magnetic rack. The totality of the 20µL of sample was loaded on a NuPAGE™ Novex™ 3-8% Tris-Acetate Protein Gels, 1.0 mm, 12-well and analyzed by Western blot. Each co-IP was performed in two replicates.

#### DNA and RNA extraction to check DNA and RNA degradation efficiency for co-IP

DNA was extracted with the DNA extraction kit from Qiagen (DNeasy Blood & Tissue Kit, 69504, Qiagen) and resuspended in 25µL of water. DNA concentration was assessed with Qubit broad range kit (Qubit™ dsDNA BR Assay Kit, Q32850, Thermo Fisher Scientific) or nanodrop. 100 ng of the non-treated samples were taken and an equal volume from the nuclease treated samples were taken and used to quantify by qPCR.

RNA was extracted with TRIzol following manufacturer recommendations. After precipitation, RNA was resuspended in 25µL of water. RNA concentration was assessed with nanodrop. For each reverse transcription reaction, 1µg of the non-treated samples were taken and an equal volume for the nucleases treated samples were taken. Reverse transcription was performed with VILO IV (SuperScript™ IV VILO™ Master Mix, 11756050, Thermo Fisher Scientific) and incubated 20 min at 25°C, 10 min at 50°C and 5 min at 85°C. cDNA was diluted by 20 and 2µL was used for the qPCR.

#### qPCR

qPCR was directly done on the cDNAs using Fast SYBR™ Green Master Mix (ThermoFisher Scientific, 4385612) and analyzed on a StepOnePlus™ Real-Time PCR System (ThermoFisher Scientific). See SI item table 1 for qPCR primer sequences.

#### CRISPR sgRNA lentivirus production

TKOv3 library lentivirus was produced by co-transfection of lentiviral vectors psPAX2 (packaging vector, Addgene #12260) and pMD2.G (envelope vector, Addgene #12259) with TKOv3 lentiCRISPRv2 (pLCV2) plasmid library, using X-treme Gene9 transfection reagent (Roche). Briefly, HEK293T cells were seeded at a density of 9×10^6^ cells per 15cm plate and incubated overnight, after which cells were transfected with a mixture of psPAX2 (4.8μg), pMDG.2 (3.2μg), TKOv3 plasmid library (8μg), and X-treme Gene9 (48μL) in Opti-MEM (GIBCO), in accordance with the manufacturer’s protocol. 24 hours after transfection, the medium was changed to serum-free, high BSA growth medium (DMEM with1% BSA (Sigma) and 1% penicillin/streptomycin (GIBCO). Virus-containing medium was harvested 48 hours after transfection, centrifuged at 1500rpm for 5 minutes, and stored at -80°C. Functional titers in cells to be screened were determined by virus titration: 24 hours after infection, the medium was replaced with puromycin-containing medium (1mg/ml), and cells were incubated for 48 hours. The multiplicity of infection (MOI) of the titrated virus was determined 72 hours after infection by comparing the survival of infected cells to infected unselected and non-infected selected control cells.

#### Pooled genome-wide CRISPR screen in HAP1 CTCF degron cell lines

CRISPR TKOv3 screens were performed as previously described (Aregger et al., 2019; Hart et al., 2015, 2017) in the HAP1-CTCFdegron and HAP1-CTCFdegron-TIR1 cells. For HAP1-CTCFdegron, 54M cells were infected and for HAP1-CTCFdegron-TIR1, 110M cells were infected with the TKOv3 lentiviral library at MOI of ∼0.3 for reaching >200-fold coverage of the library after puromycin selection. 24 hours after infection, the infected cells were selected by changing the medium to puromycin-containing medium (1µg/mL). 72 hours after infection was considered as T0. 30M cells were harvested and 135M puromycin-selected cells were seeded in medium without puromycin to keep the >200-fold coverage of the library for each of the three screen replicates and each of the three different conditions (non-treated (NT), 25µM auxin, 500µM auxin). At T3, 20M cells were harvested. The remaining cells were divided into three conditions (non-treated (NT), 25µM auxin, 500µM auxin). Each condition was screened in triplicates with the cells seeded at the appropriate number to keep the 200-fold coverage of the library (i.e. 15M cells per triplicate per condition). Cells were passaged every 3 days and maintained in absence (NT) or in presence of auxin (25µM, 500µM IAA) at 200-fold coverage until reaching T15. For T6, T9, T12 and T15, 20M cells were harvested for each triplicate in each condition. Genomic DNA was extracted for T0, T6 and T15 of each triplicate of each condition using the Wizard Genomic DNA Purification kit (Promega, A1125) following manufacturer’s instructions. Sequencing libraries were prepared by amplifying sgRNA inserts via a 2-step PCR. The first PCR enriches for the sgRNA regions in the genome and the second PCR adds indices by using primers that include Illumina TruSeq adaptors. Briefly, in PCR1 50µg of genomic DNA were amplified in 15 parallel reactions to maintain the 200-fold coverage. DNA was amplified using the NEBNext Ultra II Q5 Master Mix. PCR1 primers were FW1: GAGGGCCTATTTCCCATGATTC, RS1 : GTTGCGAAAAAGAACGTTCACGG. For PCR2, 5µL of the pooled PCR1 product was used and amplified using the NEBNext Ultra II Q5 Master Mix and primers with i5 and i7 indices. PCR2 primers were FW2 : AATGATACGGCGACCACCGAGATCTACACNNNNNNNNACACTCTTTCCCTACACGACGCTCTTCCGATCTTTGTGGAAAGGACGAAACACCG, RS2 : CAAGCAGAAGACGGCATACGAGATNNNNNNNNGTGACTGGAGTTCAGACGTGTGCTCTTCCGATCTACTTGCTATTTCTAGCTCTAAAAC, with NNNNNNNN being the indices of the TruSeq adaptors. After PCR2, the 200bp amplified band was excised from an agarose gel and purified using QIAGEN MinElute Gel Extraction kit (Qiagen, 28604). Libraries were quantified using a Qubit fluorometer and pooled for sequencing. Libraries were sequenced on an Illumina HiSeq2500. Each read was completed with standard primers for dual indexing. The first 21 cycles of sequencing were dark cycles, or base additions without imaging. The actual 26-bp read begins after the dark cycles and contains two index reads, reading the i7 first, followed by i5 sequences. For the T0 libraries we aimed for a total of 30M reads each and for the T6 and T15 libraries we aimed for a total of 20M reads each. The actual number of reads can be found in the supplemental table.

#### ChIP-seq

##### Cell fixation

Cells were seeded in 15cm plates in order to have ∼15M cells per antibody. Cells were fixed in the plates with 1% formaldehyde in HBSS for 10 min at room temperature. To quench the formaldehyde and terminate the cross-linking reaction 125mM glycine was added and incubated for 5 min at room temperature. Plates were then transferred and incubated on ice for 15 min. Formaldehyde solution was then removed and cells were washed with dPBS. Cells were scraped from the plates in dPBS containing protease inhibitors, spun and supernatant was removed.

##### Cell lysis

Each 5M cells were resuspended in 1 mL of hypotonic lysis buffer (20mM TRIS HCl pH8.0, 85mM KCl, 0.5% IGEPAL, 1X HALT protease inhibitor) and incubated 15 min on ice. Cells were then spun down 5 min, 500g, 4°C, supernatant was removed and 5M cells were resuspended in 300µL of sonication buffer (20mM TRIS HCl pH8.0, 0.2% SDS, 0.5% Sodium Deoxycholate, 1X HALT protease inhibitor).

##### Sonication

Cells were sonicated using a bioruptor pico, at 4C, following the manufacturer instructions and with the following settings: 30 s on, 30 s off, for 15 cycles. After sonication, tubes were briefly spun, combined together and spun for 10 min, 12 000g, 4°C. The chromatin fraction is in the supernatant.

##### Assessment of the sonication

To check if sonication was efficient, 10µL of each sample was combined with 200µL of de-crosslinking solution (50mM TRIS HCl, pH 8.0, 200mM NaCl, 0.2mg/mL of proteinase K) incubated overnight at 65°C. The next day de-crosslinked samples were treated with 0.1 mg/mL RNAseA for 15 min at 37°C and DNA was extracted with phenol/chloroform in phase lock tubes (first wash: phenol chloroform, second wash : chloroform). DNA was concentrated using an amicon column and eluted in 10µL of water before loading the totality on a 1.5% agarose gel. Most of the DNA fragments are between 150-400bp.

##### Chromatin dilution and antibody incubation

300µL of chromatin from 5M cells was diluted with 1 200µL of dilution IP buffer (20mM TRIS HCl pH 8.0, 150mM NaCl, 2 mMEDTA, 1% TRITON, 1X HALT protease inhibitor). For input control, 10% of the diluted chromatin was set aside. Antibody was added to each tube (CTCF: 20µL, RAD21: 4µL, DDX55: 4µL, TAF5L: 12µL) and incubated overnight at 4°C on a rotator.

##### Immunoprecipitation

20µL per tube of proteinG Dynabeads were washed in IP buffer 1X (20mM TRIS pH8, 2mM EDTA, 150mM NaCl, 1% TRITON, 0.1% SDS, 1X HALT protease inhibitor), added to the tube and incubated 2 hours at 4°C on a rotator.

To get rid of the non-specific binding, first two 5 min washes with 600µL of IP buffer 1 X were done (20mM TRIS pH8, 2mM EDTA, 150mM NaCl, 1% TRITON, 0.1% SDS, 1X HALT protease inhibitor), second two 5 min washes with 600µL of wash buffer B (20mM Tris HCl, pH 8.0, 500mM NaCl, 2mM EDTA, 0.5% Na Deoxycholate, 1% Triton X-100, 1X HALT protease inhibitor), third, two 5 min washes with 600µL of wash buffer C (20mM Tris HCl, pH 8.0, 1mM EDTA, 0.5% Na Deoxycholate, 250mM LiCl, 1% Triton X-100, 1X HALT protease inhibitor) and finally two 5 min washed with 600µL of TLE (10mM Tris HCl, pH 8.0, 0.1 mM EDTA, 1X HALT protease inhibitor)

##### Elution and DNA purification

DNA was eluted twice by adding 50µL of elution buffer (50mM NaHCO3, 1% SDS) to the beads and incubating 30 min at 65°C. The saved input DNA was diluted in the same elution buffer and treated similarly. RNA was degraded by incubating the samples with 0.1 mg/mL for 30 min at 37°C.

Samples were de-crosslinked by adding 200mM NaCl and 0.2mg/mL of proteinase K overnight at 65°C. The next day, DNA samples were extracted with phenol/chloroform in phase lock tubes (first wash: phenol chloroform, second wash : chloroform). DNA was precipitated with 0.3M Sodium Acetate (pH 5.2), 0.1mg/ml glycogen and 2X volume of ethanol. Samples were incubated on ice for 3 hours to precipitate DNA. DNA was precipitated by spinning for 30 min at 16 000 g at 4°C, washed once with ethanol 70%, spun 10 min at 16 000g, supernatant was removed and DNA pellets were air dried and resuspended in 40µL of water. DNA concentration was assessed by Qubit and the quality of the ChIP-seq library by qPCR.

##### Library preparation

###### End repair

For end repair, 35 μL of sample were transferred to a PCR tube, then 15μL of the end-repair mix (1X NEB ligation buffer (NEB B0202S), 17.5mM dNTP mix, 7.5U T4 DNA polymerase (NEBM0203L), 25U T4 polynucleotide kinase (NEB M0201S), 2.5U Klenow polymerase Polymerase I (NEB M0210L)) were added. The reactions were then incubated at 37°C for 30 minutes, followed by incubation at 75°C for 20 minutes to inactivate Klenow polymerase. An ampure beads purification was then performed (1.6X) and DNA was eluted in 41µL of water.

###### A-tailing

dATP was added to the 3’ ends by adding 9μL of A-tailing mix (5μL NEB buffer 2.1, 1μL of 1mM dATP, 5U Klenow exo (NEB M0212S)) to the 41μL of DNA sample from the previous step. The reaction was incubated in a PCR machine at 37°C for 30 minutes, then at 65°C for 20 minutes. An ampure beads purification was then performed (1.6X) and DNA was eluted in 40µL of water. *Illumina adapter ligation and PCR* The TruSeq DNA LT kit Set A (Illumina, #15041757) was used. 12.5μL of ligation mix (5μL Illumina paired-end adapters (non-diluted if library concentration is more than 100ng, diluted by 15 if library concentration is more than 10ng, diluted by 25 if library concentration is around 5ng), 4μL T4 DNA ligase Invitrogen, 2.5μL 5x T4 DNA ligase buffer (Invitrogen 5X), 1µL of 10mM ATP) was added to the 40μL sample from the previous step. The ligation samples were then incubated at room temperature for 2 hours. An ampure beads purification was then performed (1X) and DNA was eluted in 20µL of water. One PCR reaction (14 cycles) was then performed as follows (20μL DNA, 5μL of Primers mix (TruSeq DNA LT kit Set A 15041757), 20μL Master Mix (TruSeq DNA LT kit Set A 15041757), 5μL of water). An ampure beads purification was then performed (0.64X-1.1X) to select DNA fragments between 200-500bp to remove primers and longer amplified DNA fragments that could inhibit the sequencing reaction and DNA was eluted in 25µL of water. The libraries were sequenced using 50bp paired end reads on a HiSeq4000. Each ChIP-seq was performed in two replicates.

#### RNA-seq

RNA was extracted using RNeasy Mini Kit (Qiagen, 74104) with QIAshredder (Qiagen, 79654). RNA was sent to the BGI Hong Kong facility (DNBseq Eukaryotic Transcriptome resequencing) for library preparation and sequencing DNBseq platform MGISEQ-G400 (100bp paired ends). Each RNA-seq was performed in two replicates.

#### Hi-C

Hi-C was performed as described previously (Belaghzal et al., 2017) with minor modifications.

##### Cell fixation

Cells were fixed in the plates with 1% formaldehyde in HBSS for 10 min at room temperature. 125mM glycine was added to the plate and cells were incubated for 5 min at room temperature to quench the formaldehyde and terminate the cross-linking reaction. Plates were then incubated on ice for 15 min. Formaldehyde solution was then removed and cells were washed with dPBS. Cells were scraped from the plates, spun and supernatant was removed. If Hi-C was not performed immediately after cell fixation, pellets were flash frozen in liquid nitrogen and kept at -80°C until starting the cell lysis.

##### Cell lysis

1-5M formaldehyde cross-linked cells were incubated in 1000μl of cold lysis buffer (10mM Tris-HCl pH8.0, 10mM NaCl, 0.2% (v/v) Igepal CA630, mixed with 10μl of 10X protease inhibitors (Thermofisher 78438)) on ice for 15 minutes. Next, cells were lysed with a Dounce homogenizer and pestle A (Kimble Kontes # 885303-0002) by moving the pestle up and down 30 times, incubating on ice for one minute followed by 30 more strokes with the pestle. The suspensions were centrifuged for 5 minutes at 2,000g at RT using a tabletop centrifuge (Centrifuge 5810R, (Eppendorf)). The supernatants were discarded and the pellets were washed twice with ice cold 500μL 1x NEBuffer 3.1 (NEB). After the second wash, the pellets were resuspended in 720µL of 1x NEBuffer 3.1 and split into two tubes. 18µL were kept at -20°C to assess the chromatin integrity later. Chromatin was solubilized by addition of 38μl 1% SDS per tube and the mixture was resuspended and incubated at 65°C for 10 minutes. Tubes were put on ice and 43μL 10% Triton X-100 was added.

##### Chromatin digestion

Chromatin was digested by adding 400 Units DpnII (NEB) per tube at 37°C for overnight digestion with alternating rocking. Digested chromatin samples were incubated at 65°C for 20 minutes to inactivate the DpnII enzymes, spun shortly and transferred to ice. 10µL were kept at - 20°C to assess the digestion efficiency later.

##### Biotin fill-in

DNA ends were marked with biotin-14-dATP by adding 60μL of biotin fill-in master mix (1X NEB 3.1, 0.25mM dCTP, 0.25mM dGTP, 0.25mM dTTP, 0.25mM biotin-dATP (ThermoFisher#19524016), 50U Klenow polymerase Polymerase I (NEB M0210L)). The samples were incubated at 23°C for 4 hours with agitation and then placed on ice.

##### Blunt end ligation

Ligations were performed by adding 665μL of ligation mix (240μL of 5x ligation buffer (1.8X) (Invitrogen), 120μL 10% Triton X-100, 12μL of 10mg/mL BSA, 50μL T4 DNA ligase (Invitrogen 15224090), and 243μL ultrapure distilled water (Invitrogen)). The reactions were then incubated at 16°C for 4 hours with some agitation. After the ligation, the crosslink was reversed by adding 50µL of 10mg/mL proteinase K (Fisher BP1750I-400) and incubated at 65°C for 2 hours followed by a second addition of 50μL 10 mg/mL Proteinase K and overnight incubation at 65°C.

##### DNA purification

Reactions were cooled to room temperature and the DNA was extracted by adding an equal volume of saturated phenol pH 8.0: chloroform (1:1) (Fisher BP1750I-400), vortexing for 1 minute, transferred to a phase-lock tube and spun at 16,000g for 5 minutes. DNA was precipitated by adding 1/10th of 3 M sodium acetate pH 5.2, 2 volumes of ice-cold 100% ethanol and incubated for at least an hour at -80°C. Next, the DNA was pelleted at 16,000 g at 4°C for 30 minutes. The supernatants were discarded, the pellets were dissolved in 500μL 1X TLE and transferred to a 15mL AMICON Ultra Centrifuge filter (UFC903024 EMD Millipore). 10mL of TLE was added to wash the sample, the columns were spun at 4,000 g for 10 minutes and the flow-throughs were discarded. A second wash with 10mL of TLE was done and the sample was transferred to a 0.5mL AMICON Ultra Centrifuge filter (UFC5030BK EMD Millipore) and spun at 16,000g for 10 minutes to reduce the sample to 50µL. RNA was degraded by adding 1μL of 10 mg/mL RNAase A and incubated at 37°C for 30 minutes. DNA was quantified by loading on a 1% gel 1µL of the Hi-C sample, the chromatin integrity and the digestion controls.

##### Biotin removal from unligated ends

To remove biotinylated nucleotides at DNA ends that did not ligate, the Hi-C samples were treated with T4 DNA polymerase. 5μg of Hi-C library were incubated with 5μL 10x NEBuffer 3.1, 0.025mM dATP, 0.025mM dGTP and 15U T4 DNA polymerase (NEB # M0203L) in 50µL. Reactions were incubated at 20°C for 4 hours, the enzymes were then inactivated at 75°C for 20 minutes and placed at 4°C.

##### DNA shearing

The samples were pooled and the volume was brought up to 130μL 1X TLE. The DNA was sheared to a size of 100-300 bp using a Covaris instrument (Duty Factor 20%, Cycles per Burst 200, peak power 50, average power 17.5 and process time 180 sec). The volume was brought up to 500μL with TLE for Ampure fractionation. To enrich for DNA fragments of 100-300bp an Ampure XP fractionation was performed (Beckman Coulter, A63881) and the DNA was eluted with 50µL of water. The size range of the DNA fragments after fractionation was checked by running an aliquot on a 2% agarose gel.

##### End repair

To proceed for end repair, 45μL of Hi-C sample was transferred to a PCR tube, then 25μL of the end-repair mix (3.5X NEB ligation buffer (NEB B0202S), 17.5mM dNTP mix, 7.5U T4 DNA polymerase (NEBM0203L), 25U T4 polynucleotide kinase (NEB M0201S), 2.5U Klenow polymerase Polymerase I (NEB M0210L)) was added. The reactions were then incubated at 37°C for 30 minutes, followed by incubation at 75°C for 20 minutes to inactivate Klenow polymerase.

##### Biotin pull down

To pull down biotinylated DNA fragments, 50μL of MyOne streptavidin C1 beads mix (Thermo Fisher 65001) was transferred to a 1.5mL tube. The beads were washed twice by adding 400μL of TWB (5mM Tris-HCl pH8.0, 0.5mM EDTA, 1M NaCl, 0.05% Tween20) followed by incubation for 3 minutes at RT. After the washes, the beads were resuspended in 400μL of 2X Binding Buffer (BB) (10mM Tris-HCl pH8, 1mM EDTA, 2M NaCl) and mixed with the 400μL DNA from the previous step in a new 1.5mL tube. The mixtures were incubated for 15 minutes at RT with rotation. The DNA bound to the beads was washed first with 400μL of 1X BB and then with 100μL of NEB2.1 1X. Finally, beads with bound DNA were resuspended in 41μL of NEB2.1 1X. *A-tailing* Then, dATP was added to the 3’ ends by adding 9μL of A-tailing mix (5μL NEB buffer 2.1, 5μL of 1mM dATP, 3U Klenow exo (NEB M0212S)) to the 41μL of beads with bound DNA from the previous step. The reaction was incubated in a PCR machine (at 37°C for 30 minutes, then at 65°C for 20 minutes, followed by cooling down to 4°C). Next, the tube was placed on ice immediately. The beads with bound DNA were washed twice using 100μL 1X T4 DNA Ligase Buffer (Invitrogen). Finally, beads with bound DNA were resuspended in 36.25μL 1X T4 DNA Ligase buffer (Invitrogen).

##### Illumina adapter ligation and PCR

The TruSeq DNA LT kit Set A (Illumina, #15041757) was used. 10μL of ligation mix (3μL Illumina paired-end adapters, 4μL T4 DNA ligase Invitrogen, 2.75μL 5x T4 DNA ligase buffer (Invitrogen 5X)) was added to the 36.25μL beads with bound DNA from the previous step. The ligation samples were then incubated at room temperature for 2 hours on a rotator. The beads with bound DNA were washed twice with 400μl of TWB, then twice using 100μL NEB2.1 1X. Finally, the samples were resuspended in 20μL of NEB2.1 1X. Two trial PCR reactions (6 and 8 cycles) were performed as follows (0.9μL DNA bound to beads, 1.5 μL of Primers mix (TruSeq DNA LT kit Set A 15041757), 6µL Master Mix (TruSeq DNA LT kit Set A 15041757), 6.6µL of ultrapure distilled water (Invitrogen)). The number of PCR cycles to generate the final Hi-C material for deep sequencing was chosen based on the minimum number of PCR cycles in the PCR titration that was needed to obtain enough DNA for sequencing. ClaI digestion was done as a library quality check. A downward shift of the amplified DNA to smaller sizes indicates that DNA ends were correctly filled in and ligated (creating a ClaI site). Primers were removed using Ampure XP beads. The libraries were sequenced using 50bp paired end reads on a HiSeq4000. Each Hi-C was performed in two replicates.

### QUANTIFICATION AND STATISTICAL ANALYSIS

#### CRISPR screen analysis

Library processing was performed as in (Mair et al., 2019). Reads were trimmed by removing up to 20bp after the first 8bp of the anchors used in the barcoding primers. Reads were aligned using BOWTIE version 0.12.8 (allowing for max. 2 mismatches, ignoring qualities) (Langmead et al., 2009). Read counts for each screen library were normalized to 10M reads total per sample (see SI item table S2 containing the CRISPR screen statistics). Fold change was calculated to the T0 reference sample. For precision and recall plots, we generated Bayes Factor (BF) scores with the calculated fold changes using BAGEL version 0.91 (Hart and Moffat, 2016). We then calculated precision and recall using these BFs, discarding scores for genes that were not included in version 2 training sets (union of essential (EG) and non-essential gene (non-EG) sets from https://github.com/hart-lab/bagel, 684 EGs and 926 non-EGs).The essential set was used as the true positive list for the precision_recall_curve function of the Scikit-learn library for Python, along with the above BF score subset (Pedregosa et al., 2011). For generating fold change plots of EGs and non-EGs, we calculated the mean fold change of all guides for a gene for each screen. Replicates were grouped together and mean fold change for each gene was calculated. The x axis was defined as [min(fold_change), max(fold_change)] on a 0.2 interval. Gaussian kernel density estimation of fold changes was performed separately for essential and non-essential training sets with kde.factor 0.25. using gaussian_kde from Scipy Python library (Virtanen et al., 2020). Essential and non-essential data were plotted separately for each grouped set of replicates.

##### Gene hit selection

To select genes that affect cell growth in the context of CTCF depletion, we performed three steps.

First, we compared the log2 fold changes (log2 FC), calculated to the T0 reference sample, of the HAP1-CTCFdegron screens against the HAP1-CTCFdegron-TIR1 screens for each time point (T6 and T15) and condition (NT, 25µM and 500µM auxin (IAA)) for a total of six comparisons. We then calculated the difference between the log2 FC HAP1-CTCFdegron-TIR1 with log2 FC HAP1-CTCFdegron for each condition and time point (log2 FC HAP1-CTCFdegron-TIR1 - log2 FC NO HAP1-CTCFdegron). We selected genes that had a log2 FC difference > |1|, i.e., a fold change > |2| for each time point and condition. We calculated the log2 FC difference separately for positive and negative values (i.e., < -1 or > 1). A positive value indicates that the cells were growing better when CTCF was depleted and the target gene was knocked out; we call them positive interaction hits. A negative value indicates that cell growth was affected when CTCF was depleted and the target gene was knocked out; we call them negative interaction hits. This selection resulted in hit lists for positive and negative interactions for each time point and condition. We then merged these gene lists, keeping positive or negative interaction hits separated. We next wanted to remove from this analysis the interactions caused by auxin only. Therefore, we performed the same analysis as described above for comparing NT versus 25µM or 500µM auxin only in the HAP1-CTCFdegron cells (log2 FC HAP1-CTCFdegron auxin treated (25µM or 500µM) - log 2 FC HAP1-CTCFdegron NT).

The “hits” obtained this way were labeled auxin specific genes and removed from the overall hit lists.

Second, independently of step one, we calculated the ratio of the FC auxin-treated by the FC NT (log2(FC treated with auxin (25 or 500µM auxin) / FC NT).

We compared these ratios from the HAP1-CTCFdegron screens against the HAP1-CTCFdegron-TIR1 screens for each time point (T6 and T15) and condition (25µM and 500µM auxin) for a total of four comparisons. We then calculated the difference between the ratio of HAP1-CTCFdegron-TIR1 with the ratio of HAP1-CTCFdegron for each condition and time point (ratio HAP1-CTCFdegron-TIR1 - ratio HAP1-CTCFdegron).

We selected genes that had a difference > |1|, ie a fold change > |2| for each time point and condition and merged the hit lists as described above, again keeping positive and negative interactions separate. We again removed the auxin specific genes from these lists.

Third, we calculated the union of the gene hits from step 1 and step 2, separately for positive and negative interactions. The final gene list had 469 negative interaction hits and 294 positive interaction hits.

##### GO analysis

Functional enrichment in the gene hit lists was calculated using Enrichr, GO Biological Process 2018 (Kuleshov et al., 2016).

##### CTCF concentration time course

sgRNA depletion or enrichment was assessed over decreasing CTCF levels (ranging from full amount of CTCF to nearly no CTCF. 0, HAP1-CTCFdegron, full amount of CTCF; 1, HAP1-CTCFdegron-TIR1 NT, decreased level of CTCF; 2, HAP1-CTCFdegron-TIR1 auxin 25µM, further decrease of CTCF level; 3, HAP1-CTCFdegron-TIR1 auxin 500µM, nearly no CTCF left). We calculated the log2 fold change for each gene for each CTCF concentration to its respective T0 sample. Log2 fold changes were then subtracted with the respective log2 fold changes in the HAP1-CTCFdegron samples and the resulting matrices were plotted as heatmaps. Genes that change (indicated as red in Figure 4) have a difference of log2 fold change of > |0.4|. Genes with similar patterns were manually grouped together.

#### RNA-seq analysis

##### Bulk RNA-seq analysis

RNA-seq data was mapped using a standardized pipeline in the DolphinNext environment (Yukselen et al., 2020). Briefly, reads were mapped to hg19 using STAR (Dobin et al., 2013) with default parameters to generate BAM files. RSEM was used for transcript quantification (Li and Dewey, 2011). To identify Differentially Expressed genes, DESeq2 was run using the DEBrowser interface (Kucukural et al., 2019) (see SI item table S3 containing the RNA-seq statistics). Genes that had less than 10 counts per million were removed using Low Count Filtering method. The dispersion was estimated using a parametric fit and hypothesis testing performed using the likelihood ratio test. We selected differentially expressed genes that were at least 1.25-fold up- and down-regulated, with p-values less than 0.05.

##### Splicing RNA-seq analysis

For alternative splicing (AS) analyses, reads were trimmed with Trimmomatic (0.32), aligned with STAR (2.70e) and AS events detected with RMATS (4.1.0) (Park et al., 2013; Shen et al., 2012, 2014). AS events with FDR<0.05 were considered as statistically significant AS events. DDX55 knock-out clones in absence of auxin (NT), DDX55 knock-out clones in presence of auxin (IAA), TAF5L knock-out clones in absence of auxin (NT), TAF5L knock-out clones in presence of auxin (IAA) were pooled as technical replicates for RMATS analyses.

#### ChIP-seq analysis

##### Raw data processing

50bp paired ends reads were processed using the nf-core/chipseq pipeline, version 1.1.0: https://github.com/nf-core/chipseq (Ewels et al., 2020). Briefly, fastq files were mapped to the hg19 reference genome using BWA, mapped reads were filtered to remove duplicates, unmapped reads, multi-mappers, etc, and finally bigWig files scaled to 1 million mapped reads were created. ChIP-seq samples passed QC-tests (see SI item table S4 containing the ChIP-seq statistics).

##### Peak calling

We used MACS2 to find the enriched ChIP-seq peaks with the following parameters: macs2 callpeak -q 0.01 (Zhang et al., 2008). To have a final list of ChIP-seq peaks for each individual protein (CTCF, RAD21, DDX55 and TAF5L), we took the union of the peaks called in the two ChIP-seq replicates. Peaks that were as close as 10bp or overlapping were merged together using bedtools merge -d 10. Intersection between the peaks was done using bedtools intersect (Quinlan and Hall, 2010).

##### Motif search analysis

We used MEME to find motifs in sequences corresponding to ChIP-seq peaks and intersection of ChIP-seq peaks. Following parameters were used: -mod anr -nmotifs 3 -minw 15 -maxw 50 - objfun classic -revcomp -markov_order 0 (Bailey and Elkan, 1994; Bailey et al., 2009).

##### Peak annotation

Peaks were annotated using annotatePeaks.pl from HOMER/4.6 (Heinz et al., 2010).

##### Venn diagrams

Venn diagrams were generated using intervene venn and bed files (Khan and Mathelier, 2017).

#### Aggregation stackups

Series of stackup panels were generated using custom scripts. Each panel is demonstrating behavior of a number of *signals* - in columns (e.g., ChIP-seq, insulation, EV1, RNA-seq etc.) around a set of genomic loci of interest, *features* - in rows (e.g., ChIP-seq peaks, genes TSS/TTS, subsets of the above, etc.). In a given column, *signals* are extracted using bbi.stackup function from Python API for UCSC BBI library (https://github.com/nvictus/pybbi) (Kent et al., 2010) centered on a given feature with a given flank. Every column in a given stackup panel is sorted synchronously, and a “summary” (average) profile of every *signal* is presented in the axes above.

We used following *signal*-specific parameters for stackups (unless stated otherwise in a figure caption):

- all ChIP-seq, RNA-seq, G4-seq, DRIP-seq derived signals used 200 bp-sized bins with a 5kb flank, R-loops signal used 10kb flank instead.

- Insulation signal for *20*kb *- diamond* used 2kb - sized bins and flank of 40kb, whereas insulation with 100kb *- diamond* used 5kb - sized bins and 200kb flanks.

- EV1 signal used 25kb - sized bins and 150kb - sized flanks.

When specified on a figure caption, signals extracted for gene-related features (TSS/TTS) were “flipped” for genes on the negative strand, to ensure that the body of a given gene is on the right side of the stackup for TSS, and on the left side of the stackup for TTS.

Stackups were sorted according to the signal specified in a figure caption using average of the central bins, with the exception of stackups sorted on R-loops and stackups sorted on EV1 signal: average of an entire row was used for sorting R-loops signal at TTS, due to the wide nature of the signal; “drop” of EV1 signal, calculated as a difference of average EV1 between the left and the right flank, was used for sorting stackups using EV1 - this allowed us to reveal apparent B to A and A to B transitions near the center of each feature.

Linear color scales were used throughout the stackups with the following exceptions:

- RNA-seq signal was plotted using log-spaced colormap and the corresponding average profile is a geometric mean of RNA-seq signal

- discrete signals extracted from bigBED files: G4-seq and R-loop used blue color to indicate footprints of peaks and white color to indicate regions without any peaks; gene annotation used red/blue colors to indicate genes on the positive/negative strands and grey color to indicate regions without annotated transcripts.

To plot the distribution of CTCF peaks/motifs, RAD21 peaks in presence of CTCF and G4-seq, we calculated the rolling sum of those peaks with the window size 100 and plotted the sums for the non-overlapping windows along the stackup.

Insulation signal was subject to additional normalization before plotting on the stackup - we used average insulation signal of the outermost half-flank upstream and downstream to normalize each individual insulation profile on the stackup. This normalization was done to alleviate fluctuations of insulation signal along the chromosomes (e.g. near-telomeric regions, compartmental variation, etc.).

#### List of features used in the paper

##### List of biologically relevant peaks used in Figure 1

We have assembled a list of biologically relevant peaks, by combining CTCF and RAD21-peaks: we merged together all the CTCF peaks in presence of CTCF (NT) and all the RAD21 peaks in presence and absence of CTCF (NT and IAA). The final list contains 39 233 peaks.

##### CTCF-only, CTCF and G4, G4-only (for Figure 2)

To create the three categories for Figure 2, the CTCF ChIP seq peaks list was extended by 2.5kb in each direction of the CTCF summit using bedtools slop (bedtools slop -b 2500) and genomic segments separated by less than 10bp were then merged using bedtools merge (bedtools merge -d 10). The G4 list from Mao et al was extended by 2kb in each direction of the CTCF peak using bedtools slop (bedtools slop -b 2000) and genomic segments separated by less than 10bp were then merged using bedtools merge (bedtools merge -d 10). The two lists were then intersected (bedtools intersect -u) or the locations of one list that did not intersect with the other list were also taken (bedtools intersect -v). The final list contains 23 871 CTCF-only sites, 4 020 CTCF and G4 sites and 4 925 G4-only sites.

##### List of genes (Gene annotation)

We downloaded gene annotation RefGene for hg19 human reference genome from UCSC and matched it with gene description gene_info from NCBI using gene2refseq dictionary. Genes with identical names and TSS coordinates were merged together to create longest possible transcripts using exonU.py script from clodius (https://github.com/higlass/clodius). Resulting gene annotation was used to create gene-related feature lists in the manuscript.

##### Active TSS in HAP1 and K562 cells

We used publicly available active mark H3K4me3 ChIP-seq signal to rank annotated TSS from the most active to inactive (Campagne et al., 2019; Mchaourab et al., 2018). Top 13 412 TSSs were selected as Active TSSs in HAP1 (H3K4me3 signal drops to nearly 0 after the top 13 412 TSSs).

We further filtered out active TSS that were too close to a called CTCF peak (<2kb) (in HAP1), in order to avoid interfering signals from CTCF-insulation and CTCF-related interactions. The final list contains 10 933 active TSS without CTCF in HAP1.

##### Active TSS shared in HAP1 and HCT116 cells

First, we generated a list of Active TSS for HCT116 cell line following the same procedure as for HAP1 cell line, using publicly available active mark H3K4me3 for HCT116 (ENCODE, HCT116). Intersection of this list with the list of active TSS in HAP1 yielded 12 113 TSS that are active in both cell lines. To avoid interfering CTCF insulation and interactions in the analyses, we further filtered our active TSS that have CTCF by doing the union of all the CTCF peaks in HAP1 and HCT116 cell lines and intersecting it with the common list of active TSSs. The final list contains 6 802 common active TSS without CTCF.

##### Active TTS in HAP1 cells

We selected active TTSs by taking the list of active TSS in HAP1 and attributing for each active TSS its TTS. To avoid having interfering TSS insulation in the TTS stackups, we removed TTSs that have H3K4me3 nearby. To avoid having interfering CTCF insulation in the TTS stackups, we intersected the active TSSs without H3K4me3 (extended 2kb each side) with the CTCF peaks using bioframe (Abdennur et al., 2020) . The final list contains 10 933 active TTSs without CTCF.

##### Active TTS shared in HAP1 and HCT116 cells

To create a common list of active TTSs without H3K4me3 between HAP1 and HCT116 cell lines, the intersection between the active TTSs without H3K4me3 in HAP1 and the active TTSs in HCT116 was done. To avoid having interfering CTCF insulation in the TTS without H3K4me3 stackups, we removed the active TTSs without H3K4me3 that have CTCF by doing the union of all the CTCF peaks in HAP1 and HCT116 cell lines and intersecting it with the common list of active TSSs without H3K4me3. The final list contains 8 894 common active TTSs without CTCF.

##### Consensus list of R-loops

We created a consensus list of R-loops by intersecting different DRIP-seq datasets published in (Sanz et al., 2016). We intersected the NT2 and K562 datasets (extended 2kb each side) by using bedtools intersect. The final list contains 19 229 R-loops.

##### Enhancers

We created a list of HAP1 enhancers by overlapping H3K27ac peaks with HAP1 DNase-seq signals (ENCODE, HAP1). We filtered out enhancers that were too close to a called CTCF peak (<2kb), in order to avoid interfering signals from CTCF-insulation and CTCF-related interactions. We further filtered out TSS that were too close to an active TSS (H3K4me3 active mark) (<2kb), in order to avoid interfering signal from TSS insulation. The final list contains 10443 enhancers.

##### CTCF motif orientation

CTCF peaks were orientated using one CTCF motif in Jaspar (MA0139.1). If a CTCF peak had multiple CTCF motifs, the motif with the strongest p-value was attributed to the CTCF peak.

##### G4 motif orientation

G4 ChIP-seq peaks were orientated using a list of orientated G4 from G4-seq (Chambers et al., 2015; Mao et al., 2018). If a G4 peak had multiple G4 motifs, only peaks that had motifs with the same orientation were kept.

#### Hi-C analysis

##### Hi-C data processing

Hi-C libraries were processed using the distiller pipeline (https://github.com/open2c/distiller-nf). Briefly, paired-end reads were mapped to the human reference assembly hg19 using bwa mem in a single-sided fashion (-SP). Then the pairtools package (https://github.com/mirnylab/pairtools) was used to parse alignments and classify ligation products, or pairs. Uniquely mapped and rescued pairs were kept after removal of duplicates.

Pairs were further filtered, keeping only those with high mapping quality scores on both sides (MAPQ > 30), and subsequently aggregated into binned contact matrices in the cooler format (Abdennur and Mirny, 2020) at 1, 2, 5, 10, 25, 50, 100, 250, 500 and 1000kb resolutions. All contact matrices were normalized using the iterative correction normalization (Imakaev et al., 2012) after excluding the first 2 diagonals to avoid short-range ligation artifacts at a given resolution. Bins with extreme coverage were excluded using the MADmax (maximum allowed median absolute deviation) filter on genomic coverage, described in (Schwarzer et al., 2017), using the default parameters (see SI item table S5 containing the Hi-C statistics). Sex chromosomes (chrX, chrY) and mitochondral genome (chrM) were left out from Hi-C analysis. The data was initially inspected using HiGlass (Kerpedjiev et al., 2018). For the figures, Hi-C interaction heatmaps were plotted using matplotlib (Hunter, 2007).

##### Scaling plots

We used balanced Hi-C data binned at 1kb to calculate average frequency of interactions P(s) between pairs of loci separated by the same genomic distance (s) independently for each chromosomal arm, using compute_expected from cooltools (Venev et al., 2021). As Hi-C data becomes very sparse at large genomic separations, average frequencies were grouped into log-spaced bins of genomic separation and averaged using logbin_expected from cooltools. These log-binned distance decay curves for each chromosomal arm were combined together to summarize distance decay genome-wide and calculate the rate at which interactions decay with distance, i.e., scaling plot derivatives, using combine_binned_expected from cooltools.

Average distance decay P(s) calculated using compute_expected from cooltools, at a given resolution was also used as an expected (to calculate observed-over-expected signal, OE) for a number of subsequent analyses including eigendecomposition, interaction pileups and dot pileups.

##### Compartments

A and B compartments were assigned using an eigenvector decomposition procedure (Imakaev et al., 2012) implemented in the cooltools package (Venev et al., 2021). Eigenvector decomposition was performed on observed-over-expected cis contact matrices for every chromosome arm binned at 25kb and 100kb. The first eigenvectors (EV1) positively correlated with the gene density were used to assign A and B compartment identity to each bin.

The strength of compartmentalization was analysed using “saddle”-plots as implemented in cooltools (Imakaev et al., 2012). Briefly, EV1 values were digitized into 50 quantiles (excluding 2% of outlier values) EV1_i_ i=1,2,..,50, and observed-over-expected (OE) contact signal was averaged across chromosomal arms for every combination of quantiles EV1_i_ -EV1_j_.

We estimated the strength of A compartment as an enrichment of AA interactions over AB: AA / ((AB+BA)/2), where AA is an average of OE interactions between EV1 quantiles with 20% strongest A-compartment identity, and AB(=BA) is an average of OE interactions between quantiles with 20% strongest A- and B-identities. Similarly for B-compartment, strength was estimated as: BB / ((AB+BA)/2).

##### Insulation

Insulation scores were calculated using cooltools implementation of diamond insulation method (Crane et al., 2015). Insulation at TSS was calculated using a diamond size of 20kb and a bin size of 2kb, unless specified otherwise in the text and/or figure caption. Insulation at TTS was calculated using a diamond size of 100kb and a bin size of 5kb.

##### Contact frequency pileup heatmaps

In order to analyze the average interaction pattern at a number of selected locations (on-diagonal pileups) and for a number of selected pairwise features (off-diagonal pileups) we used snipping module from cooltools. Briefly, individual slices of observed-over-expected interaction heatmap centered on selected features and extended for the specified genomic distances were extracted and averaged.

##### Dot selection

Dots in the HAP1 cell line have been characterized in (Sanborn et al., 2015). The CTCF degron was built in the HAP1 cell line however the CTCF degron is now different from the HAP1 parental cell line. As described in the result part, the CTCF degron has also a lower level of CTCF. Some dots that were present in the parental HAP1 cell line are not present in the CTCF degron. In order to only select the dots that are present in the CTCF degron cell line, we overlap the Sanborn dot anchors with the CTCF degron ChIP-seq peaks (strength > 150). We only selected the dot anchors that had a CTCF peak. We then only selected the dots that had a CTCF peak on either anchor. The final list has 4661 dots. For the dots that had remaining CTCF after CTCF depletion, we took the CTCF degron list of dots and selected the dots that had remaining CTCF after CTCF depletion (CTCF ChIP-seq after auxin treatment) on either dot anchor. There are 1 604 potential dots that have remaining CTCF after CTCF depletion but they are not able to form interactions.

##### All pairwise interaction pileups

###### CTCF-TSS

All pairwise combinations between CTCF-only peaks with RAD21 and active TSS without CTCF separated by 50-500kb were generated using pair_by_distance from bioframe.

###### CTCF-TSS nearest

Closest CTCF or TSS were identified using closest from bioframe independently for upstream or downstream CTCF sites. The two lists were then concatenated and CTCF and TSS separated by 50-500kb were kept.

###### Promoter-enhancer

All pairwise combinations between active TSS without CTCF and enhancer without CTCF separated by 50-500kb were generated using pair_by_distance from bioframe.

###### Gene and enhancer-promoter meta-analysis

Demonstration of insulation profiles and interaction landscapes around genes was performed separately for *active* and *inactive* gene categories. *Active* genes were defined as genes with expression levels >= 5 TPM across all generated RNA-seq samples; *Inactive* - genes with expression levels = 0 TPM across all generated RNA-seq samples. We considered genes only with transcript sizes >= 50kb for both categories, as it is difficult to interpret Hi-C data for genes smaller than 10-bins in size (5kb bin size).

##### Rescaled pileups for genes and enhancer-promoter interactions

We used the coolpuppy package (Flyamer et al., 2020) to illustrate average interaction landscape around select genes and enhancer-promoter (EP) pairs. On diagonal interaction-pileups were generated using genomic intervals between TSS and TTS as genes and genomic intervals between enhancers and TSS as EP interaction pairs.

For both types of genomic regions, rescaled pileups were generated as follows: regions of interest were extended 100% up- and down-stream (flanking) and corresponding Hi-C matrices were extracted at 5kb resolution. Extracted matrices were normalized by the randomly shifted local controls, intelligently rescaled to 200*200 pixels using zoom_array from cooltools, and finally averaged for the regions of interest.

Rescaled pileups were generated independently for positively- and negatively-stranded genes. Pileups for negatively stranded genes were then “flipped” to match the orientation of positively stranded genes and combined with the pileups of the latter ones. Similar transformation was done to combine matching pileups of promoter-enhancer regions: upstream enhancer - (positive strand)TSS combined with the downstream enhancer - (negative strand)TSS; upstream enhancer - (negative strand)TSS combined with the downstream enhancer - (positive strand)TSS, etc.

##### Metagene analysis

We further excluded genes that have detectable levels of CTCF ChIP-seq signal in a 2kb vicinity of TSS or TTS, in order to avoid any insulation or interaction patterns associated with CTCF. Genomic intervals between TSS and TTS were extended by 100%, and then used to extract a profile of the signal (insulation, R-loop peaks, etc) from corresponding bigWig files. We used bbi.fetch with 3000 bins to extract profiles “rescaled” to the same number of bins regardless of the interval size (Kent et al., 2010). Extracted profiles were further averaged and in the case of insulation - normalized by subtracting the mean value of the outermost half of both flanks.

Jupyter notebooks used in this study can be found on GitHub

## SUPPLEMENTAL ITEMS

Table1. Primers used in this study related to Methods

Table2. Genome wide CRISPR screen statistics related to Methods

Table3. RNA-seq statistics related to Methods

Table4. ChIP-seq statistics related to Methods

Table5. Hi-C statistics related to Methods

## SUPPLEMENTAL FIGURE LEGENDS

Figure S1. CTCF depletion weakens insulation at TAD boundaries and has minor effects on compartment structures, related to Figure 1

(A) Schematic representing the strategy used to construct the HAP1-CTCFdegron-TIR1 cells.

(B) Schematic representing CTCF depletion in asynchronous HAP1-CTCFdegron-TIR1 cells (top). Western blot against CTCF in HAP1-CTCFdegron cells and in HAP1-CTCFdegron-TIR1 cells in absence and presence of auxin (NT and IAA). Ponceau is shown for loading control (bottom). Notice that the HAP1-CTCFdegron-TIR1 cells in absence of auxin (NT) have a lower level of CTCF compared to the HAP1-CTCFdegron cells.

(C) Flow cytometry for GFP in HAP1-CTCFdegron and HAP1-CTCFdegron-TIR1 cells in absence and presence of auxin (NT and IAA). Notice that the HAP1-CTCFdegron-TIR1 cells in absence of auxin (NT) have a lower level of CTCF compared to the HAP1-CTCFdegron cells.

(D) Flow cytometry for asynchronous HAP1-CTCFdegron-TIR1 cells in absence and presence of auxin (NT and IAA) stained with Propidium Iodide (PI) to assess the DNA content for cell cycle analysis.

(E) Schematic representing the strategy used to sort the HAP1-CTCFdegron-TIR1 cells in G1 in absence and presence of auxin (NT and IAA).

(F) FACS plot representing the DNA content (PI) relative to GFP for the G1 sorted HAP1-CTCFdegron-TIR1 cells in absence (GFP+, top) or presence (GFP-, bottom) of auxin.

(G) Hi-C contact frequency as a function of genomic distance, P(s) (top) and calculated derivative of P(s): dP/ds (bottom) for HAP1-CTCFdegron-TIR1 cells sorted in G1 phase in absence and presence of auxin (NT and IAA). As a reference the P(s) and its derivative for the asynchronous HAP1-CTCFdegron-TIR1 cells are also plotted.

(H) Hi-C contact frequency as a function of genomic distance, P(s) (top) and its derivative dP/ds (bottom) for asynchronous HAP1-CTCFdegron and HAP1-CTCFdegron-TIR1 cells in absence and presence of auxin (NT and IAA).

(I) Hi-C contact heatmaps at 250kb resolution with the corresponding track of the first Eigenvector (EV1) across chromosome 15 for asynchronous HAP1-CTCFdegron and HAP1-CTCFdegron-TIR1 cells in absence and presence of auxin (NT and IAA).

(J) Genome-wide saddle plots of Hi-C data binned at 100kb resolution for asynchronous HAP1-CTCFdegron and HAP1-CTCFdegron-TIR1 cells in absence and presence of auxin (NT and IAA). The numbers in the corners of the saddle plots indicate compartment strengths for A and B compartments.

(K) Effect of CTCF titration on TADs (left) and dots (right). Hi-C contact heatmaps at 50kb resolution for a 7Mb region on chromosome 10 for asynchronous HAP1-CTCFdegron and HAP1-CTCFdegron-TIR1 cells in absence and presence of auxin (NT and IAA). The corresponding differential interaction heatmaps are shown on the right. Dot pileups for dots characterized in HAP1 cells (Sanborn et al., 2015) that have a CTCF peak in either anchor in the Non-Treated sample (NT) (4 496 dots) and that have a CTCF peak in both anchors in the auxin sample (IAA) (1 545 dots) for asynchronous HAP1-CTCFdegron and HAP1-CTCFdegron-TIR1 cells in absence and presence of auxin (NT and IAA).The dots were aggregated at the center of a 100kb window at 2kb resolution. Notice that the HAP1-CTCFdegron-TIR1 cells in absence of auxin that have a lower level of CTCF have weaker TAD boundaries and dots.

(L) Stackups for the two categories identified in Figure 1 (insulation CTCF dependent (I, blue) and insulation CTCF independent (III, orange)), sorted on the change of the first Eigenvector (EV1) from left to right flank. CTCF and RAD21 ChIP-seq, calculated insulation and EV1 values in HAP1-CTCFdegron-TIR1 cells in the absence and presence of auxin (NT and IAA) were plotted along with the published K562 G4 ChIP-seq.

Figure S2. Insulation at G4 is associated with TSS but not with compartment boundaries, related to Figure 2

(A) Stackups for active TSS, sorted on K562 G4 ChIP-seq signal. CTCF and RAD21 ChIP-seq, calculated insulation and RNAseq in the HAP1-CTCFdegron-TIR1 cells in the absence and presence of auxin (NT and IAA) were plotted along with the published K562 G4 ChIP-seq, G4-seq, RNA polII ChIP-seq, K562 and HAP1 H3K4me3 ChIP-seq and genes. Genes on the forward strand are represented in red (plus) and genes on the reverse strand are represented in blue (minus), grey color corresponds to loci without annotated transcripts.

(B) Stackups for K562 G4 ChIP-seq, sorted on HAP1 H3K4me3 ChIP-seq signal. CTCF and RAD21 ChIP-seq, calculated insulation and RNAseq in the HAP1-CTCFdegron-TIR1 cells in the absence and presence of auxin (NT and IAA) were plotted along with the published K562 G4 ChIP-seq, G4-seq, RNA polII ChIP-seq, K562 and HAP1 H3K4me3 ChIP-seq and genes. Genes on the forward strand are represented in red (plus) and genes on the reverse strand are represented in blue (minus), grey color corresponds to loci without annotated transcripts.

(C) Dot pileup aggregation plots with control orientations for Figure 2, in absence and presence of auxin (NT and IAA), for all the pairwise combinations of each category represented in Figure 2A (I, CTCF-only, II, CTCF and G4, III, G4-only) separated by 50-500kb for a 100kb window at 2kb resolution. The black arrows represent the CTCF motif and the direction of the arrow, the motif orientation. The black squares represent the G4 and the location of the black square, the G4 orientation (right, plus strand) (left). CTCF (upstream)-TSS pairwise interactions and CTCF (downstream)-TSS pairwise interactions are plotted. A quantification of the aggregation pileups is plotted at the CTCF site (mean of the 5 central bins at the CTCF site) (middle). CTCF (upstream)-TSS pairwise interactions and CTCF (downstream)-TSS pairwise interactions are plotted without any CTCF peaks or TSS in between them. A quantification of the aggregation pileups is plotted at the CTCF site (mean of the 5 central bins at the CTCF site) (nearest, right). The black arrows represent the CTCF motif and the direction of the arrow, the motif orientation. The double arrows represent the TSS and the directions of the arrow, the TSS orientation.

(D) Stackups for two categories plotted in Figure 2A (CTCF and G4 (blue) and G4-only (orange)), sorted on the change of the first Eigenvector (EV1) signal from left to right flank. CTCF and RAD21 ChIP-seq, calculated insulation and first Eigenvector (EV1) in HAP1-CTCFdegron-TIR1 cells in the absence and presence of auxin (NT and IAA) were plotted along with the published K562 G4 ChIP-seq and HAP1 RNA polII ChIP-seq.

Figure S3. HAP1-RPB1-AID characterization and insulation for the three categories of boundaries in CTCF, RAD21 and RPB1 degrons, related to Figure 3.

(A) Stackups for active TTS (without CTCF, with R-loops), sorted on the change of first Eigenvector (EV1) signal from left to right flank. RAD21 ChIP-seq, calculated insulation, EV1 values and RNA-seq in HAP1-CTCFdegron-TIR1 cells in the absence and presence of auxin (NT and IAA) were plotted along with the published HAP1 RNA polII ChIP-seq, HAP1 H3K4me3 ChIP-seq, consensus list of R-loops and genes. Genes were flipped according to their orientations to have the body of the gene on the left of the TTS.

(B) Schematic of the HAP1-RPB1-AID construct (top) and western blot against RPB1 in WT HAP1 cells and in the HAP1-RPB1-AID cells showing RPB1 depletion after 4 hours of auxin treatment (IAA). Ponceau is shown for loading control. Flow cytometry for HAP1-RPB1-AID cells in absence and presence of auxin (NT and IAA) stained with Propidium Iodide (PI) to assess the DNA content for cell cycle analysis.

(C) Hi-C contact frequency as a function of genomic distance, P(s) (top) and its derivative dP/ds (bottom) for HAP1-RPB1-AID cells in absence and presence of auxin (NT and IAA).

(D) Hi-C contact heatmaps at 100kb resolution with the corresponding track of the first Eigenvector (EV1) across a 60Mb region on chromosome 2 for HAP1 cells and HAP1-RPB1-AID cells in absence and presence of auxin (NT and IAA). Genome-wide saddle plots of Hi-C data binned at 100kb resolution for HAP1 cells and HAP1-RPB1-AID cells in absence and presence of auxin (NT and IAA). The numbers in the corners of the saddle plots indicate compartment strengths for A and B compartments.

(E) RPB1 depletion weakens insulation at some boundaries. Hi-C contact heatmaps at 25kb resolution with the corresponding distribution of of the first Eigenvector values (EV1) and published RPB1 ChIP-seq signal for a 10Mb region on chromosome 14 for HAP1 cells and HAP1-RPB1-AID cells in absence and presence of auxin (NT and IAA). The differential interaction heatmap (presence of auxin / absence of auxin) is shown on the bottom. Dot pileups for dots found in HAP1 cells (Sanborn et al., 2015) that have a CTCF peak in either anchor in the CTCF degron NT sample (4 496 dots) for HAP1-RPB1-AID cells in absence and presence of auxin (NT and IAA). The dots were aggregated at the center of a 100kb window at 2kb resolution.

(F) Stackups for the three categories of insulation: the CTCF-only peaks (with RAD21) (blue), the active TSS common between HAP1 and HCT116 cells (without CTCF) (orange) and the active TTS common between HAP1 and HCT116 cells (without CTCF and H3K4me3) (green), sorted on the Non-Treated (NT) RAD21 ChIP-seq signal. The CTCF and RAD21 ChIP-seq signals, the RNA-seq and the insulation in the HAP1-CTCFdegron-TIR1, the insulation in the HAP1-RPB1-AID in the absence and presence of auxin (NT and IAA) were plotted along with the published K562 R-loops, HAP1 RNA polII ChIP-seq and the HAP1 H3K4me3 ChIP-seq signals. The insulation in the HCT116-RAD21-AID cells together with the RAD21 ChIP-seq in the HCT116-RAD21-AID cells were also plotted in absence and presence of auxin (NT and IAA). The active TSS common between HAP1 and HCT116 cells (without CTCF) were plotted underneath with a different scale to show the remaining RAD21 after RAD21 depletion (red dashed rectangle).

(G) Average insulation profiles across CTCF-only peaks (with RAD21) (left plot), across scaled active genes (without CTCF at TSS and TTS) (middle plot) and across scaled inactive genes (right plot) for the HCT116-RAD21-AID cells in absence and presence of auxin (NT and IAA).

Figure S4. Genome wide CRISPR screen after CTCF depletion, related to Figure 4

(A) Western blot against CTCF in HAP1-CTCFdegron and HAP1-CTCFdegron-TIR1 cells in absence and presence of auxin (NT and IAA) with the two different auxin concentrations (25μM and 500μM) used in the screen showing the partial CTCF depletion with 25μM IAA and the nearly total CTCF depletion with 500μM IAA. Ponceau is shown for loading control.

(B) Plot showing the cumulative number of doublings relative to the days cells were passaged during the screen for HAP1-CTCFdegron and HAP1-CTCFdegron-TIR1 for the three auxin concentrations (NT, 25μM and 500μM).

(C) Fold change distribution of essential and nonessential gene sets at indicated time points for the screens for HAP1-CTCFdegron and HAP1-CTCFdegron-TIR1 cells for the three auxin concentrations (NT, 25μM and 500μM) (left). Precision-recall curves based on Bayes Factors (BFs) of predefined essential and non-essential gene sets for the screens for HAP1-CTCFdegron and HAP1-CTCFdegron-TIR1 cells for the three auxin concentrations (NT, 25μM and 500μM) at the indicated time points (right).

(D) Scatter plots of the log2FC for HAP1-CTCFdegron-TIR1 screens against HAP1-CTCFdegron screens for T6 and T15 time points. Genes with a fold change of ≥ 2 between HAP1-CTCFdegron and HAP1-CTCFdegron-TIR1 screens are highlighted in red. Genes linked to CTCF are indicated (cohesin, TOP2A) along with the two studied gene hits (DDX55 and TAF5L).

(E) Scatter plots of the log2FC for HAP1-CTCFdegron cells in absence and presence of auxin (NT and IAA) at T6 and T15 time points. Genes with a fold change of ≥ 2 between absence and presence of auxin (NT and IAA) are highlighted in red and considered as auxin specific genes. Genes linked to CTCF are indicated (cohesin genes, TOP2A) along with the two studied gene hits (DDX55 and TAF5L).

Figure S5. DDX55, TAF5L and TAF6L interact with CTCF and cohesin. Chromatin binding of DDX55 and TAF5L is reduced after CTCF depletion, related to Figure 5.

(A) Western blot co-IP replicates against DDX55 and TAF5L in HAP1-CTCFdegron-TIR1 cells in absence and presence of auxin (NT and IAA), treated with either turbonuclease (DNA - and RNA -) or RNAseA (RNA -).

(B) Western blot co-IP replicates against DDX55 and TAF5L in HCT116-RAD21-AID cells in absence and presence of auxin (NT and IAA), treated with either turbonuclease (DNA - and RNA -) or RNAseA (RNA -).

(C) Western blot co-IP (two replicates) against TAF6L in HAP1-CTCFdegron-TIR1 cells in absence and presence of auxin (NT and IAA), treated with either turbonuclease (DNA - and RNA -) or RNAseA (RNA -).

(D) Representative western blots against CTCF, RAD21, DDX55 and β-ACTIN (loading control) to check the CTCF depletion efficiency in the HAP1-CTCFdegron-TIR1 co-IP (left) and the RAD21 depletion efficiency in the HCT116-RAD21-AID cells co-IP (right).

(E) Efficient DNA digestion by turbonuclease (TURBO) during DDX55, TAF5L and TAF6L co-IPs. DNA digestion was assessed by qPCR using primers specific to ACTB and POLR2A gene locations. Error bars are SD, n = 20.

(F) Efficient RNA digestion by turbonuclease (TURBO) and RNaseA (RNASEA) during DDX55, TAF5L and TAF6L co-IPs. RNA digestion was assessed by qPCR using primers specific to ACTB and POLR2A gene locations. Error bars are SD, n = 20.

(G) Stackups for CTCF-only sites with RAD21, sorted on the Non-Treated (NT) DDX55 ChIP-seq signal. CTCF ChIP-seq, RAD21 ChIP-seq, calculated insulation, DDX55 ChIP-seq, TAF5L ChIP-seq and RNA seq signals in HAP1-CTCFdegron-TIR1 cells in absence and presence of auxin (NT and IAA) were plotted along with published HAP1 H3K4me3 ChIP-seq and genes. Genes on the forward strand are represented in red (plus) and genes on the reverse strand are represented in blue (minus), grey color corresponds to loci without annotated transcripts. For TTS, stackups were flipped according to the orientation of the genes, to have the body of the gene on the left of the TTS.

(H) Stackup quantification for the CTCF-only sites (with RAD21) for DDX55 and TAF5L ChIP-seq for both replicates: a distribution of ratios between auxin-treated and non-treated signals. A fold change < 1 represents less binding of DDX55 or TAF5L at CTCF-only sites (with RAD21) after CTCF depletion.

Figure S6. Characterization of DDX55 and TAF5L depletion in HAP1-CTCFdegron-TIR1 cells, related to Figure 6.

(A) Western blots against CTCF, DDX55, TAF5L and β-ACTIN (loading control) showing DDX55 and TAF5L depletions, by siRNA and mutations in DDX55 and TAF5L genes, compared to siRNA controls and mutations at the AAVS1 non-coding sequence (CTRL clone).

(B) Flow cytometry for HAP1-CTCFdegron-TIR1 cells in absence and presence of auxin (NT and IAA) with siRNA (siCTRL (control), siDDX55 and siTAF5L) (left) and CTRL (control), DDX55 and TAF5L knock-out clones (right) stained with Propidium Iodide (PI) to assess the DNA content for cell cycle analysis.

(C) Hi-C contact frequency as a function of genomic distance P(s) and its derivative dP/ds for HAP1-CTCFdegron-TIR1 cells in absence and presence of auxin (NT and IAA) treated with siRNA (siCTRL, siDDX55 and siTAF5L) (top) and CTRL, DDX55 and TAF5L knock-out clones (bottom).

(D) Genome-wide saddle plots of Hi-C data binned at 100kb resolution for HAP1-CTCFdegron-TIR1 cells in absence and presence of auxin (NT and IAA) treated with siRNA (siCTRL, siDDX55 and siTAF5L) (top) and CTRL, DDX55 and TAF5L knock-out clones (bottom). The numbers in the corners of the saddle plots indicate compartment strengths for A and B compartments.

(E) RNA-seq expression (TPM) for HAP1-CTCFdegron-TIR1 cells in absence and presence of auxin (NT and IAA) treated with siRNA (siCTRL, siDDX55 and siTAF5L) and CTRL, DDX55 and TAF5L knock-out clones for key genes (DDX55, TAF5L, CTCF, RAD21, SMC1A, SMC3, WAPAL, NIPBL, STAG1 and STAG2).

Figure S7. Effects on enhancer-promoter interactions and gene expression, related to Figure 7.

(A) Aggregated and scaled Hi-C maps for enhancer (without CTCF or TSS)-promoter (without CTCF) pairs with or without intervening CTCF peaks for HAP1-CTCFdegron-TIR1 cells in absence and presence of auxin (NT and IAA) treated with siRNA (siCTRL, siDDX55 and siTAF5L) and CTRL, DDX55 and TAF5L knock-out clones.

(B) Number of Differentially Expressed genes (left) and alternatively spliced genes (right) in HAP1-CTCFdegron-TIR1 cells in absence and presence of auxin (NT and IAA) treated with siRNA (siCTRL, siDDX55 and siTAF5L) and CTRL, DDX55 and TAF5L knock-out clones. Grey bars indicate the CTCF depletion, blue bars indicate the siRNA depletion and red bars indicate the knock-out clones.

## ACKNOWLEDGMENTS

We thank members of the Dekker, the Mirny laboratories and the members of Open Chromosome Collective for creating a collaborative atmosphere and insightful discussions. We thank the UMass Flow Cytometry Core Facility for FACS sorting the cell lines. We thank Masato Kanemaki for sharing the HCT116-RAD21-AID cell line. We thank Elphege Nora and Benoit Bruneau for sharing CTCF-degron plasmids. This work was supported by a grant from the National Human Genome Research Institute (NHGRI) to J.D. (HG003143), and a grant from National Institute of General Medical Sciences (NIGMS) to A.A.P (GM133762). J.D. is an investigator of the Howard Hughes Medical Institute. Some of the schematic figures were created with BioRender.com.

