## Supplementary figures and images for "A cohesin traffic pattern genetically linked to gene regulation"

### Supplemental Figures

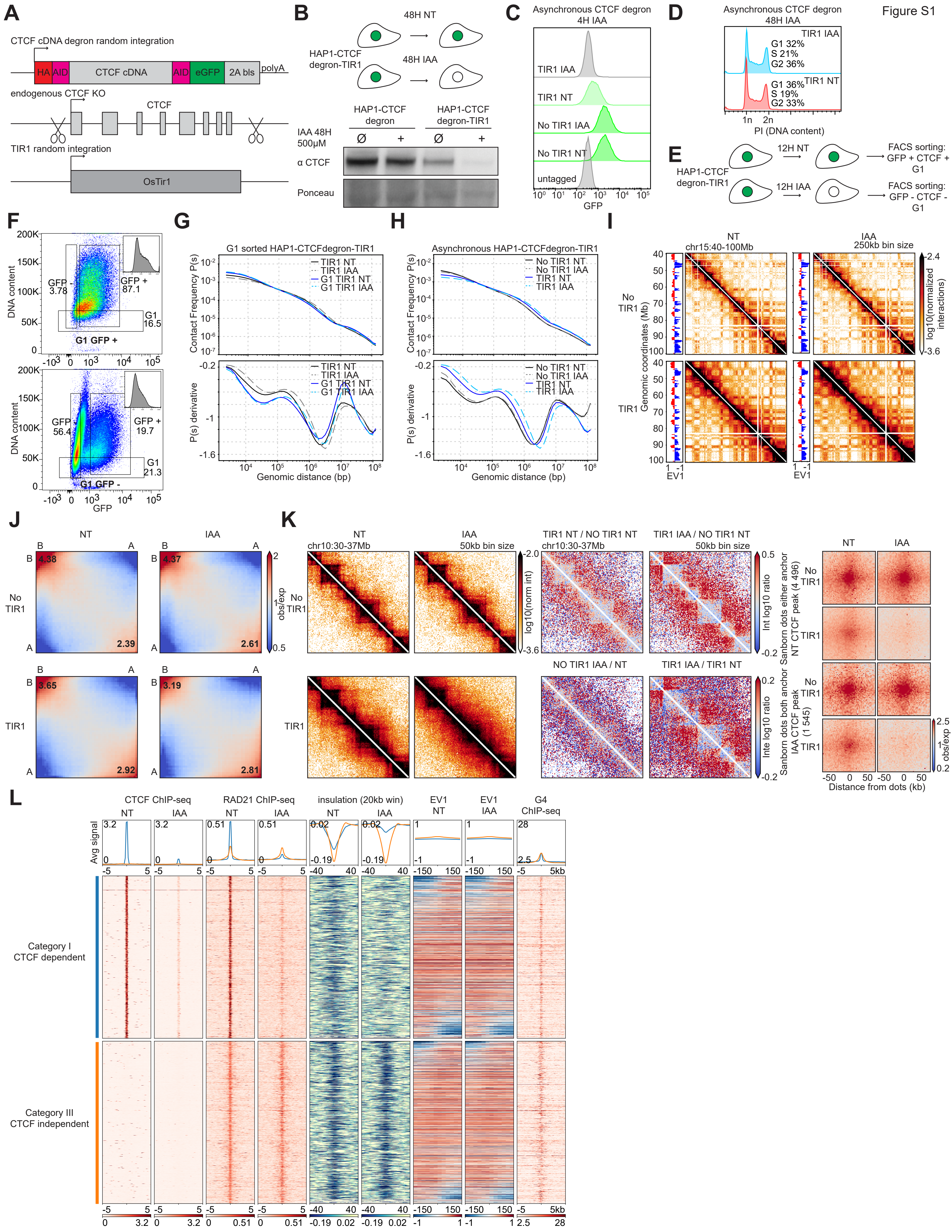

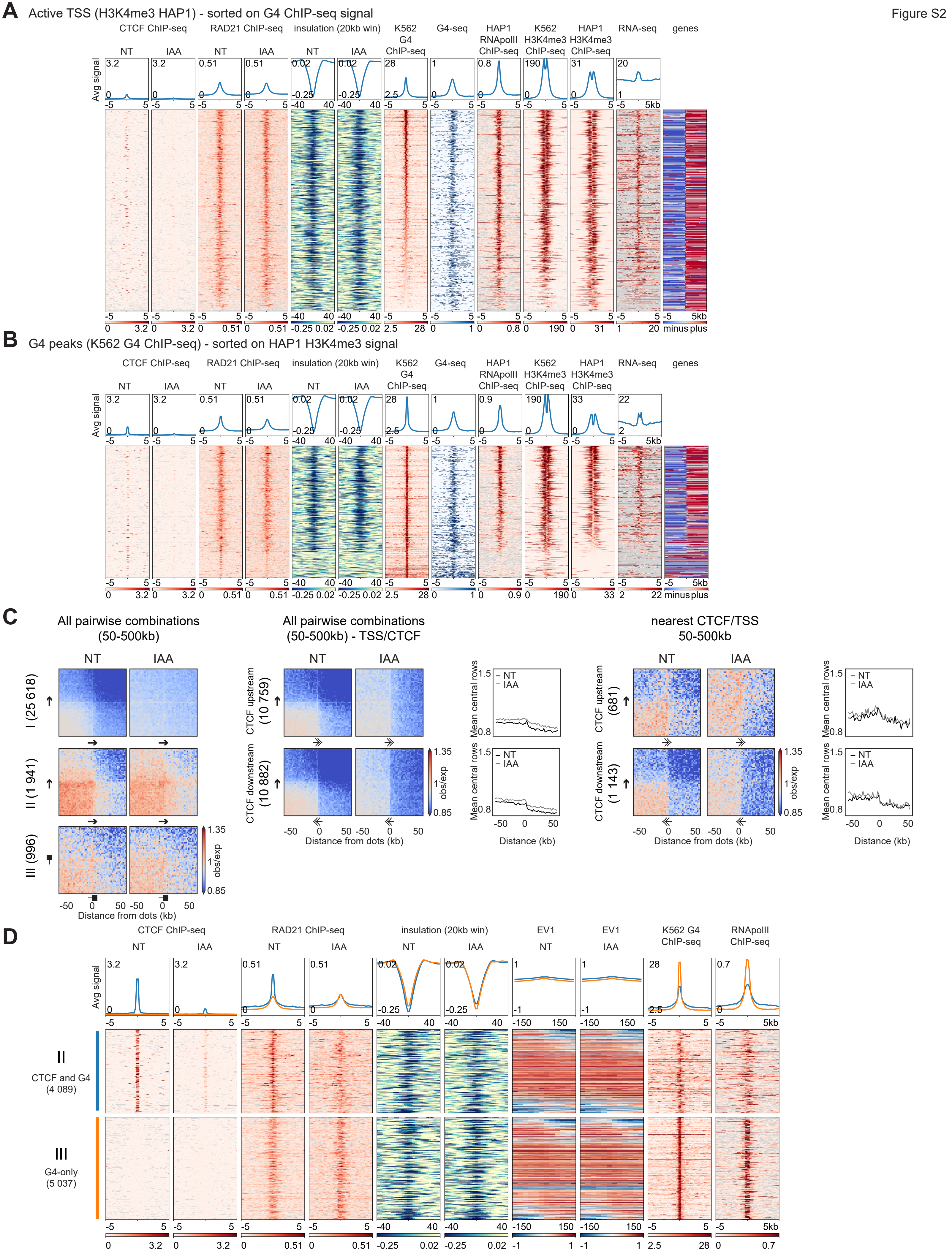

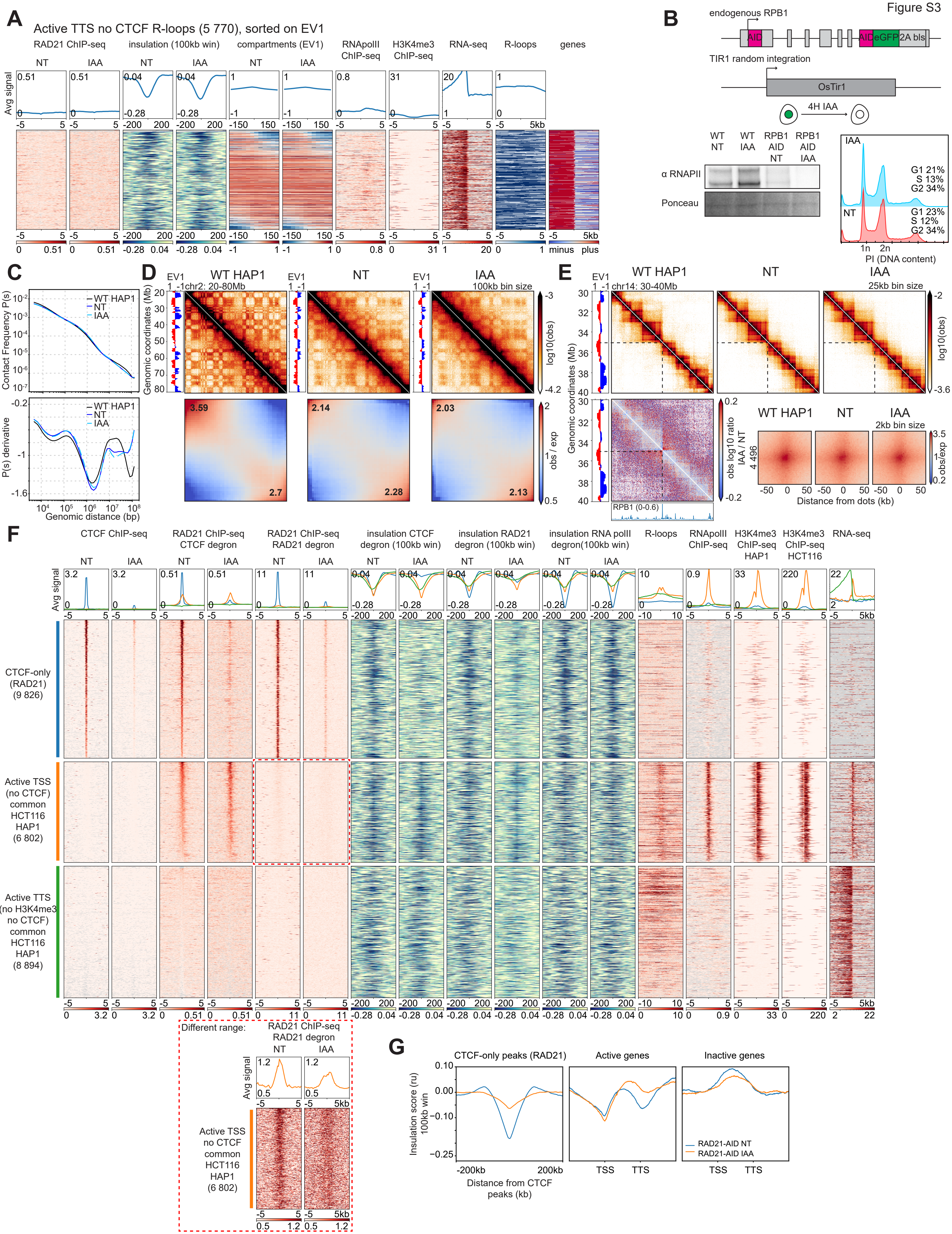

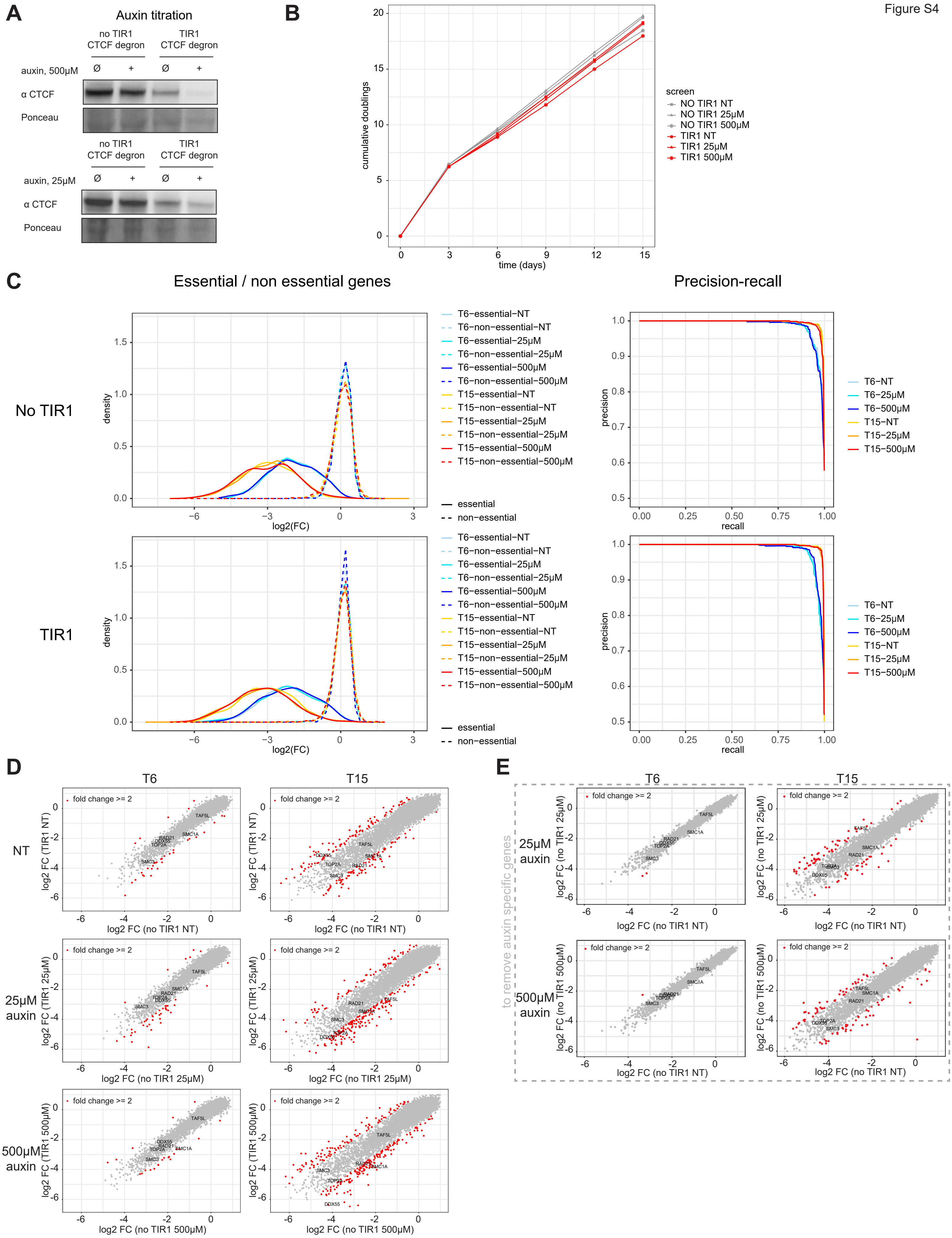

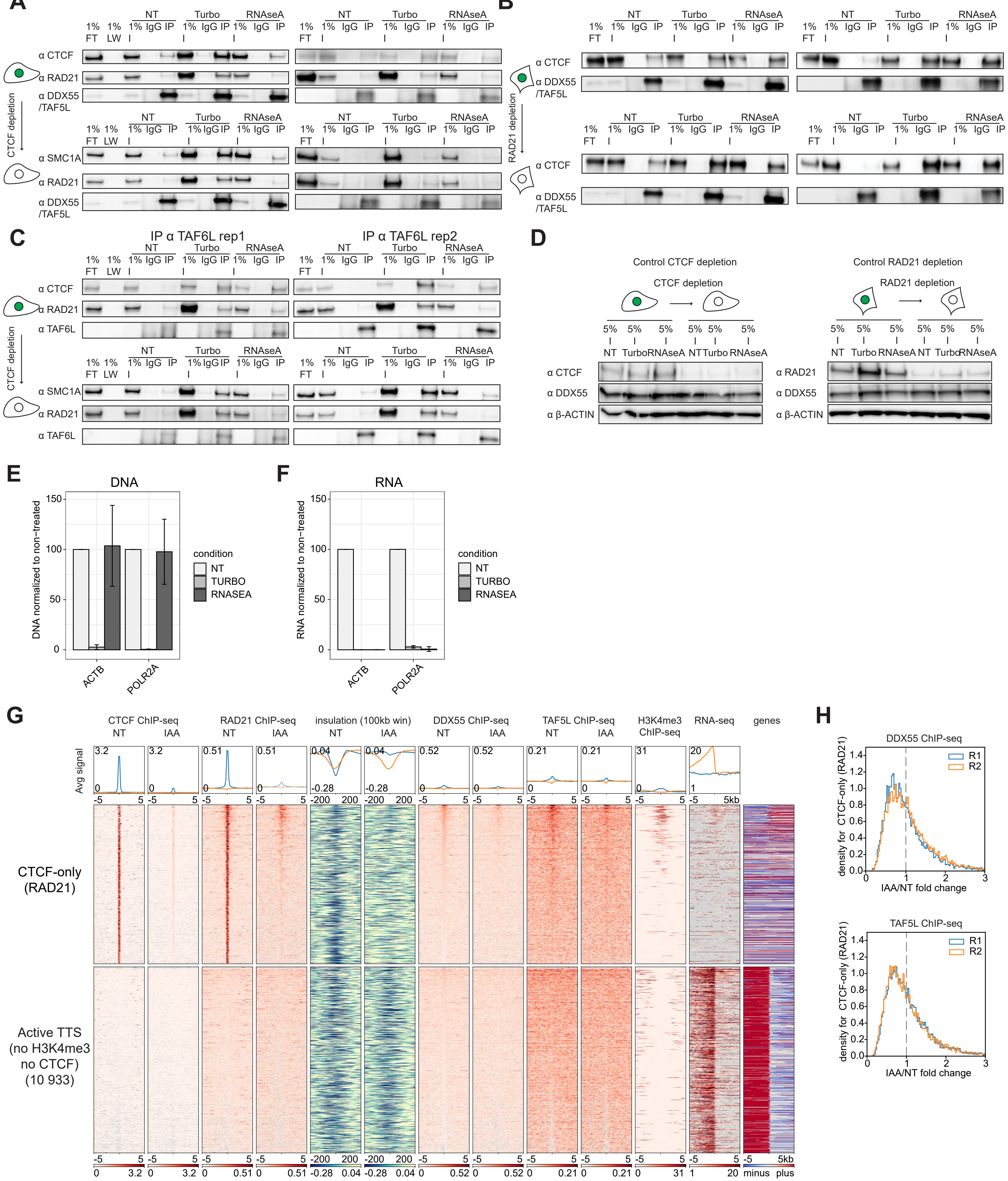

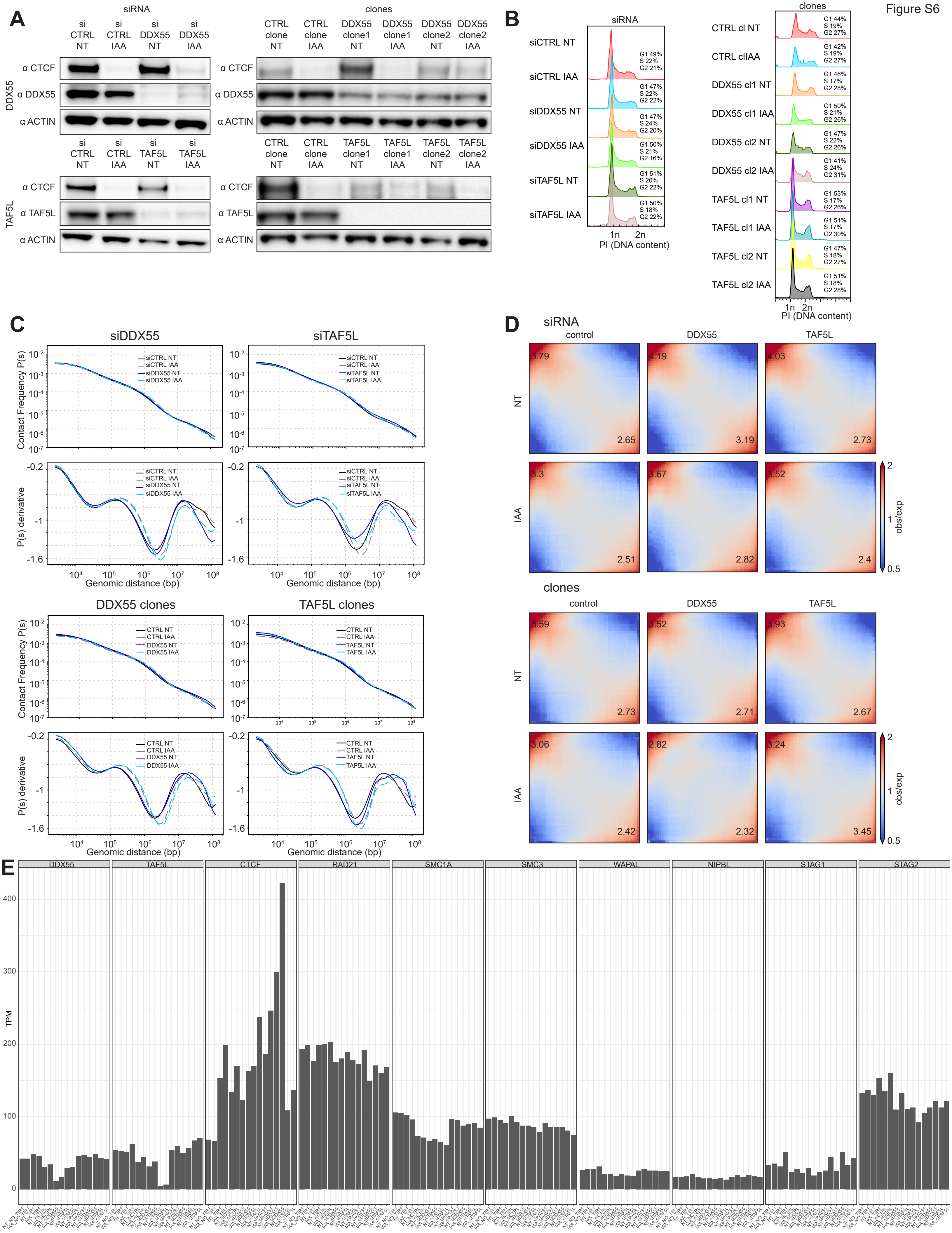

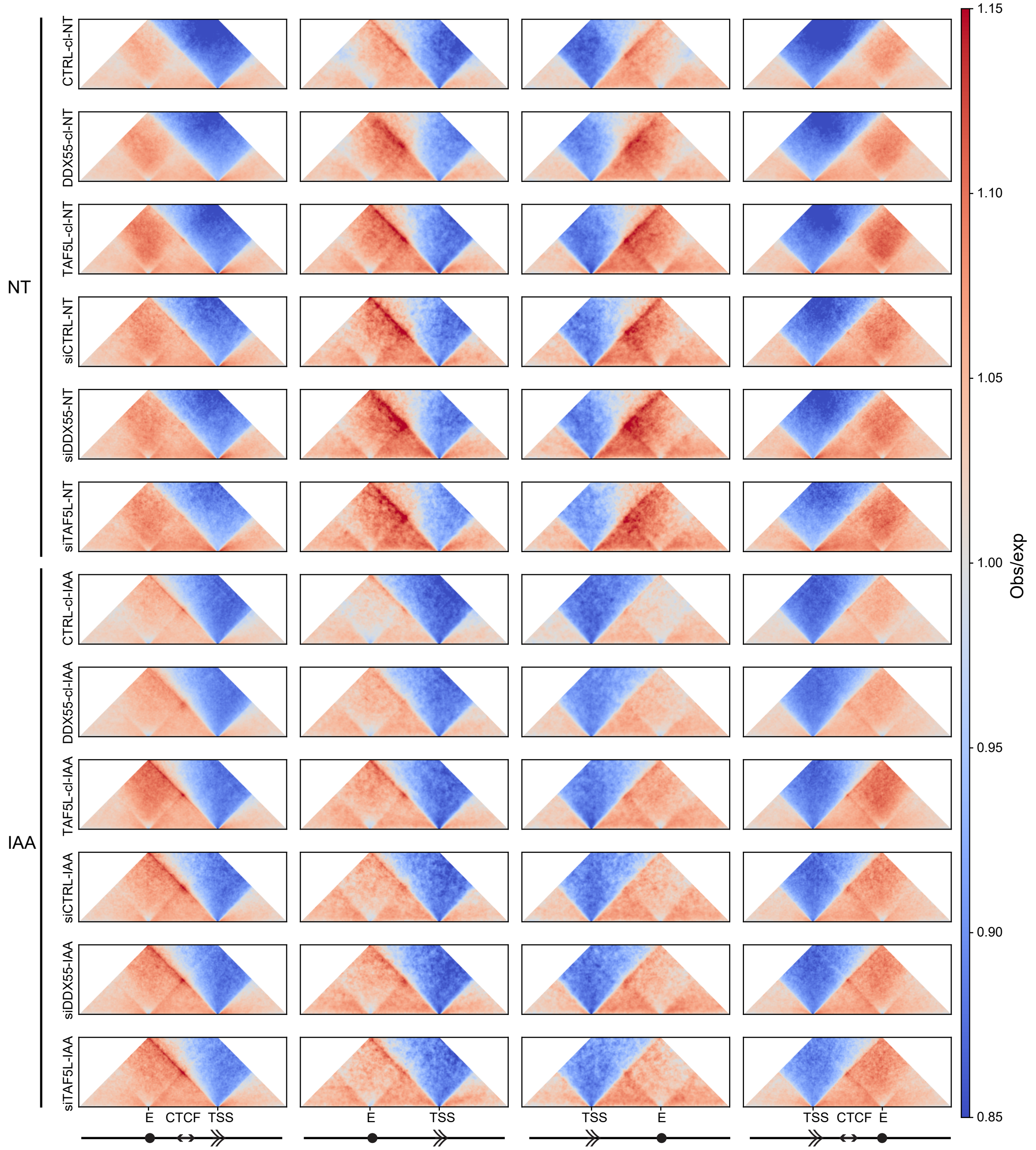

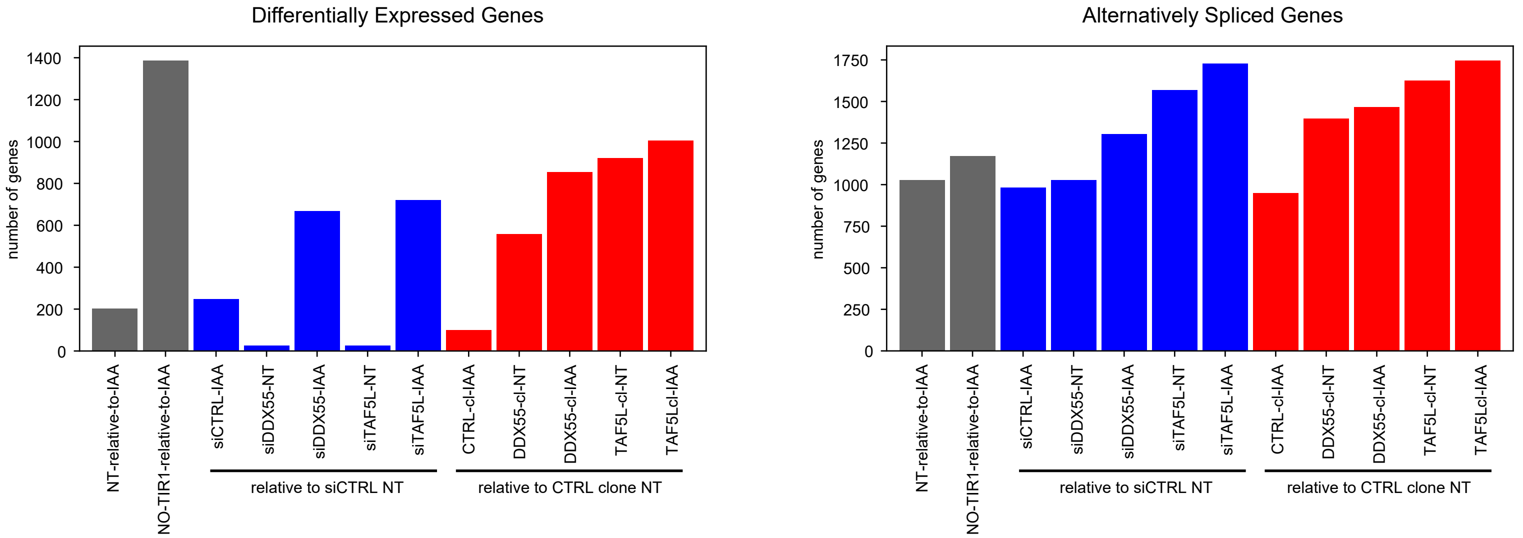
